# Accommodating individual travel history, global mobility, and unsampled diversity in phylogeography: a SARS-CoV-2 case study

**DOI:** 10.1101/2020.06.22.165464

**Authors:** Philippe Lemey, Samuel Hong, Verity Hill, Guy Baele, Chiara Poletto, Vittoria Colizza, Áine O’Toole, John T. McCrone, Kristian G. Andersen, Michael Worobey, Martha I. Nelson, Andrew Rambaut, Marc A. Suchard

**Affiliations:** KU Leuven Department of Microbiology, Immunology and Transplantation, Rega Institute, Laboratory of Clinical and Evolutionary Virology, Leuven, Belgium; Centre for Immunology, Infection and Evolution, University of Edinburgh, King’s Buildings, Edinburgh, EH9 3FL, UK; INSERM, Sorbonne Université, Institut Pierre Louis d’Epidémiologie et de Santé Publique IPLESP, F75012 Paris, France; Department of Immunology and Microbiology, Scripps Research, La Jolla, CA 92037, USA; Department of Ecology and Evolutionary Biology, University of Arizona, Tucson, AZ 85721, USA; Division of International Epidemiology and Population Studies, Fogarty International Center, National Institutes of Health, Bethesda, Maryland 20892, USA; Department of Biomathematics, David Geffen School of Medicine, University of California Los Angeles, Los Angeles, CA 90095, USA; Department of Biostatistics, Fielding School of Public Health, University of California Los Angeles, Los Angeles, CA 90095, USA; Department of Human Genetics, David Geffen School of Medicine, University of California Los Angeles, Los Angeles, CA 90095, USA

## Abstract

Spatiotemporal bias in genome sequence sampling can severely confound phylogeographic inference based on discrete trait ancestral reconstruction. This has impeded our ability to accurately track the emergence and spread of SARS-CoV-2, the virus responsible for the COVID-19 pandemic. Despite the availability of unprecedented numbers of SARS-CoV-2 genomes on a global scale, evolutionary reconstructions are hindered by the slow accumulation of sequence divergence over its relatively short transmission history. When confronted with these issues, incorporating additional contextual data may critically inform phylodynamic reconstructions. Here, we present a new approach to integrate individual travel history data in Bayesian phylogeographic inference and apply it to the early spread of SARS-CoV-2, while also including global air transportation data. We demonstrate that including travel history data for each SARS-CoV-2 genome yields more realistic reconstructions of virus spread, particularly when travelers from undersampled locations are included to mitigate sampling bias. We further explore methods to ameliorate the impact of sampling bias by augmenting the phylogeographic analysis with lineages from undersampled locations in the analyses. Our reconstructions reinforce specific transmission hypotheses suggested by the inclusion of travel history data, but also suggest alternative routes of virus migration that are plausible within the epidemiological context but are not apparent with current sampling efforts. Although further research is needed to fully examine the performance of our travel-aware phylogeographic analyses with unsampled diversity and to further improve them, they represent multiple new avenues for directly addressing the colossal issue of sample bias in phylogeographic inference.

## Introduction

Since its emergence in late 2019, SARS-CoV-2 has rapidly spread across the world, prompting governments to enact restrictions on human mobility that are unprecedented on a global scale. While Coronavirus Disease 2019 (COVID-19) has exposed critical gaps in public health preparedness and research, an observed strength of the COVID-19 response has been the rapid speed with which whole-genome sequences of SARS-CoV-2 have been generated globally (>46,000 genomes as of June 13, 2020). The success of global sequence production has at least been partially facilitated by protocols and networks that arose during the response to the 2013-2016 Ebola virus disease (Ebola) epidemic in West Africa. That Ebola epidemic was particularly important for spurring development of tools and methods for real-time in-country virus sequencing, including in resource-limited settings ^e.g. 1^.

Genomic data represent a key resource for testing hypotheses about how and when SARS-CoV-2 became established in different locations. For example, phylogenetic approaches may be able to distinguish community transmission from new introductions from travelers ^2^, whether viral outbreaks were associated with multiple introductions ^3^, how long viruses may have been transmitting undetected in a community (‘cryptic transmission’ ^4^) and when widespread domestic spread began in the United States ^5^. These studies have already greatly contributed to charting the course of an unfolding pandemic, informing public health decisions about when to enforce lockdown measures of various degrees of stringency. Recently, many countries have entered a new phase in which restrictions such as school and business closures and mobility restrictions are eased or lifted. Suppressing SARS-CoV-2 spread remains our only viable defense so far and efficient test, trace and isolate systems will be crucial tools to achieve these goals. Molecular epidemiology can continue to inform public health actions during this phase, for instance through uncovering cryptic transmission or new introductions as sources of flare-ups.

Recent advances in virus sequencing and phylogenetics hold great promise for addressing key questions in infectious disease epidemiology and outbreak response ^6^. There are limitations, however, to the insights that can be obtained from the wealth of SARS-CoV-2 genome data. The current sequence diversity is relatively limited because SARS-CoV-2 emerged only recently in late 2019 and because SARS-CoV-2 transmission outpaces the rate at which substitutions accumulate ^4^. This implies that short-term transmission patterns may not leave a detectable footprint in virus genomes. In addition, large spatiotemporal biases exist in the available genome data. For instance, about 46% of currently available genomes have been sampled from the UK whereas Italy, having experienced a similar number of cases and likely an earlier epidemic onset, only represents 0.3% of the genome collection on GISAID. Both low sequence diversity and sampling bias confound the interpretation of transmission patterns, and highly similar SARS-CoV-2 genomes from the same or different locations do not necessarily imply direct linkage.

Despite a relatively slow evolutionary rate, the ‘phylodynamic threshold’ for SARS-CoV-2 was reached relatively early ^7^, meaning that sufficient divergence had accumulated over the sampling time range to infer time-calibrated phylogenies and the underlying transmission processes that generate such trees, including spatiotemporal spread. However, sampling bias presents a critical challenge for popular discrete trait ancestral reconstruction procedures ^8,9^. Although the modern phylodynamic framework includes other statistical approaches that are less sensitive to sampling bias ^e.g. 10–12^, computational complexity challenges their application to large data sets, in particular when insights are needed in short turnaround times. This explains the widespread adoption of ancestral reconstruction approaches that provide real-time tracking of pathogen evolution and spread ^13^.

When confronted with low diversity and sampling bias, evolutionary reconstructions may greatly benefit from integrating additional sources of information. Bayesian phylodynamic approaches are particularly adept for this purpose ^14^ and phylogeographic methods in particular have been extended to take advantage of human transportation data as proxies of population-level connectivity between locations ^15^. This approach has been utilized in a wide range of applications, including the identification of the key drivers of *Ebolavirus* spread in West Africa ^16^. Individual travel history of sampled patients also represents an important source of information that currently has not been used to its full potential in phylogeographic inference. Genomic data from (returning) travellers may help to uncover pathogen diversity in locations that are otherwise undersampled. This has been elegantly demonstrated by a study on Zika virus that used travel surveillance and genomics to demonstrate hidden viral transmission in Cuba ^17^. Formally integrating such travel data in phylogeographic reconstructions may therefore help to address or correct for sampling bias. In general, epidemiological information provides important context to assess genomic sampling biases, and this can be used to sub-sample genomes by location in situations where large collections are available ^18^. The question therefore emerges how such information can be formally embedded in phylodynamic models. Specifically, if particular locations remain undersampled despite their potential importance for viral spread, can the reconstructions account for hypotheses alternative to the ones supported by the sampled genomes but plausible according to the epidemiological context? Here, we extend phylogeographic methodology to incorporate travel data and, together with the integration of transportation data, we apply this to reconstruct the early spread of SARS-CoV-2. In addition, we demonstrate how epidemiological data can be used to incorporate unsampled diversity. Taken together, these approaches constitute an important step towards more realistic and more nuanced phylogeographic reconstructions.

## Materials and Methods

### SARS-CoV-2 Genome Data Set and associated travel history

To focus on the early stage of COVID-19 spread, we analyzed SARS-CoV-2 genome sequences and metadata available in GISAID on March 10th ^19^. We curated a data set of 305 genomes by removing error-prone sequences, keeping only a single genome from patients with multiple genomes available, and removing genomes obtained from people infected on the Princess Diamond Cruise ship (which we cannot include as a location in our phylogeographic analysis). We assigned each genome a global lineage designation based on the nomenclature scheme outlined in Rambaut *et al* ^20^ using pangolin v1.1.14 (https://github.com/hCoV-2019/pangolin), lineages data release 2020-05-19 (https://github.com/hCoV-2019/lineages). We aligned the remaining 286 genomes using MAFFT ^21^ and partially trimmed the 5’ and 3’ ends. All sequences were associated with exact sampling dates in their meta-information, except for 18 genomes with known months of sampling (one from Anhui, China and 17 from Shandong, China). Upon visualizing root-to-tip divergence as a function of sampling time using TempEst ^22^ based on an ML tree inferred with IQ-TREE ^23^, we removed one potential outlier. We formally tested for temporal signal using BETS ^24^. The final 284 genomes were sampled from 28 different countries, with Chinese samples originating from 13 provinces, one municipality (Beijing) and one special administrative area (Hong Kong), which we considered as separate locations in our (discrete) phylogeographic analyses.

We searched for travel history data associated with the genomes in the GISAID records, media reports and publications and retrieved recent travel locations for 64 genomes (22.5%, Supplementary Table 1): 43 travelled/returned from Hubei (Wuhan), 1 from Beijing, 3 from China without further detail (which we associated with an appropriate ambiguity code in our phylogeographic analysis that represents all sampled Chinese locations), 2 from Singapore, 1 from Southeast Asia (which we also associated with an ambiguity code that represents all sampled Southeast Asian locations), 7 from Italy and 7 from Iran. In this dataset, Italy is better represented by recent travel locations than actual samples (*n* = 4) and Iran is exclusively represented by travellers returning from this country. For 46 of the 64 genomes, we retrieved the date of travel, which represents the most recent time point at which the ancestral lineage circulated in the travel location.

**Table 1:** Inferred discrete location generalized linear model (GLM) of transmission under three different usage strategies of travel history information. We report posterior inclusion probabilities and posterior mean (95% highest probability density intervals) log conditional effect sizes for air travel, geographic distance and asymmetry out of Hubei.

| Information usage strategy | Air travel |  | Geographic distance |  | Asymmetry out of Hubei |  |
| --- | --- | --- | --- | --- | --- | --- |
|  | Inclusion probability | Log conditional effect size | Inclusion probability | Log conditional effect size | Inclusion probability | Log conditional effect size |
| Sampling location and travel history | >.999 | 0.852<br>(0.629,1.106) | 0.105 | -0.044<br>(-0.111,0.031) | >.999 | 2.396<br>(1.958,2.859) |
| Sampling location only | >.999 | 1.140<br>(0.850,1.468) | 0.065 | -0.016<br>(-0.096,0.057) | >.999 | 1.829<br>(1.167,2.417) |
| Travel origin location | >.999 | 0.975<br>(0.650,1.289) | 0.125 | 0.060<br>(-0.031,0.148) | >.999 | 2.334<br>(1.788,2.924) |

### Incorporating travel history in Bayesian phylogeographic inference

Discrete trait phylogeographic inference attempts to reconstruct an ancestral location-transition history along a phylogeny based on the discrete states associated with the sampled sequences. In our Bayesian approach, the phylogeny is treated as random which is critical to accommodate estimation uncertainty when confronted with sparse sequence information. Here, we aim to augment these location-transition history reconstructions on random trees with travel history information obtained from (returning) travellers. When such information is available, the tip location state for a sequence can either be set to the location of sampling, as is done in the absence of such information, or the location from which the individual travelled (assuming that this was the location from which the infection was acquired). Neither of these options is satisfactory: using the location of sampling ignores important information about the ancestral location of the sequence, whereas using the travel location together with the collection date represents a data mismatch and ignores the final transitions to the location of sampling. These events are particularly important when the infected traveller then produces a productive transmission chain in the sampling location.

Incorporating information about ancestral locations cannot be achieved simply through the parameterization of the discrete diffusion model, which follows a continuous-time Markov chain (CTMC) process determined by relative transition rates between all pairs of locations that applies homogeneously (or time inhomogeneously ^25^) along the phylogeny. Instead, we need to shape the realization of this process according to the travel histories by augmenting the phylogeny with ancestral nodes that are associated with a location state (but not with a known sequence) and hence enforce that ancestral location at a particular, possibly random point in the past of a lineage. Further, the location state can also be ambiguous, allowing equal or weighted probability to be assigned to multiple possible locations ^26^. We illustrate this procedure for an empirical example that includes 9 SARS-CoV-2 genomes in Fig. 1.

**Figure 1:**
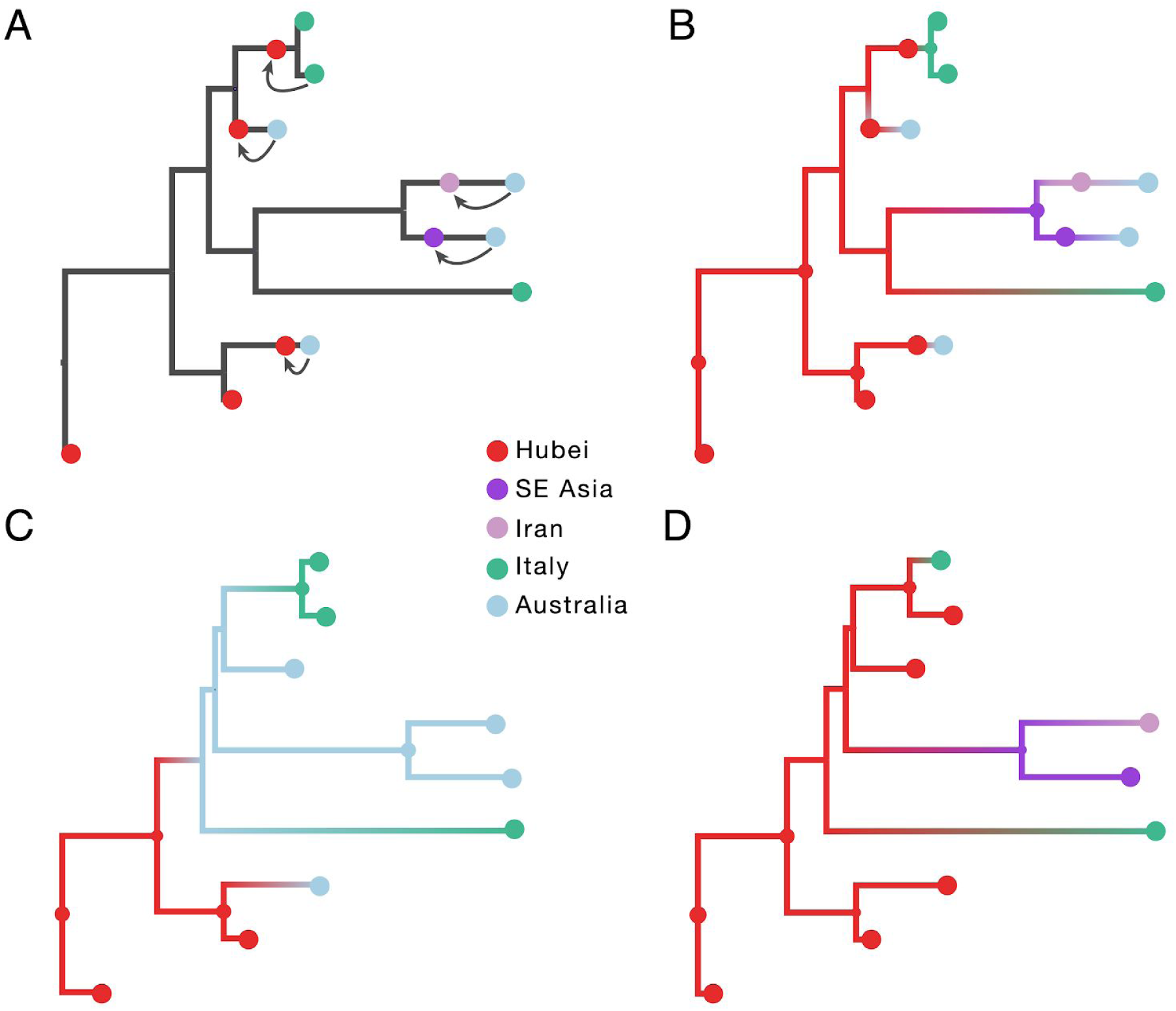
Incorporating travel history data in phylogeographic reconstruction. Panel A illustrates the concept of introducing ancestral nodes associated with locations from which travellers returned. The ancestral nodes are indicated by arrows for five cases relating them to the genomes sampled from the travellers. The ancestral nodes are introduced at different times in the ancestral path of each sampled genome. Panels B, C and D represent the results from analyses using sampling location and travel history, sampling location only, and travel origin location respectively. The branch color reflects the modal state estimate at the child node. There is some topological variability but only involving nodes that are poorly supported.

The empirical example includes two genomes from Hubei, four from Australia, and three from Italy. Travel history is available for five genomes (one sampled in Italy and four in Australia) and Fig. 1A demonstrates how this information is incorporated. When a sampled traveller returned from location *i* to location *j*, we denote time *T*_*i*→*j*_ as the time when the traveller started the return journey to *j*. At this time point in the ancestral path of the tip (indicated with arrows for the 5 relevant tips), we introduce an ancestral node and associate it with location *i* in order to inform the reconstruction that at this point in time the lineage was in location *i*. The upper arrow represents the information introduced for the traveller that returned from Hubei to Italy. The same procedure is applied to the four genomes from travellers returning to Australian from Hubei, Iran, Southeast Asia and Hubei again (from top to bottom in Fig. 1A).

In subsequent panels, we compare a travel-aware reconstruction (B) to a reconstruction using the standard sampling location (C) and a reconstruction using the location of origin for the travellers (D). Using the location of sampling (C) results in an unrealistic Australian ancestry and two transitions from Australia to Italy, likely because Australia is represented by the largest number of tips. Using the location of travel origin (D) results in a reconstruction that better matches the travel-aware reconstruction in terms of inferring an ancestry in Hubei, but misses transitions along four tip branches and differs from the reconstruction including travel history for the upper two Italian genomes. Specifically it implies a transition from Hubei for the Italian patient that does not have travel history.

We note that *T*_*i*→*j*_ can be treated as a random variable in case the time of travelling to the sampling location is not known (with sufficient precision). We make use of this ability for the genomes associated with a travel location but without a clear travel time. In our Bayesian inference, we specify normal prior distributions over *T*_*i*→*j*_ informed by an estimate of time of infection and truncated to be positive (back-in-time) relative to sampling date. Specifically, we use a mean of 10 days before sampling based on a mean incubation time of 5 days ^27^ and a constant ascertainment period of 5 days between symptom onset and testing ^28^, and a standard deviation of 3 days to incorporate the uncertainty on the incubation time. Finally, we indicate that not only information about the sampled traveller can be incorporated, but also about prior transmission history. We apply this for two cases in our data set. One of the genomes was sampled from a German patient who was infected after contact with someone who came from Shanghai. The person travelling from Shanghai was assumed to be infected after being visited by her parents from Wuhan a few days before she left. In this case, we incorporate Wuhan (Hubei) as an ancestral location with an associated time that accounts for the travel time from Shanghai with a number of additional days and associated uncertainty. Another genome was obtained from a French person who had been in contact with a person who is believed to have contracted the virus at a conference in Singapore ^29^. In this case, we incorporate Singapore as an ancestral location with a known travel time (Supplementary Table 1).

### Incorporating unsampled diversity in Bayesian phylogeographic inference

To investigate how unsampled diversity may impact phylogeographic reconstructions, we include in our Bayesian inference taxa that are associated with a location but not with observed sequence data. We identify undersampled locations by considering the ratio of available genomes to the cumulative number of cases for each location (obtained from Our World in Data, https://ourworldindata.org/coronavirus-source-data). To keep all available data, we opt not to downsample genomes, but to add a number of unsampled taxa to specific locations in order to achieve a minimal ratio of taxa (sampled and unsampled) to cumulative number of cases. In Fig. 2A, we plot the number of available genomes against the number of cases on March 10th 2020 on a log-log scale. In our case, we set the minimal ratio arbitrarily to 0.005. Although higher ratios may be preferred, this comes at the expense of adding larger numbers of unsampled taxa and hence computationally more expensive Bayesian analyses. Our choice for the minimal ratio requires adding 458 taxa for 14 locations (colored symbols in Fig. 2), so about 1.6 times the number of available genomes. Most of the unsampled taxa are assigned to Hubei (*n* = 307), followed by Italy (*n* = 47), Iran (*n* = 40), and South Korea (*n* = 30).

**Figure 2:**
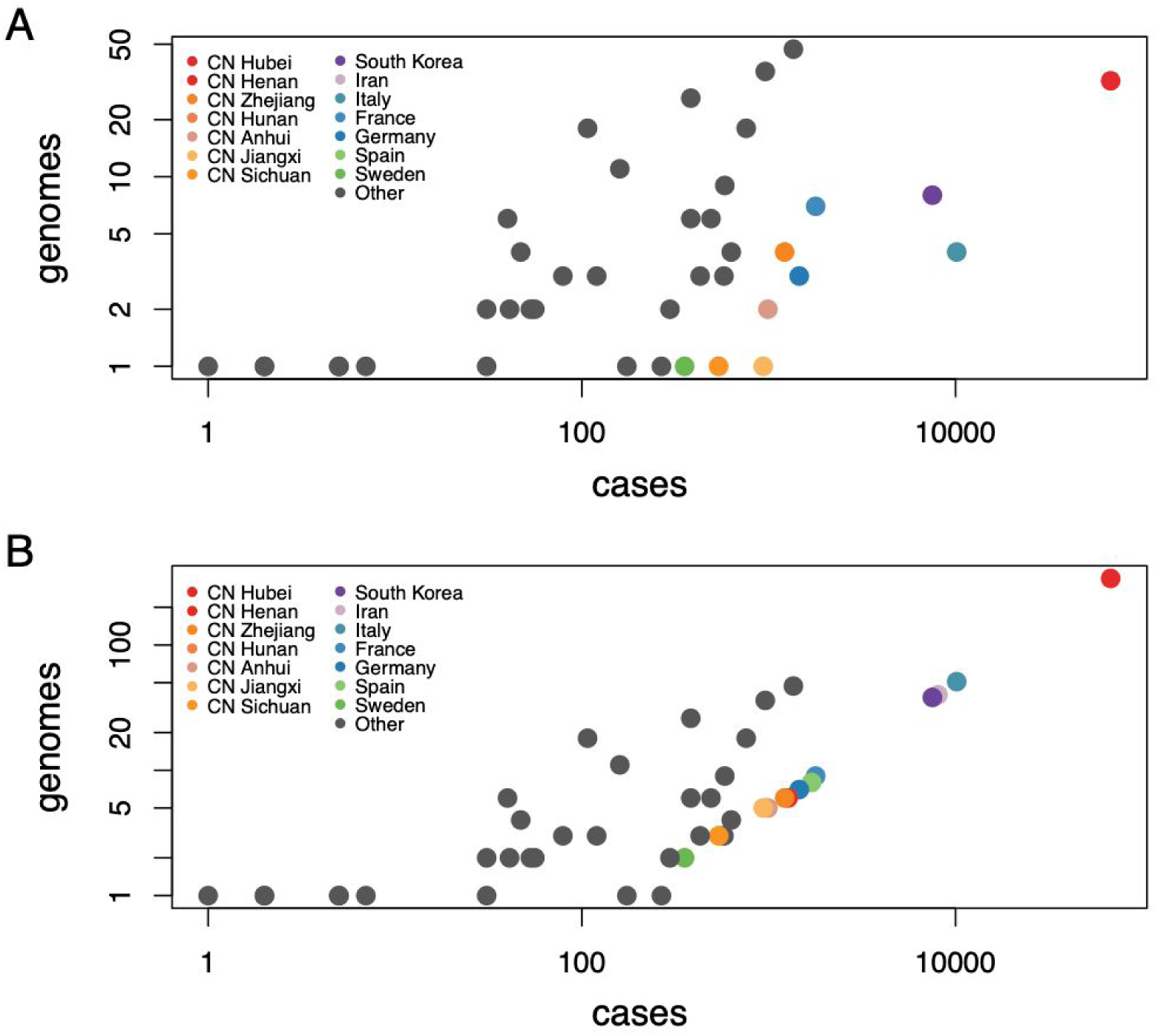
SARS-CoV-2 genomes and cases counts. (A) scatter plot of the number of available genomes against the number of cases on March 10 2020 on a log-log scale. (B) same as (A) but with additional unsampled taxa for 14 locations.

We integrate over all possible phylogenetic placements of such taxa using standard Markov chain Monte Carlo (MCMC) transition kernels. In the absence of sequence data, time of sampling represents an important source of information for the analysis in addition to sampling location. Here, we use epidemiological data in order to estimate a probabilistic distribution for the sampling times of unsampled taxa. Specifically, we follow Grubaugh et al. ^5^ in estimating the number of prevalent infectious individuals on day *t* (*P*_*t*_) by multiplying the number of incident infections up to day *t* by the probability that an individual who became infectious on day *i* was still infectious on day *t*:

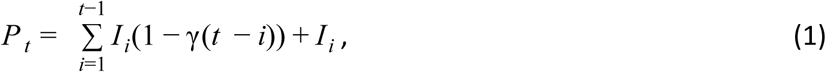

Where *γ*(*t-i*) is the cumulative distribution function of the infectious period. We also follow Grubaugh et al. ^5^ in modeling the infectious period as a gamma distribution with mean 7 days and standard deviation 4.5 days. Based on the estimated distributions of prevalent infections for the relevant locations over the time period of our analysis, we specify exponential or (truncated) normal prior distributions on the sampling times of unsampled taxa (Fig. 3), and integrate over all possible times using MCMC in the full Bayesian analysis.

**Figure 3:**
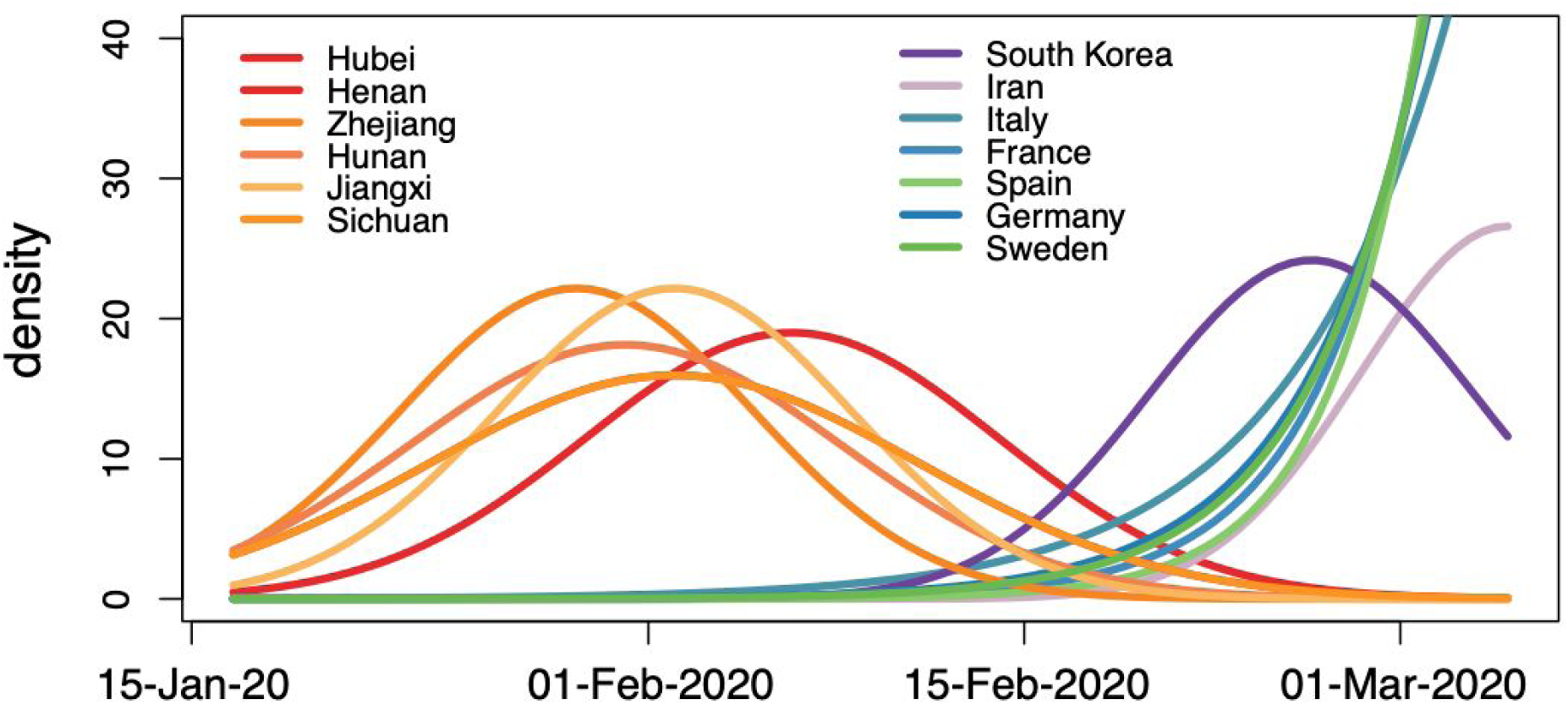
Prior probability distributions for the ages of the taxa representing unsampled diversity for different locations. The shapes of these probability distributions are based on estimated numbers of prevalent infections over time. The same normal prior probability distribution applies to taxa from Sichuan and Henan.

### Bayesian phylogeographic inference incorporating global mobility

We implement our approach to incorporate travel history in discrete phylogeographic inference in the BEAST framework (v.1.10.4 ^30^). In this framework, we assume that discrete trait data **X**, in our case location data associated with both sampled and unsampled taxa, and aligned molecular sequence data **Y** arise according to continuous-time Markov (CTMC) chain processes on a random phylogeny F with the following model posterior distribution:

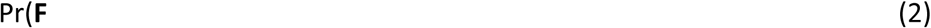

where Pr(**X** | **F**, **T**, Λ) and Pr(**Y** | **F**,ϕ) represent the discrete trait likelihood and sequence likelihood respectively, **T** is the additional time information for the ancestral nodes, Λ and *ϕ* characterize the discrete-trait and molecular sequence CTMC parameterizations along **F** respectively. Likelihoods Pr(**X** | **F**, **T**, Λ) and Pr(**Y** | **F**, *ϕ*) are calculated using Felsenstein’s Pruning algorithm ^31^, a computation that is efficiently performed using the high-performance BEAGLE library ^32^. We note that for the travel histories, the ancestral locations and times are introduced only for evaluating the discrete trait location likelihood Pr(**X** | **F**, **T**, Λ). The ancestral locations and times do not affect the sequence likelihood Pr(**Y** | **F**, *ϕ*), nor the likelihood of the coalescent model we use as our tree prior Pr(**F**).

For the sequence data, we use the HKY nucleotide substitution CTMC model ^33^ with a proportion of invariant sites and gamma-distributed rate variation among sites ^34^, an uncorrelated relaxed molecular clock model with a lognormal distribution ^35^ and an exponential growth coalescent tree prior. The uncertainty in collection date for 18 genomes was accommodated in the inference by integrating their age over the respective month of sampling. Our discrete location diffusion model involves 44 locations, represented by a limited number of sampled (and unsampled) taxa and ancestral nodes associated with travel locations. In order to avoid having to estimate a huge number of location transition parameters in a high dimensional CTMC, and to further inform the phylogenetic placement of unsampled taxa, we adopt a generalized linear model (GLM) formulation of the discrete trait CTMC that parametrizes the transition rates as a function of a number of potential covariates ^15^. As covariates, we consider i) air travel data, ii) geographic distance and iii) an estimable asymmetry coefficient for Hubei to account for the fact that the early stage of COVID-19 spread was dominated by importations from Hubei (with underdetected cases of COVID-19 probably having spread in most locations around the world ^36^). The air travel data consist of average daily symmetric fluxes between the 44 locations in January and February, 2013 (International Air Transport Association, http://www.iata.org). The geographic distance covariate only considers distances for pairs of locations in the same continent, which are based on centroid coordinates. We estimate the effect size of each of these covariates as well as their inclusion probability (specifying a 0.5 prior inclusion probability for each covariate).

We approximate the posterior distribution of our full probabilistic model using MCMC sampling. We run sufficiently long chains to ensure adequate effective sample sizes for continuous parameters as diagnosed using Tracer ^37^. We summarize posterior tree distributions using maximum clade credibility (MCC) trees and visualize them using FigTree. However, due to the limitations of single-tree representations when facing extensive phylogenetic uncertainty, we also propose new summaries below. A tutorial explaining how to perform travel-aware phylogeographic analyses in BEAST can be found at http://beast.community/travel_history.

### Phylogeographic visualizations

Due to the relatively low sequence variability over the short time scale of spread, phylogenetic reconstructions of SARS-CoV-2 are inherently uncertain, which also complicates inferring and interpreting location transition histories. If nodes in an MCC tree are associated with low posterior support, their conditional modal state annotation will be determined by a limited number of corresponding samples from the posterior tree distribution. The addition of unsampled taxa adds an additional challenge because the absence of sequence data makes them highly volatile in phylogenetic reconstruction reducing posterior node support to impractically low values for many nodes.

In order to marginalize over phylogenetic clustering in our visualization of phylogeographic history, we generate Markov jump estimates of the transition histories that are averaged over the entire posterior in our Bayesian inference ^15,38^. We study the ancestral transition history of specific taxa of interest by summarizing their Markov jump estimates as trajectories over time between a number of relevant states. A new BEAST tree sample tool (TaxaMarkovJumpHistoryAnalyzer available in the BEAST codebase at https://github.com/beast-dev/beast-mcmc) and associated R package constructs these estimates. We also visualize posterior expected Markov jumps estimates between all locations using circular migration flow plots, which have been successfully used to visualize migration data ^39^, including phylogeographic estimates ^40^. When summarizing these jumps from analyses that include unsampled diversity, we ignore branches that only have unsampled taxa as descendants. We only plot jumps that have a posterior expectation larger than 0.75.

## Results

### Travel history uncovers more realistic phylogeographic patterns

To focus on the early dynamics of SARS-CoV-2 spread, we analyze a data set consisting of curated genomes available in GISAID on March 10th, 2020 (*n* = 284). Having collected travel history data for over 20% of the sampled patients, we extend phylogeographic reconstruction methodology to incorporate this source of information (cfr. Methods). Specifically, we augment the sampled genomes from known travellers with their recent travel location and either the time of their return journey or estimated time of infection. Critically, we include travelers returning from severely undersampled locations, including Italy, Iran, and Hubei, China, the pandemic’s original epicenter.

In our Bayesian analysis of the complete data set, we model a discrete diffusion process between 44 locations: 29 countries and 15 locations within China, including 13 provinces, one municipality (Beijing) and one special administrative area (Hong Kong). We fit a GLM-parameterization of the discrete diffusion process and consider air travel data, within-continent geographic distances and an estimable asymmetry coefficient for transitions from and to Hubei as covariates for the diffusion rates. In Table 1, we compare the posterior estimates for the inclusion probabilities and conditional effect sizes (on a log scale) of these covariates in an analysis that incorporates (i) sampling location and travel history (travel-aware phylogeographic inference), (ii) sampling location only and (C) travel origin location. Regardless of what location data we use in the analyses, they consistently indicate that in this early stage, SARS-CoV-2 spread is shaped by air travel and not by geographic distance, and that there is strong asymmetric flow out of Hubei. Interestingly, this asymmetry is somewhat stronger for the analyses that incorporate travel locations compared to the analysis using only sampling locations. One explanation is that the majority of travellers were returning from Hubei, and adding this information contributes appropriately to the intensity of outflux from Hubei.

As expected for a low degree of sequence variability over this limited time range, our Bayesian phylogeographic reconstructions are burdened by a high degree of topological uncertainty. In Fig. 4, posterior support for the nodes in the MCC tree is represented by the size of the node circles, illustrating that only a limited number of clusters are reasonably well supported. This renders a single phylogenetic tree summary inadequate to interpret phylogeographic history. We side-step this problem by studying spatial trajectories that marginalize over phylogenetic variability, in the ancestral history of single taxa.

**Figure 4:**
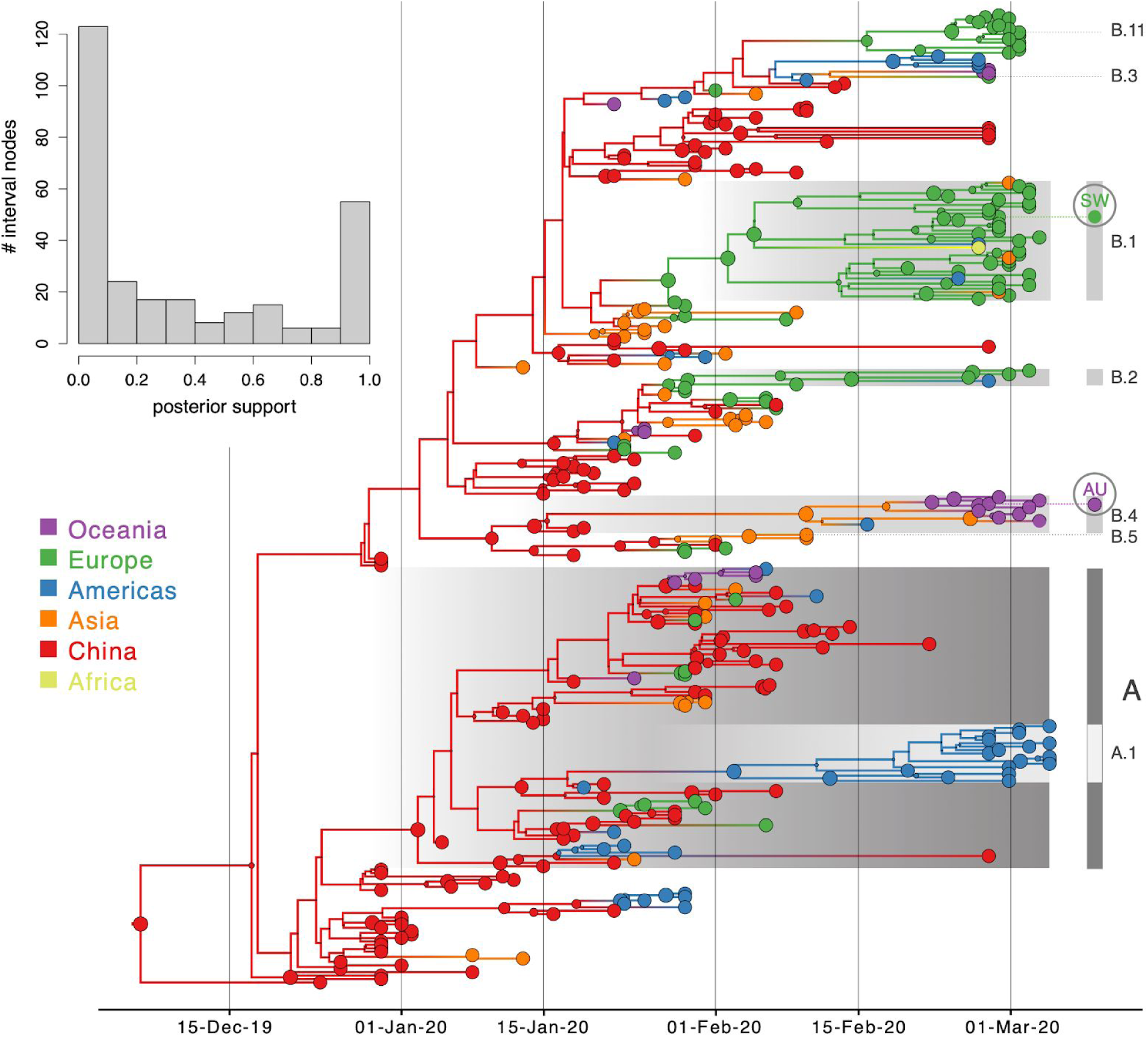
Bayesian phylogeographic reconstruction for the full data set incorporating travel history data. Although the analysis was performed using 44 location states, nodes and branches are shaded according to an aggregated color scheme for clarity (the reconstruction with a full color scheme can be found in the Supplementary information). Lineage classifications are highlighted for specific clusters: lineage A is embedded in lineage B. For lineage B, only specific sub-lineages are indicated. The taxa further investigated using trajectory plots are indicated at the tips of the trees. The inset represents a histogram of the node posterior support values. Viruses from Switzerland (SW) and Australia (AU) that are investigated as case studies are labeled.

In Fig. 5, we consider a case study involving the spatial path of a virus that was collected in Switzerland in February 2020 (EPI_ISL_413021). The Swiss virus is part of lineage B.1 (SW in Fig. 4) and is positioned within a cluster of viruses primarily from Europe that has been the subject of controversy. The basal clustering of a genome from the first detected case in Germany led to speculation that the virus spread from Germany to Italy ^41^. Using the standard location of sampling (Fig. 5 A), the trajectory plot traces the origins of the Swiss virus back to the Netherlands, an inference that is likely due to a relatively large number of Dutch genomes in the cluster. Going further back in time, the virus appears to have spread from Germany to the Netherlands. The original ancestry prior to Germany becomes uncertain, and could be Hubei, Guangdong, or other locations. Italy is not part of the trajectory at all because Italy is undersampled (only two genomes in the cluster). Using the locations of origin for the travellers (Fig. 5B), the trajectory is more ambiguous about the spatial path, and whether the Swiss virus came from Italy or the Netherlands. The travel-aware reconstruction, which includes both sampling location and traveler’s location of origin (Fig. 5C), almost fully resolves the ancestry of the Swiss virus. The Swiss virus was imported from Italy, and not the Netherlands. The fact that this cluster contains five genomes from travellers returning from Italy to various countries, including Germany, Scotland, Mexico, Nigeria, and Brazil, is instrumental in positioning Italy at the root of this cluster and helps correct for Italy’s lack of data. The inclusion of genomes associated with a Hubei travel history also strengthens the original Hubei ancestry in the phylogeography, as trajectories appear to coalesce earlier in Hubei. Although the trajectory suggests an introduction into Italy from Germany, the support for this is not overwhelming and solely due to the single genome sample from Germany basal to the Italian cluster. We return to this transmission hypothesis in the next section. In Supplementary Information, we include another trajectory for a virus in this cluster from a Brazilian traveller (EPI_ISL_412964) returning from Italy that confirms the pattern inferred for the Swiss sample.

**Figure 5:**
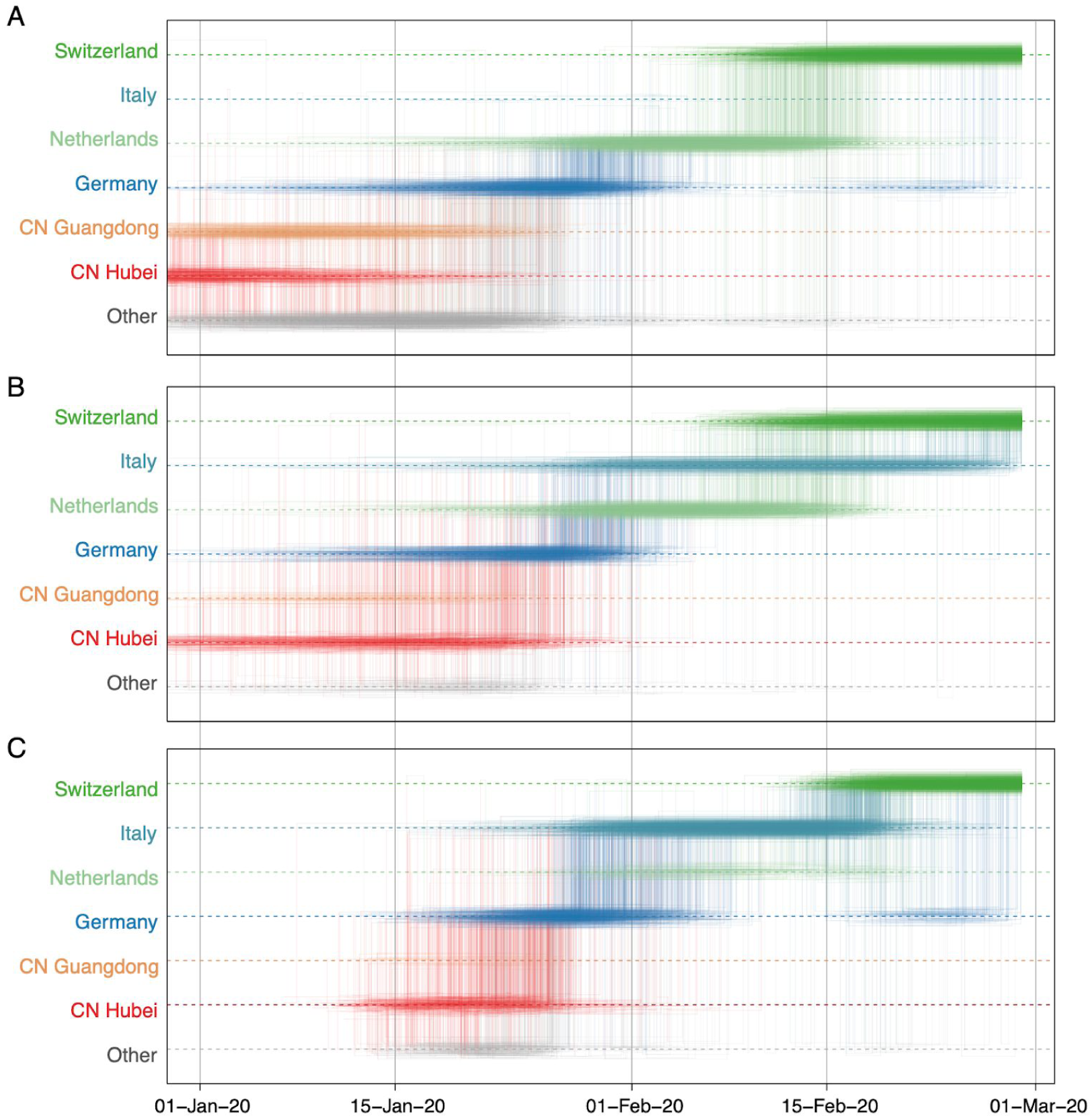
Markov jump trajectory plot depicting the ancestral transition history between locations from Hubei up the sampling location for a Swiss genome (EPI_ISL_413021) in lineage B1 using (A) sampling location only, (B) travel origin location and (C) sampling location and travel history. The trajectories are summarized from a posterior tree distribution with Markov jump history annotation.

In a second case study, we consider a virus from Australia sampled in February 2020 (EPI_ISL_412975). The virus is positioned within a clade of other closely related viruses from Australia (lineage B.4, Fig. 4), some of which were sampled from travellers returning from Iran ^42^. Using sampling location alone does not provide any support for Iranian ancestry, since the data set does not include any genomes directly sampled from Iran (Fig. 6A). Using the locations of origin for the travellers does support Iranian ancestry (Fig. 6B), but with considerable ambiguity. However, the travel-aware reconstruction, including both sampling location and traveler’s location of origin, clearly supports an ancestry that includes Iran (Fig. 6C). This Iran-Australia case study provides an example where enforcing the travel location somewhat deeper in the evolutionary history (at return dates of the travellers) imparts more information that is critical for correctly reconstructing ancestral relationships.

**Figure 6:**
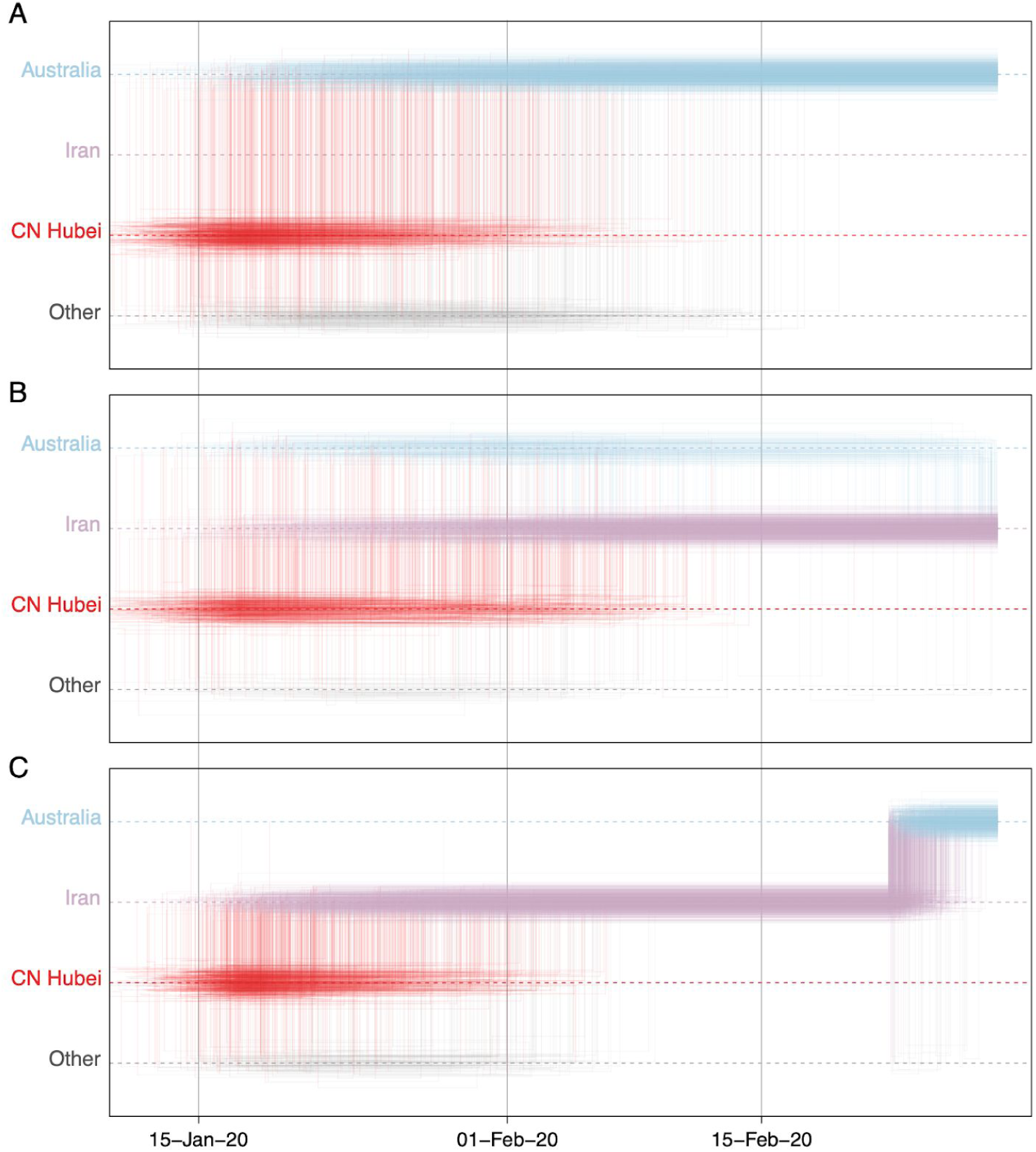
Markov jump trajectory plot depicting the ancestral transition history between locations from Hubei up the sampling location for an Australian genome (EPI_ISL_412600) in lineage B.4 using (A) sampling location only, (B) travel origin location and (C) sampling location and travel history. The trajectories are summarized from a posterior tree distribution with Markov jump history annotation.

### Unsampled diversity reinforces reconstructions informed by travel data and unveils alternative transmission hypotheses

To further explore the sensitivity of phylogeographic analyses to sampling bias, we incorporate unsampled taxa in our reconstructions, in addition to travel history data. We add unsampled taxa for locations that are undersampled according to case counts (cfr. Methods), in this case primarily for Hubei (*n* = 307), followed by Italy (*n* = 47), Iran (*n* = 40), and South Korea (*n* = 30). We specify a prior distribution over their tip ages (‘sampling times’) based on estimates of prevalent infections (cfr. Methods). Using this framework, we revisit the trajectory estimate for the Swiss B.1 genome (Fig. 7). In contrast to the reconstructions with no unsampled taxa (Fig. 5), we now mainly observe a direct transition from Hubei to Italy (posterior probability = 0.88), implying that a second introduction from Hubei that is independent from the introduction into Germany may have seeded the Italian clade. This hypothesis arises from the inclusion of unsampled Hubei taxa that now cluster between the German virus and the Italian clade (Fig. 7), even though the branch connecting the German genome to the Italian clade only represents a single substitution. The many unsampled Italian taxa that fall in this clade further reinforce its Italian ancestry.

**Figure 7:**
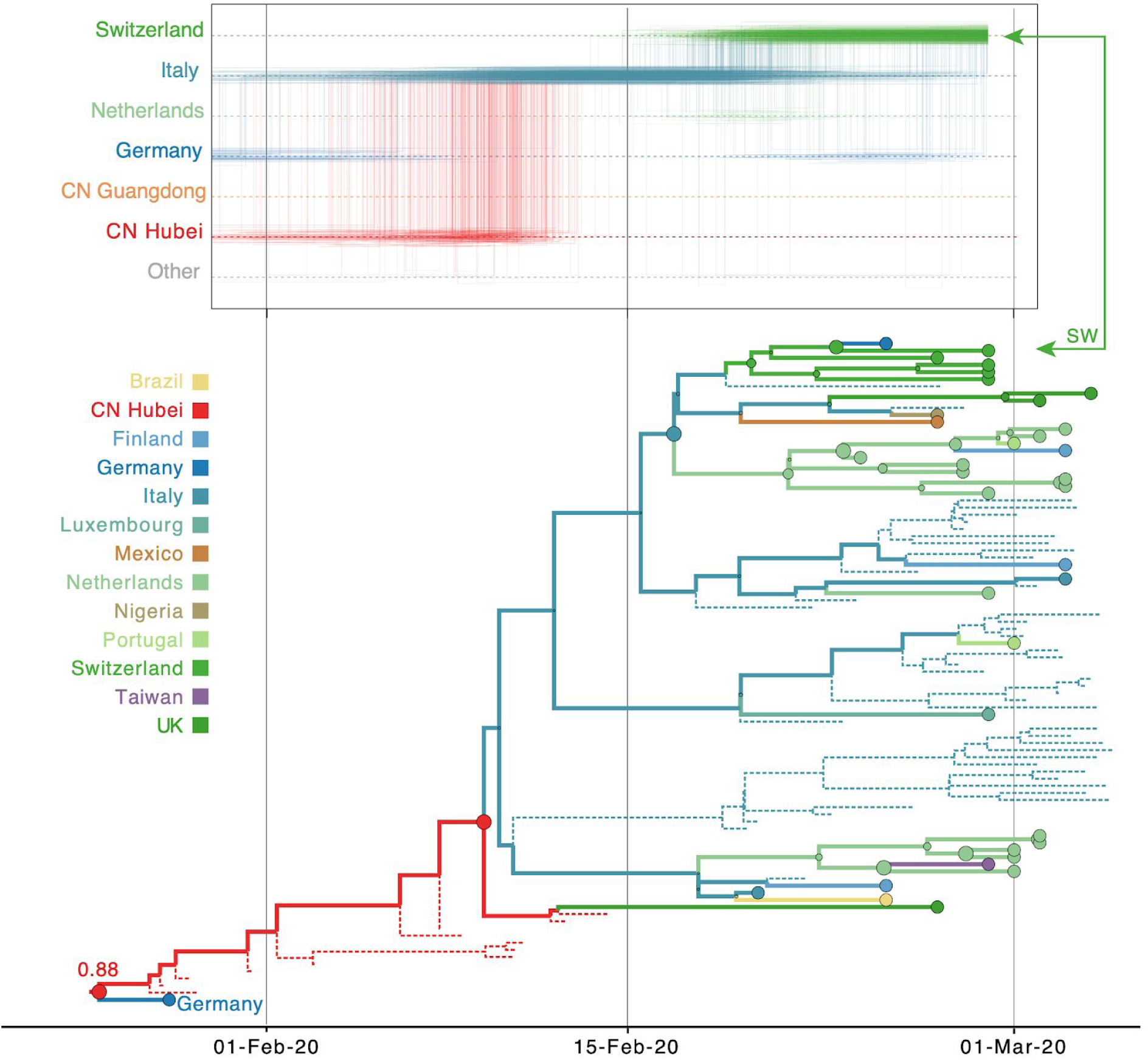
Markov jump trajectory plot as in Fig. 5 for a Swiss genome (EPI_ISL_413021) in lineage B1 and B1 subtree for the Bayesian phylogeographic analysis incorporating travel data Fig. 7 and unsampled diversity. Dotted lines represent branches associated with unsampled taxa assigned to Italy and Hubei, China. The tip for the Swiss genome corresponding to the trajectory is indicated with an arrow. Because of the color similarity between Italy and Germany, the basal German virus is labeled. The value at the root represents the posterior location state probability.

The inclusion of unsampled taxa also provides more resolution on the Hubei ancestry of the Iran-Australia case study in lineage B.4 (EPI_ISL_412975). We estimate a similar trajectory as the analysis using travel history without unsampled taxa (Fig. 6), but with a more recent coalescence in Hubei because of unsampled Hubei taxa clustering basal to an Iranian clade (Fig. 8). The genomes from four Australian travellers returning from this country, one direct contact, another Australian genome without travel history, as well as a genome from a traveller returning from Iran to New Zealand are effectively embedded in unsampled Iranian diversity. The most basal virus in this clade is from a Canadian traveller returning from Iran. Although the basal nature of this virus is not well supported, there is good posterior support for the monophyly of all the sampled genomes in this clade. Notably, this virus was sampled before the first report of COVID-19 in Iran on February 19^th 43^, but our reconstruction indicates that considerable diversification, and hence transmission, already took place prior to this report.

**Figure 8:**
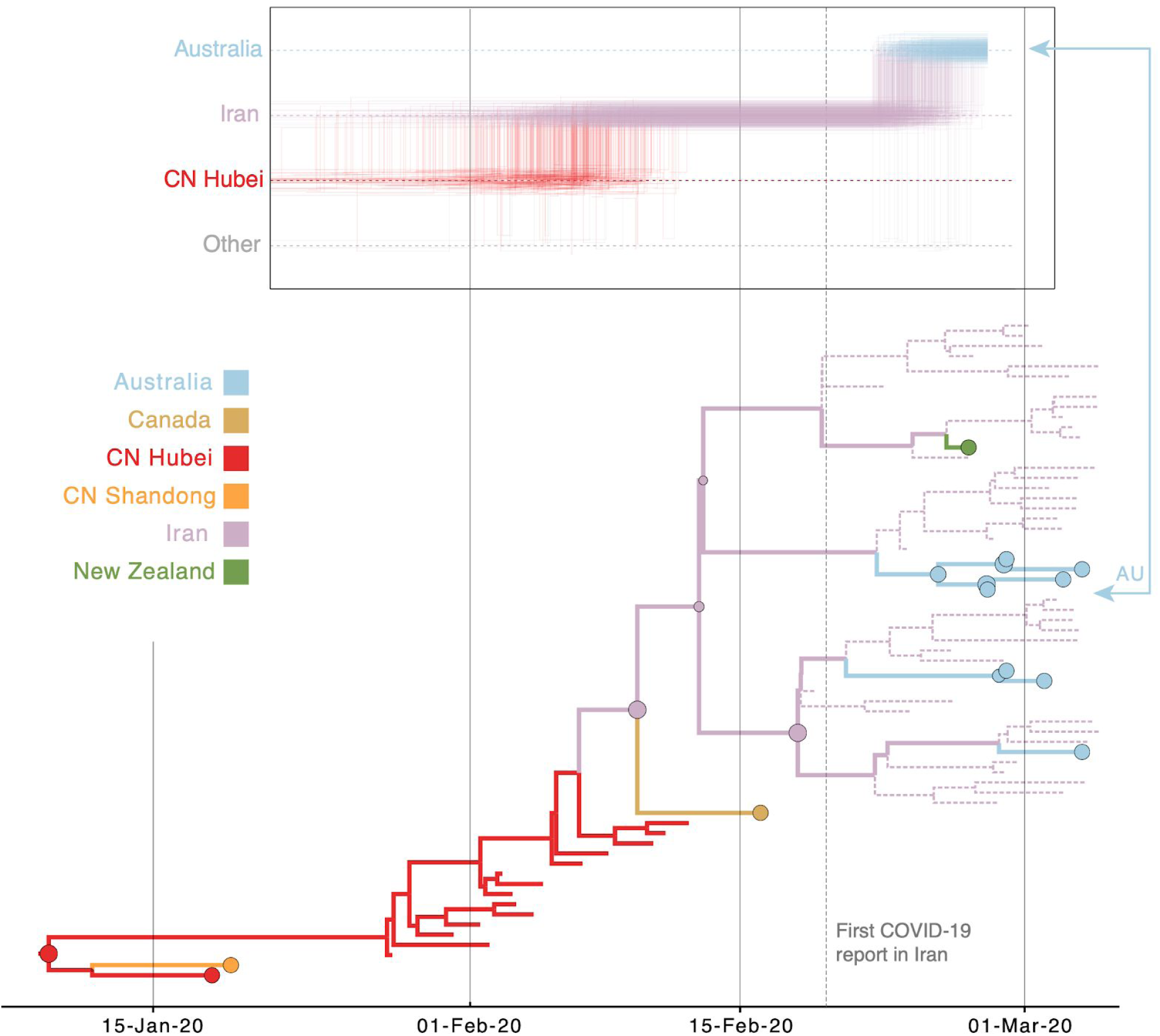
Markov jump trajectory plot as in Fig. 5 for the Australian genome (EPI_ISL_412600) in lineage B.4 and B4 subtree for the Bayesian phylogeographic analysis incorporating travel data and unsampled diversity. Dotted lines represent branches associated with unsampled taxa assigned to Iran and Hubei, China. The tip for the Australian genome corresponding to the trajectory is indicated with an arrow.

In addition to focusing on specific transmission patterns, we also summarize the overall dispersal dynamics in a way that marginalizes over plausible phylogenetic histories for the four different analyses we perform (Fig. 9). While introductions from Hubei represent the dominant pattern when using sampling location only (Fig. 9A), this is far more pronounced for the other analyses. Using sampling location suggests unrealistic dispersal from locations such as Australia and The Netherlands that largely disappear in the other analyses, specifically when using travel history data without or with unsampled diversity (Fig. 9C & D). In the travel-aware analyses, European countries experience more introductions from Italy and to a lesser extent also more directly from Hubei. As also illustrated by the specific examples (Fig. 5 & 6), the considerable number of secondary transmissions from Italy and Iran is revealed by using travel history data; the substantial addition of unsampled taxa from these relatively undersampled locations does not further contribute to this pattern. However, adding unsampled diversity from Hubei suggests that occasional introductions from other locations are more likely direct introductions from Hubei, such as introductions from Guangdong Province to Shandong Province, South Korea and Japan, and a supposedly Canadian introduction in the US.

**Figure 9:**
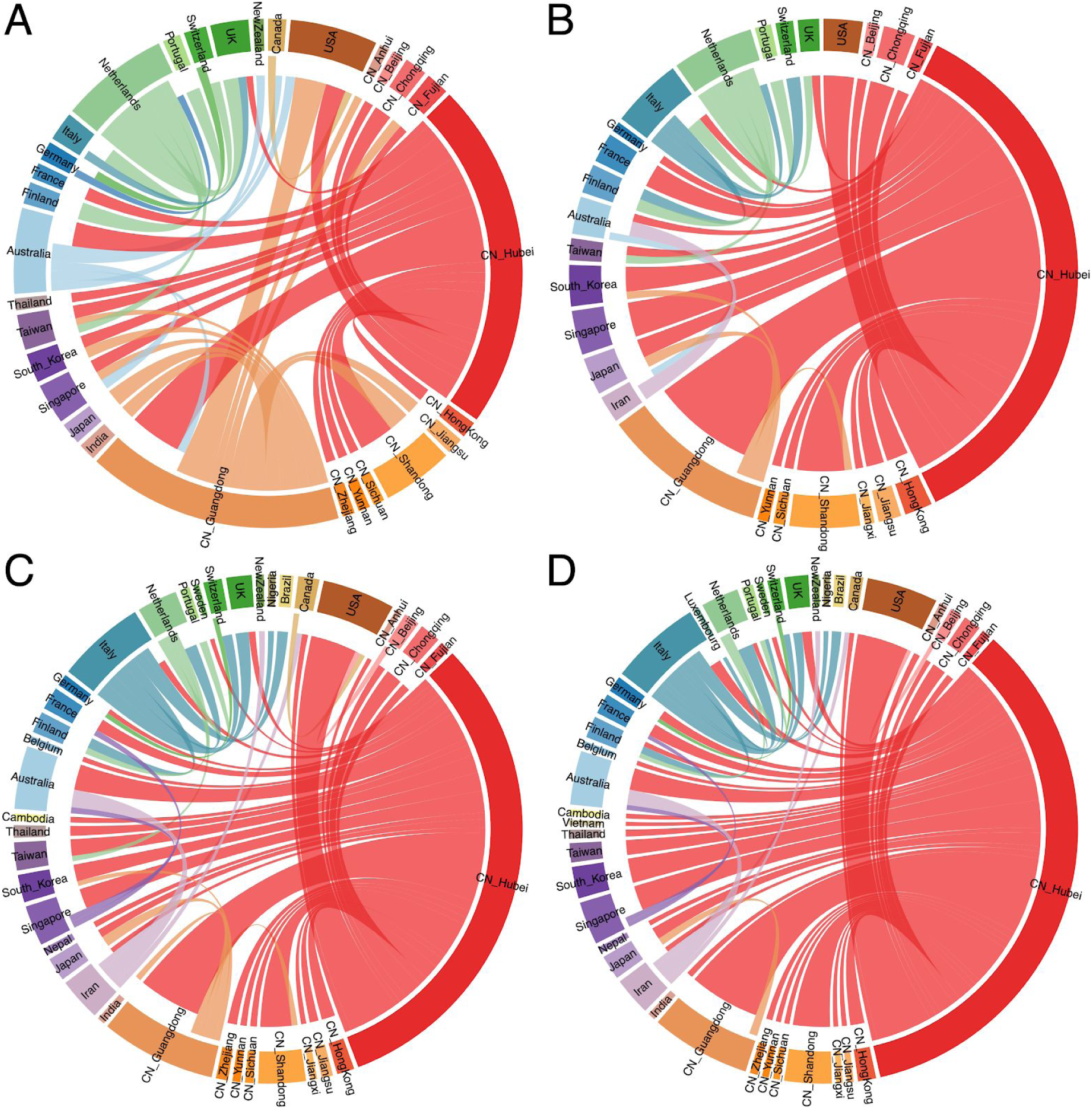
Circular migration flow plots summarizing Markov jump estimates for the analyses using (A) sampling location only, (B) travel origin location, (C) sampling location and travel history and (D) sampling location and travel history, with unsampled diversity. The plots show the relative number of transitions between locations. The direction of the transitions are encoded both by the origin colour and by the gap between link and circle segment at the destination (migration into a location is associated with a larger gap than migration out of a location). As part of the Supplementary Information, we include the same figure but with a transparency for transitions from Hubei to emphasize the ‘secondary’ dispersal dynamics.

## Discussion

International travelers had a central role in the early global spread of the SARS-CoV-2 virus. To track whether COVID cases were new imports or community transmission, detailed travel histories were collected from many of the early COVID patients. To date, however, phylogeographic approaches using discrete trait reconstruction have not been able to fully incorporate travel history data. Researchers had to select whether to assign a sample to the location of sampling, typically the home country, or to the location visited by the traveler. Either way, half of the information was lost. During a period when there were major gaps in the availability of SARS-CoV-2 genomes from many key locations, losing half of the spatial information provided by travelers has been suboptimal. Here, by developing a new phylogeographic approach that introduces ancestral nodes in the phylogeny that are associated with locations visited by travelers, we provide a method to formally recapture all the rich information provided by travelers. Most importantly, we demonstrate that the travel-aware approach can dramatically improve phylogeographic inferences about the specific country-to-country paths followed by the SARS-CoV-2 virus during the early stages of global spread. As expected, the inclusion of travel history data is most informative when travelers are arriving from locations such as Italy and Iran that experienced early SARS-CoV-2 outbreaks, but for which few genome sequences were available, resulting in large gaps in the phylogeny.

We are currently studying a pandemic as it unfolds, and as new SARS-CoV-2 sequences continue to be generated globally at an explosive pace, there will be ongoing opportunities to reassess the probability of conclusions drawn from earlier data sets. More genome data from Hubei has already become available which will reduce the need for unsampled diversity from the pandemic origin in phylogeographic reconstructions. We have also observed that two genomes from Iran are now available in GISAID that cluster with the genomes from Australian travellers returning from this country (data not shown), which reinforces the Iran-Australia connection observed in our travel history reconstructions. We anticipate that the generation of additional data from Italy will also support the spatial connections inferred here from traveler data. However, we intentionally tested the performance of our methods using an early SARS-CoV-2 data set that was heavily burdened by spatiotemporal sampling bias. It is possible that our methods could not account for all gaps in data at this stage, and further exploration is needed of how well our methods perform at different magnitudes of sampling gaps.

We retrieved travel history data for about 20% of the early available genomes, but many more travel-based introductions may be undocumented. While such information is sometimes available in GISAID records, this is not commonly included metadata and there is otherwise no specific resource available to retrieve such information. This may at least partly be explained by concerns about the risk of patient identification. Travel history data may be particularly important when analyzing low diversity data using Bayesian joint inference of sequence and traits because sharing the same location state can contribute to the phylogenetic clustering of taxa. In general, it is crucial to consider the uncertainty of Bayesian time-measured phylogenetic reconstructions because, even for sparse sequence information, a single tree sample (including the MCC tree) will appear highly resolved with branching structures that may not be supported by substitutions (Fig. 4). However, this is only a single transmission hypothesis compatible with the data, while many other hypotheses will be plausible as should be reflected by the different topologies in the posterior and hence by low node support values. For this reason, we resorted to posterior summaries that focus on the location transition patterns in the ancestral history of single taxa or on Markov jump estimates, both averaging over all plausible trees.

While downsampling genomic data from locations in unbalanced data sets has become a common practice ^15,18^, we present here an alternative approach that adds unsampled taxa to assess the sensitivity of inferences to sampling bias. We emphasize that even though the inclusion of unsampled taxa is informed by epidemiological data, these unsampled taxa should never be considered as additional observations. The unsampled diversity reveals alternative hypotheses that may not be captured by the available genome sampling but are worth considering in the context of biased sampling. However, reconstructions using unsampled taxa do not provide evidence for any single hypothesis with the same weight as actual genomic data. We envision reconstructions built with unsampled taxa as being exploratory in nature, and most useful as added support for conclusions drawn independently from other analytical approaches, for example evolutionary simulations or epidemiological studies.

It is important for future users of these methods to understand exactly how different kinds of empirical data are used to determine where unsampled taxa will attach to the phylogenetic backbone of sampled genomes. The first important aspect is the relative positioning in time of unsampled tips, which in our case is drawn from distribution curves of estimated prevalent infections over time. So, this together with the relative abundance of unsampled taxa by location is informed by epidemiological data. Second, the locations of unsampled taxa will determine their clustering when jointly inferring the phylogeny based on sequences and discrete location traits. In this respect unsampled taxa will preferentially cluster with taxa representing the same locations, either sampled genomes or other unsampled ones. However, unsampled taxa can, and do in our experience, branch off lineages representing different location states. The relative preference for which location transitions this involves will be determined by the matrix of transition rates of the discrete trait CTMC, which in our case are informed by covariates such as air travel. This implies that in the SARS-CoV-2 analysis unsampled taxa can be positioned with taxa from countries that are highly connected by airline travel, given the importance of air travel in the early spread of the virus. Finally, the branch lengths, or time it takes for unsampled taxa to find a common ancestor with other taxa, will be influenced by the coalescent prior. We opted for a simple exponential growth coalescent prior in our analyses, but more flexible tree priors are available that can also be informed by epidemiological data ^e.g. 44^.

By formally accommodating the possibility of unsampled diversity in our phylogenetic reconstructions, we provide alternative scenarios for how SARS-CoV-2 spread globally and entered specific countries. Most importantly, many early introductions in different locations were likely from Hubei, in line with modelling estimates that point at the underdetection of exported COVID-19 cases from Wuhan ^36^. In addition, our findings reinforce estimates shaped by travel history that point at early introductions into both Italy and Iran, two countries that are not well represented by genomic sampling, and subsequent transmission events from these countries to other locations. Due to the low genomic variability of SARS-CoV-2, it may be more appropriate to refer to unsampled transmission chains rather than unsampled diversity because many unsampled taxa may represent highly similar or even identical genomes. With the large number of SARS-CoV-2 genomes now available, the question arises how scalable the incorporation of unsampled taxa will be. For computationally expensive Bayesian inferences, the approach may need to go hand in hand with downsampling procedures or more detailed examination of specific sublineages. Furthermore, as averaging over all plausible phylogenetic ‘placements’ of large numbers of ‘volatile’ unsampled taxa can be a challenging task, further developments are needed to make the estimation more efficient for larger scale datasets.

We firmly acknowledge that many aspects of these analyses require further detailed examination and refinement in other pathogen systems with different types of data gaps and sampling biases. For example, we used an arbitrary cut-off for the ratio of available genomes to case counts in order to decide which locations required representation by unsampled taxa (without accounting for differences in reporting rates for case counts), so it would be useful to investigate how sensitive the reconstructions are with respect to such decisions. Our spatial diffusion GLM may benefit from 2020 travel data that are impacted by travel restrictions. The time dependency imposed by travel restrictions could potentially be modelled with an epoch version of the discrete trait CTMC ^25^. This could also be important for the asymmetry factor we included for transitions from Hubei as these will be severely impacted by the travel ban imposed on January 23 ^45^. Finally, in addition to investigating the impact of model and data adjustments on empirical examples, simulation studies would greatly assist in evaluating the performance of the phylogeographic reconstructions in controlled scenarios.

In conclusion, we demonstrate how travel history data can be formally integrated into discrete phylogeographic reconstructions and that this, together with accounting for unsampled diversity, can mitigate spatiotemporal sampling bias in reconstructions of the early spread of SARS-CoV-2. More research is needed on the specifications of such analyses and we hope that this work will stimulate developments to further integrate epidemiological information and other data sources into phylodynamic reconstructions.

## Acknowledgments

We would like to thank all the authors who have kindly shared genome data on GISAID, and we have included a table (Supplementary Table S2) outlining the authors and institutes involved.

The research leading to these results has received funding from the European Research Council under the European Union’s Horizon 2020 research and innovation programme (grant agreement no. 725422-ReservoirDOCS) and from the European Union’s Horizon 2020 project MOOD (grant agreement no. 874850). The Artic Network receives funding from the Wellcome Trust through project 206298/Z/17/Z. PL acknowledges support by the Research Foundation - Flanders (‘Fonds voor Wetenschappelijk Onderzoek - Vlaanderen’, G066215N, G0D5117N and G0B9317N). GB acknowledges support from the Interne Fondsen KU Leuven / Internal Funds KU Leuven under grant agreement C14/18/094, and the Research Foundation – Flanders (‘Fonds voor Wetenschappelijk Onderzoek - Vlaanderen’, G0E1420N). MAS and KGA acknowledge support from National Institutes of Health grant U19 AI135995. We also gratefully acknowledge support from NVIDIA Corporation with the donation of parallel computing resources used for this research. This work was supported by the Multinational Influenza Seasonal Mortality Study (MISMS), an on-going international collaborative effort to understand influenza epidemiology and evolution, led by the Fogarty International Center, NIH. The content is solely the responsibility of the authors and does not necessarily represent official views of the National Institutes of Health.

## Notes

### Competing Interest Statement

The authors have declared no competing interest.

