## Supplementary Information for "Accommodating individual travel history, global mobility, and unsampled diversity in phylogeography: a SARS-CoV-2 case study"

**Affiliations:**

**Bayesian Evaluation of Temporal Signal (BETS)**

As part of the BETS analysis to assess temporal signal in the data, we estimate the performance of the strict clock and the relaxed clock with an underlying lognormal distribution, with and without making use of the sequence sampling times. We make use of generalized stepping-stone samling while accommodating phylogenetic uncertainty [1] to estimate log marginal likelihoods for these four models in BEAST 1.10 [2], running an initial Markov chain of 25 million iterations followed by 100 power posteriors of 1 million iterations each.

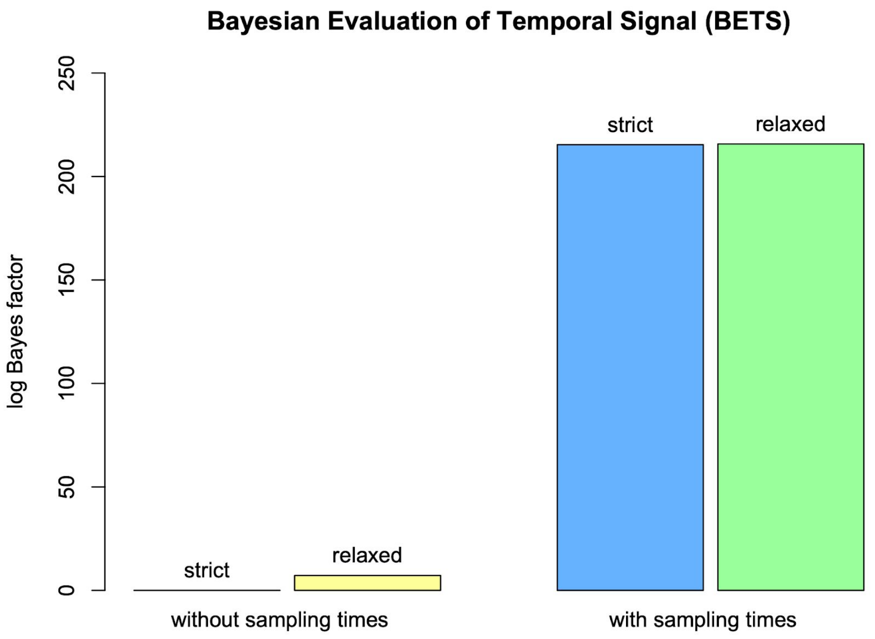


Figure S1: BETS reveals very strong evidence (log Bayes factors > 215) in favor of the models including sampling times, indicative of strong temporal signal in the data. The relaxed clock offers a marginal improvement of 0.3 log units in fit over the strict clock when using sampling times.

Figure S1 summarizes the results of the BETS analysis, which were repeated across multiple independent replicates and with increasing computational settings. When including the sampling times in the analysis of the data, model fit increases significantly and yields very strong evidence (log Bayes factors > 215) in favor of using the sampling time information [3], with strict and relaxed clock offering near-identical performance (log Bayes factor of 0.3 in favor of the latter).

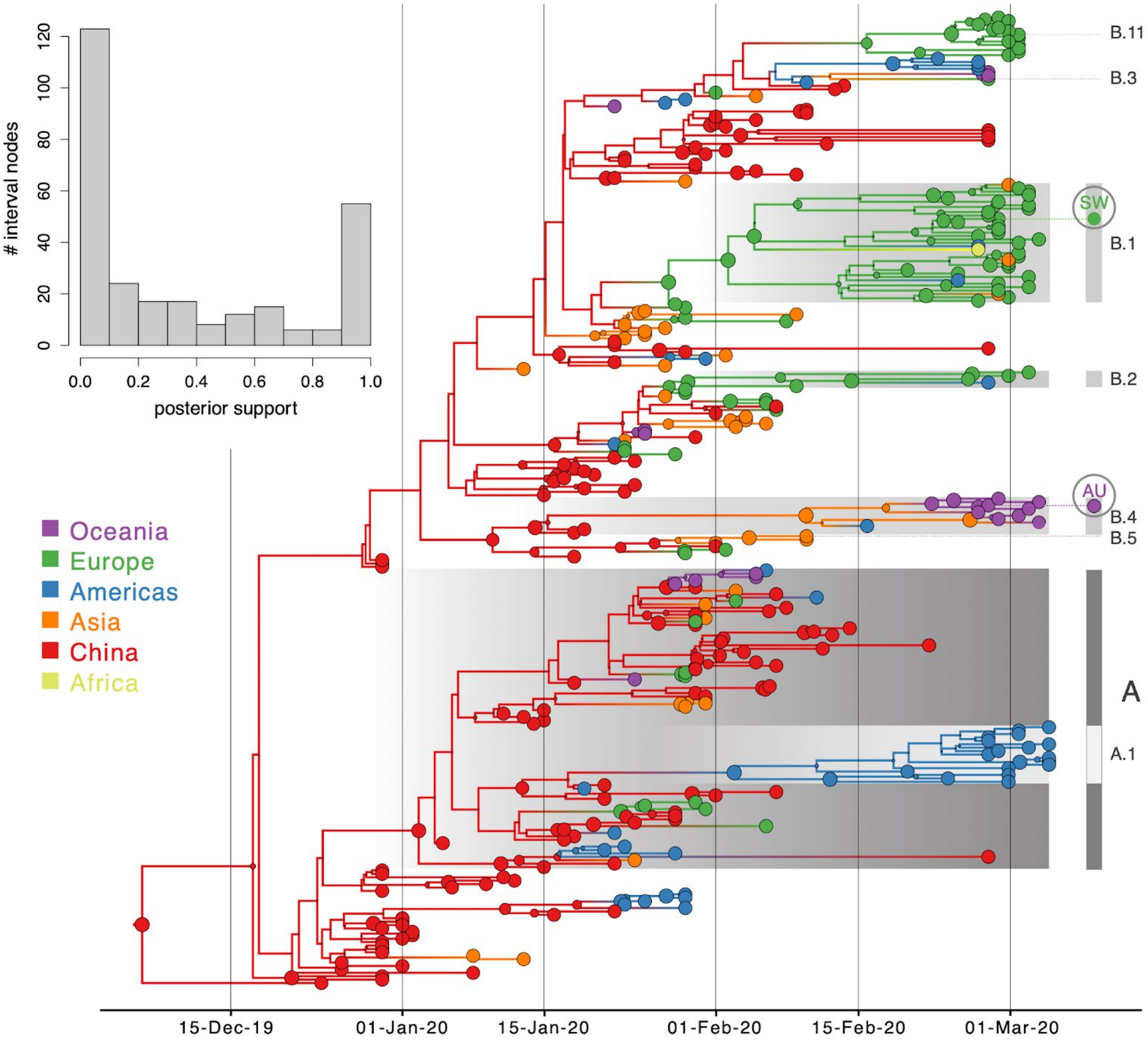


Figure S2. Bayesian phylogeographic reconstruction for the full data set incorporating travel history data. This figure mirrors Fig.4 in the main manuscript, but uses color shading for all the locations included in the analysis.

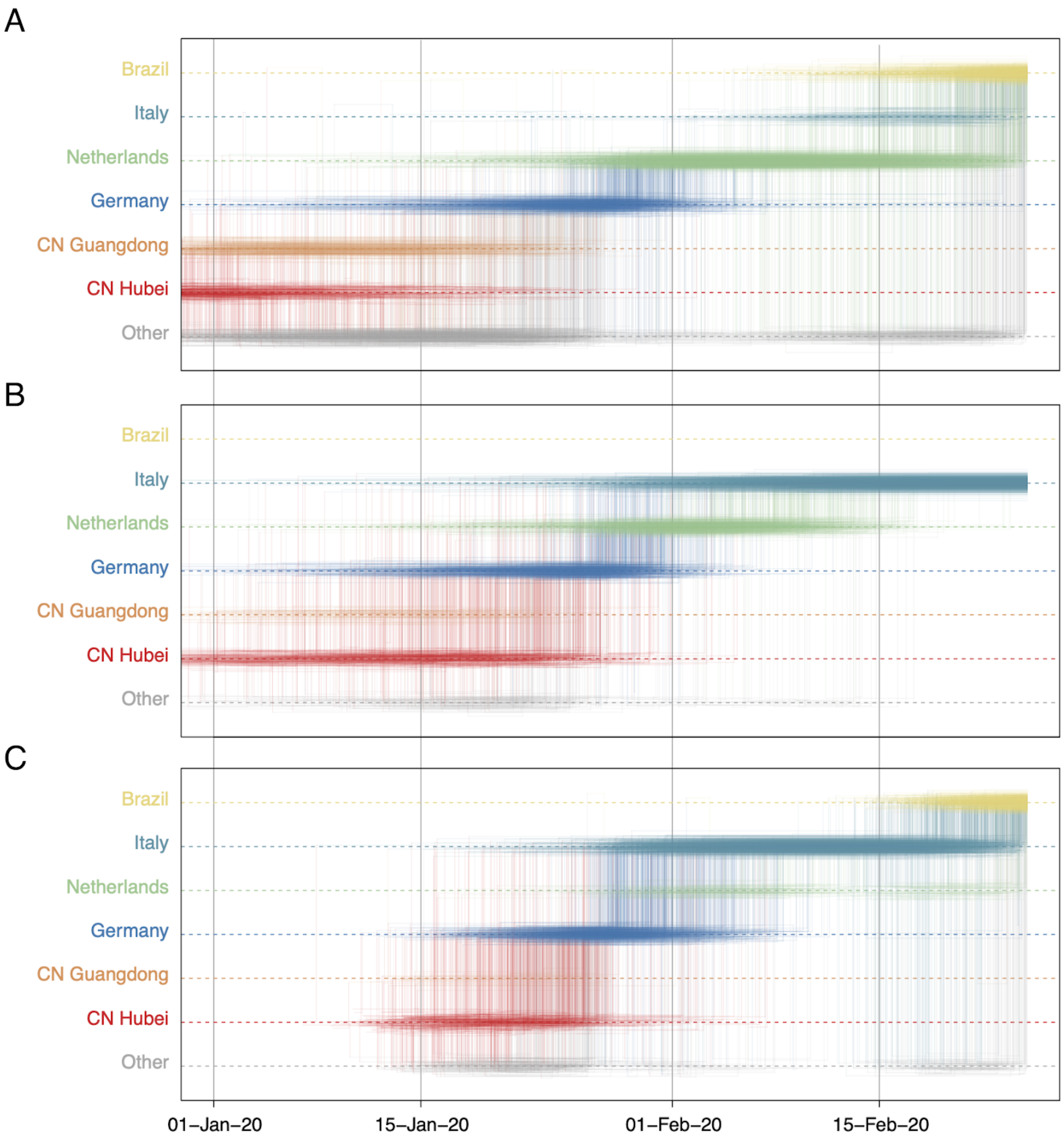


Figure S3. Markov jump trajectory plot depicting the ancestral transition history between locations from Hubei up the sampling location for a Brazilian genome (EPI_ISL_412964) in lineage B1 using (A) sampling location only, (B) travel origin location and (C) sampling location and travel history. The trajectories are summarized from a posterior tree distribution with Markov jump history annotation

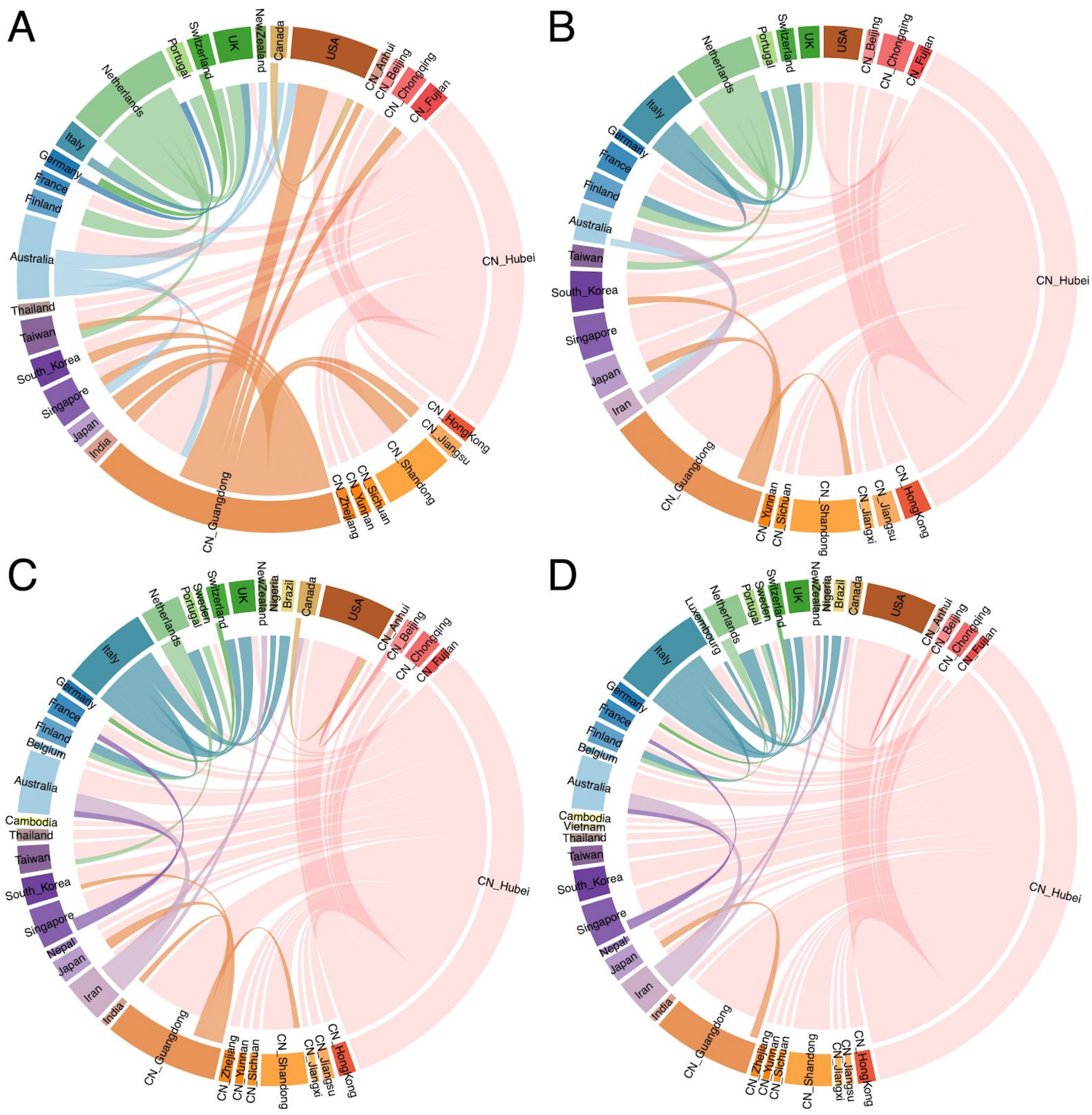

Figure S4. Circular migration flow plots summarizing Markov jump estimates for the analyses using (A) sampling location only, (B) travel origin location, (C) sampling location and travel history and (D) sampling location and travel history, with unsampled diversity. This figure mirrors Fig. 9 in the main manuscript, but uses transparency for transitions from Hubei to emphasize the 'secondary' migration dynamics.

**Table S1.** Travel history data associated with 64 of the 284 genomes.

| GISAID accession number | sampling location | sampling date | travel location | Travel date (days before collection date) |
|---|---|---|---|---|
| EPI_ISL_403932 | CN Guangdong | 2020-01-14 | CN Hubei | 16 |
| EPI_ISL_403936 | CN Guangdong | 2020-01-17 | CN Hubei | 6 |
| EPI_ISL_403937 | CN Guangdong | 2020-01-18 | CN Hubei | 6 |
| EPI_ISL_403962 | Thailand | 2020-01-08 | CN Hubei | 1* |
| EPI_ISL_403963 | Thailand | 2020-01-13 | CN Hubei | 1* |
| EPI_ISL_404227 | CN Zhejiang | 2020-01-16 | CN Hubei | NA |
| EPI_ISL_404228 | CN Zhejiang | 2020-01-17 | CN Hubei | NA |
| EPI_ISL_404253 | USA | 2020-01-21 | CN Hubei | 12 |
| EPI_ISL_404895 | USA | 2020-01-19 | CN Hubei | 9 |
| EPI_ISL_406034 | USA | 2020-01-23 | CN Hubei | NA |
| EPI_ISL_406036 | USA | 2020-01-22 | CN Hubei | NA |
| EPI_ISL_406223 | USA | 2020-01-22 | CN Hubei | NA |
| EPI_ISL_406597 | France | 2020-01-23 | CN Hubei | NA |
| EPI_ISL_406844 | Australia | 2020-01-25 | CN Hubei | 6 |
| EPI_ISL_406862 | Germany | 2020-01-28 | CN Hubei (2) | 9 + x (2) |
| EPI_ISL_406970 | CN Zhejiang | 2020-01-20 | CN Hubei | NA |
| EPI_ISL_406973 | Singapore | 2020-01-23 | CN Hubei | NA |
| EPI_ISL_407079 | Finland | 2020-01-29 | CN Hubei | 6 |
| EPI_ISL_407313 | CN Zhejiang | 2020-01-19 | CN Hubei | NA |
| EPI_ISL_407893 | Australia | 2020-01-24 | CN Hubei | 6 |

| | | | | |
|---|---|---|---|---|
| EPI_ISL_407894 | Australia | 2020-01-28 | CN Hubei | 1 |
| EPI_ISL_407896 | Australia | 2020-01-30 | CN Hubei | 3 |
| EPI_ISL_407976 | Belgium | 2020-02-03 | CN Hubei | 2 |
| EPI_ISL_408008 | USA | 2020-01-29 | CN Hubei | NA |
| EPI_ISL_408010 | USA | 2020-01-29 | CN Hubei | 6 |
| EPI_ISL_408489 | Taiwan | 2020-01-31 | CN Hubei | 11 |
| EPI_ISL_408668 | Vietnam | 2020-01-24 | CN Hubei | 7 |
| EPI_ISL_408670 | USA | 2020-01-31 | CN Beijing | NA |
| EPI_ISL_408976 | Australia | 2020-01-22 | CN Hubei | 2 |
| EPI_ISL_408977 | Australia | 2020-01-25 | CN Hubei | 2 |
| EPI_ISL_409067 | USA | 2020-01-29 | CN Hubei | 1 |
| EPI_ISL_410218 | Taiwan | 2020-02-05 | CN Hubei | NA |
| EPI_ISL_410301 | Nepal | 2020-01-13 | CN Hubei | 1* |
| EPI_ISL_410486 | France | 2020-02-08 | Singapore (2) | 15 (2) |
| EPI_ISL_410545 | Italy | 2020-01-29 | CN Hubei | 6 |
| EPI_ISL_410717 | Australia | 2020-02-05 | CN Hubei | 9 |
| EPI_ISL_410718 | Australia | 2020-02-05 | CN Hubei | 9 |
| EPI_ISL_411902 | Cambodia | 2020-01-27 | CN Hubei | 4 |
| EPI_ISL_411929 | South Korea | 2020-01 | CN Hubei | NA |
| EPI_ISL_411951 | Sweden | 2020-02-07 | CN Hubei | 14 |
| EPI_ISL_411954 | USA | 2020-02-06 | CN Hubei | 1 |
| EPI_ISL_411955 | USA | 2020-02-10 | CN Hubei | 5 |
| EPI_ISL_411956 | USA | 2020-02-11 | CN Hubei | 6 |
| EPI_ISL_412116 | UK | 2020-02-09 | China | 2 |
| EPI_ISL_412912 | Germany | 2020-02-25 | Italy | NA |

| | | | | |
|---|---|---|---|---|
| EPI_ISL_412964 | Brazil | 2020-02-25 | Italy | 4 |
| EPI_ISL_412965 | Canada | 2020-02-16 | Iran | 7 |
| EPI_ISL_412971 | Finland | 2020-02-25 | Italy | 2 |
| EPI_ISL_412972 | Mexico | 2020-02-27 | Italy | 5 |
| EPI_ISL_412975 | Australia | 2020-02-28 | Iran | 6 |
| EPI_ISL_413014 | Canada | 2020-01-25 | CN Hubei | 7 |
| EPI_ISL_413015 | Canada | 2020-01-23 | CN Hubei | NA |
| EPI_ISL_413016 | Brazil | 2020-02-28 | Italy | 1 |
| EPI_ISL_413213 | Australia | 2020-02-29 | Iran | 6 |
| EPI_ISL_413221 | UK | 2020-03-02 | Italy | 7 |
| EPI_ISL_413490 | New Zealand | 2020-02-27 | Iran | 1 |
| EPI_ISL_413522 | India | 2020-01-27 | China | NA |
| EPI_ISL_413523 | India | 2020-01-27 | China | 10 |
| EPI_ISL_413550 | Nigeria | 2020-02-27 | Italy | 2 |
| EPI_ISL_413594 | Australia | 2020-02-28 | Singapore | 1* |
| EPI_ISL_413596 | Australia | 2020-02-28 | SE Asia | NA |
| EPI_ISL_413597 | Australia | 2020-03-02 | Iran | 3 |
| EPI_ISL_413598 | Australia | 2020-03-04 | Iran | 3 |
| EPI_ISL_413599 | Australia | 2020-03-04 | Iran | NA |

* when the collection date was the same as the travel date, a minimum of 1 day prior to the collection date was used as the date of travel.

(2) the travel location and date was used for the person having infected the sampled person. For EPI_ISL_406862, x was set to 5 days.

**Table S2.** Acknowledgements table for the 284 genomes used in this study

| Accession ID | Virus name | Location | Collection date | Originating lab | Submitting lab | Authors |
|---|---|---|---|---|---|---|
| **EPI_ISL_402119** | hCoV-19/Wuhan/IVDC-HB-01/2019 | Asia / China / Hubei / Wuhan | 2019-12-30 | National Institute for Viral Disease Control and Prevention, China CDC | National Institute for Viral Disease Control and Prevention, China CDC | Wenjie Tan, Xiang Zhao, Wenling Wang, Xuejun Ma, Yongzhong Jiang, Roujian Lu, Ji Wang, Weimin Zhou, Peihua Niu, Peipei Liu, Faxian Zhan, Weifeng Shi, Baoying Huang, Jun Liu, Li Zhao, Yao Meng, Xiaozhou He, Fei Ye, Na Zhu, Yang Li, Jing Chen, Wenbo Xu, George F. Gao, Guizhen Wu |
| **EPI_ISL_402120** | hCoV-19/Wuhan/IVDC-HB-04/2020 | Asia / China / Hubei / Wuhan | 2020-01-01 | National Institute for Viral Disease Control and Prevention, China CDC | National Institute for Viral Disease Control and Prevention, China CDC | Wenjie Tan, Xiang Zhao, Wenling Wang, Xuejun Ma, Yongzhong Jiang, Roujian Lu, Ji Wang, Weimin Zhou, Peihua Niu, Peipei Liu , Faxian Zhan, Weifeng Shi, Baoying Huang, Jun Liu, Li Zhao, Yao Meng, Xiaozhou He, Fei Ye, Na Zhu, Yang Li, Jing Chen, Wenbo Xu, George F. Gao, Guizhen Wu |
| **EPI_ISL_402121** | hCoV-19/Wuhan/IVDC-HB-05/2019 | Asia / China / Hubei / Wuhan | 2019-12-30 | National Institute for Viral Disease Control and Prevention, China CDC | National Institute for Viral Disease Control and Prevention, China CDC | Wenjie Tan, Xuejun Ma, Xiang Zhao, Wenling Wang, Yongzhong Jiang, Roujian Lu, Ji Wang , Peihua Niu, Weimin Zhou, Faxian Zhan, Weifeng Shi, Baoying Huang, Jun Liu, Li Zhao , Yao Meng, Fei Ye, Na Zhu, Xiaozhou He, Peipei Liu, Yang Li, Jing Chen, Wenbo Xu, George F. Gao, Guizhen Wu |
| **EPI_ISL_402123** | hCoV-19/Wuhan/IPBCAMS-WH-01/2019 | Asia / China / Hubei / Wuhan | 2019-12-24 | Institute of Pathogen Biology, Chinese Academy of Medical Sciences & Peking Union Medical College | Institute of Pathogen Biology, Chinese Academy of Medical Sciences & Peking Union Medical College | Lili Ren, Jianwei Wang, Qi Jin, Zichun Xiang, Zhiqiang Wu, Chao Wu, Yiwei Liu |
| **EPI_ISL_402124** | hCoV-19/Wuhan/WIV04/2019 | Asia / China / Hubei / Wuhan | 2019-12-30 | Wuhan Jinyintan Hospital | Wuhan Institute of Virology, Chinese Academy of Sciences | Peng Zhou, Xing-Lou Yang, Ding-Yu Zhang, Lei Zhang, Yan Zhu, Hao-Rui Si, Zhengli Shi |
| **EPI_ISL_402125** | hCoV-19/Wuhan-Hu-1/2019 | Asia / China | 2019-12-31 | unknown | National Institute for Communicable Disease Control and Prevention (ICDC) Chinese Center for Disease Control and Prevention (China CDC) | Zhang,Y.-Z., Wu,F., Chen,Y.-M., Pei,Y.-Y., Xu,L., Wang,W., Zhao,S., Yu,B., Hu,Y., Tao,Z.-W., Song,Z.-G., Tian,J.-H., Zhang,Y.-L., Liu,Y., Zheng,J.-J., Dai,F.-H., Wang,Q.-M., She,J.-L. and Zhu,T.-Y. |
| **EPI_ISL_402127** | hCoV-19/Wuhan/WIV02/2019 | Asia / China / Hubei / Wuhan | 2019-12-30 | Wuhan Jinyintan Hospital | Wuhan Institute of Virology, Chinese Academy of Sciences | Peng Zhou, Xing-Lou Yang, Ding-Yu Zhang, Lei Zhang, Yan Zhu, Hao-Rui Si, Zhengli Shi |
| **EPI_ISL_402128** | hCoV-19/Wuhan/WIV05/2019 | Asia / China / Hubei / Wuhan | 2019-12-30 | Wuhan Jinyintan Hospital | Wuhan Institute of Virology, Chinese Academy of Sciences | Peng Zhou, Xing-Lou Yang, Ding-Yu Zhang, Lei Zhang, Yan Zhu, Hao-Rui Si, Zhengli Shi |
| **EPI_ISL_402129** | hCoV-19/Wuhan/WIV06/2019 | Asia / China / Hubei / Wuhan | 2019-12-30 | Wuhan Jinyintan Hospital | Wuhan Institute of Virology, Chinese Academy of Sciences | Peng Zhou, Xing-Lou Yang, Ding-Yu Zhang, Lei Zhang, Yan Zhu, Hao-Rui Si, Zhengli Shi |
| **EPI_ISL_402130** | hCoV-19/Wuhan/WIV07/2019 | Asia / China / Hubei / Wuhan | 2019-12-30 | Wuhan Jinyintan Hospital | Wuhan Institute of Virology, Chinese Academy of Sciences | Peng Zhou, Xing-Lou Yang, Ding-Yu Zhang, Lei Zhang, Yan Zhu, Hao-Rui Si, Zhengli Shi |
| **EPI_ISL_402132** | hCoV-19/Wuhan/HBCDC-HB-01/2019 | Asia / China / Hubei / Wuhan | 2019-12-30 | Wuhan Jinyintan Hospital | Hubei Provincial Center for Disease Control and Prevention | Bin Fang, Xiang Li, Xiao Yu, Linlin Liu, Bo Yang, Faxian Zhan, Guojun Ye, Xixiang Huo, Junqiang Xu, Bo Yu, Kun Cai, Jing Li, Yongzhong Jiang. |
| **EPI_ISL_403928** | hCoV-19/Wuhan/IPBCAMS-WH-05/2020 | Asia / China / Hubei / Wuhan | 2020-01-01 | Institute of Pathogen Biology, Chinese Academy | Institute of Pathogen Biology, Chinese Academy of Medical | Lili Ren, Jianwei Wang, Qi Jin, Zichun Xiang, Zhiqiang Wu, Chao Wu, Yiwei Liu |

| | | | | | | |
|---|---|---|---|---|---|---|
| | | | | of Medical Sciences & Peking Union Medical College | Sciences & Peking Union Medical College | |
| **EPI_ISL_ 403929** | hCoV-19/Wuhan/IPBCAMS-WH-04/2019 | Asia / China / Hubei / Wuhan | 2019-12-30 | Institute of Pathogen Biology, Chinese Academy of Medical Sciences & Peking Union Medical College | Institute of Pathogen Biology, Chinese Academy of Medical Sciences & Peking Union Medical College | Lili Ren, Jianwei Wang, Qi Jin, Zichun Xiang, Zhiqiang Wu, Chao Wu, Yiwei Liu |
| **EPI_ISL_ 403930** | hCoV-19/Wuhan/IPBCAMS-WH-03/2019 | Asia / China / Hubei / Wuhan | 2019-12-30 | Institute of Pathogen Biology, Chinese Academy of Medical Sciences & Peking Union Medical College | Institute of Pathogen Biology, Chinese Academy of Medical Sciences & Peking Union Medical College | Lili Ren, Jianwei Wang, Qi Jin, Zichun Xiang, Zhiqiang Wu, Chao Wu, Yiwei Liu |
| **EPI_ISL_ 403931** | hCoV-19/Wuhan/IPBCAMS-WH-02/2019 | Asia / China / Hubei / Wuhan | 2019-12-30 | Institute of Pathogen Biology, Chinese Academy of Medical Sciences & Peking Union Medical College | Institute of Pathogen Biology, Chinese Academy of Medical Sciences & Peking Union Medical College | Lili Ren, Jianwei Wang, Qi Jin, Zichun Xiang, Zhiqiang Wu, Chao Wu, Yiwei Liu |
| **EPI_ISL_ 403932** | hCoV-19/Guangdong/20SF012/2020 | Asia / China / Guangdong / Shenzhen | 2020-01-14 | Guangdong Provincial Center for Diseases Control and Prevention; Guangdong Provincial Public Health | Department of Microbiology, Guangdong Provincial Center for Diseases Control and Prevention | Min Kang, Jie Wu, Jing Lu, Tao Liu, Baisheng Li, Shujiang Mei, Feng Ruan, Lifeng Lin, Changwen Ke, Haojie Zhong, Yingtao Zhang, Lirong Zou, Xuguang Chen, Qi Zhu, Jianpeng Xiao, Jianxiang Geng, Zhe Liu, Jianxiong Hu, Weilin Zeng, Xing Li, Yuhuang Liao, Xiujuan Tang, Songjian Xiao, Ying Wang, Yingchao Song, Xue Zhuang, Lijun Liang, Guanhao He, Huihong Deng, Tie Song, Jianfeng He, Wenjun Ma |
| **EPI_ISL_ 403933** | hCoV-19/Guangdong/20SF013/2020 | Asia / China / Guangdong / Shenzhen | 2020-01-15 | Guangdong Provincial Center for Diseases Control and Prevention; Guangdong Provincial Public Health | Department of Microbiology, Guangdong Provincial Center for Diseases Control and Prevention | Min Kang, Jie Wu, Jing Lu, Tao Liu, Baisheng Li, Shujiang Mei, Feng Ruan, Lifeng Lin, Changwen Ke, Haojie Zhong, Yingtao Zhang, Lirong Zou, Xuguang Chen, Qi Zhu, Jianpeng Xiao, Jianxiang Geng, Zhe Liu, Jianxiong Hu, Weilin Zeng, Xing Li, Yuhuang Liao, Xiujuan Tang, Songjian Xiao, Ying Wang, Yingchao Song, Xue Zhuang, Lijun Liang, Guanhao He, Huihong Deng, Tie Song, Jianfeng He, Wenjun Ma |
| **EPI_ISL_ 403934** | hCoV-19/Guangdong/20SF014/2020 | Asia / China / Guangdong / Shenzhen | 2020-01-15 | Guangdong Provincial Center for Diseases Control and Prevention; Guangdong Provincial Public Health | Department of Microbiology, Guangdong Provincial Center for Diseases Control and Prevention | Min Kang, Jie Wu, Jing Lu, Tao Liu, Baisheng Li, Shujiang Mei, Feng Ruan, Lifeng Lin, Changwen Ke, Haojie Zhong, Yingtao Zhang, Lirong Zou, Xuguang Chen, Qi Zhu, Jianpeng Xiao, Jianxiang Geng, Zhe Liu, Jianxiong Hu, Weilin Zeng, Xing Li, Yuhuang Liao, Xiujuan Tang, Songjian Xiao, Ying Wang, Yingchao Song, Xue Zhuang, Lijun Liang, Guanhao He, Huihong Deng, Tie Song, Jianfeng He, Wenjun Ma |
| **EPI_ISL_ 403935** | hCoV-19/Guangdong/20SF025/2020 | Asia / China / Guangdong / Shenzhen | 2020-01-15 | Guangdong Provincial Center for Diseases Control and Prevention; Guangdong Provincial Public Health | Department of Microbiology, Guangdong Provincial Center for Diseases Control and Prevention | Min Kang, Jie Wu, Jing Lu, Tao Liu, Baisheng Li, Shujiang Mei, Feng Ruan, Lifeng Lin, Changwen Ke, Haojie Zhong, Yingtao Zhang, Lirong Zou, Xuguang Chen, Qi Zhu, Jianpeng Xiao, Jianxiang Geng, Zhe Liu, Jianxiong Hu, Weilin Zeng, Xing Li, Yuhuang Liao, Xiujuan Tang, Songjian Xiao, Ying Wang, Yingchao Song, Xue Zhuang, Lijun Liang, Guanhao He, Huihong Deng, Tie Song, Jianfeng He, Wenjun Ma |
| **EPI_ISL_ 403936** | hCoV-19/Guangdong/20SF028/2020 | Asia / China / Guangdong / Zhuhai | 2020-01-17 | Guangdong Provincial Center for Diseases Control and | Department of Microbiology, Guangdong Provincial Center for Diseases | Min Kang, Jie Wu, Jing Lu, Tao Liu, Baisheng Li, Shujiang Mei, Feng Ruan, Lifeng Lin, Changwen Ke, Haojie Zhong, Yingtao Zhang, Lirong Zou, Xuguang Chen, Qi Zhu, Jianpeng Xiao, Jianxiang |

| | | | | | | |
|---|---|---|---|---|---|---|
| | | | | Prevention; Guangdong Provincial Public Health | Control and Prevention | Geng, Zhe Liu, Jianxiong Hu, Weilin Zeng, Xing Li, Yuhuang Liao, Xiujuan Tang, Songjian Xiao, Ying Wang, Yingchao Song, Xue Zhuang, Lijun Liang, Guanhao He, Huihong Deng, Tie Song, Jianfeng He, Wenjun Ma |
| **EPI_ISL_403937** | hCoV-19/Guangdong/20SF040/2020 | Asia / China / Guangdong / Zhuhai | 2020-01-18 | Guangdong Provincial Center for Diseases Control and Prevention; Guangdong Provincial Public Health | Department of Microbiology, Guangdong Provincial Center for Diseases Control and Prevention | Min Kang, Jie Wu, Jing Lu, Tao Liu, Baisheng Li, Shujiang Mei, Feng Ruan, Lifeng Lin, Changwen Ke, Haojie Zhong, Yingtao Zhang, Lirong Zou, Xuguang Chen, Qi Zhu, Jianpeng Xiao, Jianxiang Geng, Zhe Liu, Jianxiong Hu, Weilin Zeng, Xing Li, Yuhuang Liao, Xiujuan Tang, Songjian Xiao, Ying Wang, Yingchao Song, Xue Zhuang, Lijun Liang, Guanhao He, Huihong Deng, Tie Song, Jianfeng He, Wenjun Ma |
| **EPI_ISL_403962** | hCoV-19/Thailand/61/2020 | Asia / Thailand / Nonthaburi | 2020-01-08 | Bamrasnaradura Hospital | 1. Department of Medical Sciences, Ministry of Public Health, Thailand 2. Thai Red Cross Emerging Infectious Diseases - Health Science Centre 3. Department of Disease Control, Ministry of Public Health, Thailand | Pilailuk,Okada; Siripaporn,Phuygun; Thanutsapa,Thanadachakul; Supaporn,Wacharapluesadee; Sittiporn,Parnmen; Warawan,Wongboot; Sunthareeya,Waicharoen; Rome,Buathong; Malinee,Chittaganpitch; Nanthawan,Mekha |
| **EPI_ISL_403963** | hCoV-19/Thailand/74/2020 | Asia / Thailand / Nonthaburi | 2020-01-13 | Bamrasnaradura Hospital | 1. Department of Medical Sciences, Ministry of Public Health, Thailand 2. Thai Red Cross Emerging Infectious Diseases - Health Science Centre 3. Department of Disease Control, Ministry of Public Health, Thailand | Pilailuk,Okada; Siripaporn,Phuygun; Thanutsapa,Thanadachakul; Supaporn,Wacharapluesadee; Sittiporn,Parnmen; Warawan,Wongboot; Sunthareeya,Waicharoen; Rome,Buathong; Malinee,Chittaganpitch; Nanthawan,Mekha |
| **EPI_ISL_404227** | hCoV-19/Zhejiang/WZ-01/2020 | Asia / China / Zhejiang | 2020-01-16 | Zhejiang Provincial Center for Disease Control and Prevention | Department of Microbiology, Zhejiang Provincial Center for Disease Control and Prevention | Yin Chen, Yanjun Zhang, Haiyan Mao, Junhang Pan, Xiuyu Lou, Yiyu Lu, Juying Yan, Hanping Zhu, Jian Gao, Yan Feng, Yi Sun, Hao Yan, Zhen Li, Yisheng Sun, Liming Gong, Qiong Ge, Wen Shi, Xinying Wang, Wenwu Yao, Zhangnv Yang, Fang Xu, Chen Chen, Enfu Chen, Zhen Wang, Zhiping Chen, Jianmin Jiang, Chonggao Hu |
| **EPI_ISL_404228** | hCoV-19/Zhejiang/WZ-02/2020 | Asia / China / Zhejiang | 2020-01-17 | Zhejiang Provincial Center for Disease Control and Prevention | Department of Microbiology, Zhejiang Provincial Center for Disease Control and Prevention | Yanjun Zhang, Yin Chen, Haiyan Mao, Junhang Pan, Xiuyu Lou, Yiyu Lu, Juying Yan, Hanping Zhu, Jian Gao, Yan Feng, Yi Sun, Hao Yan, Zhen Li, Yisheng Sun, Liming Gong, Qiong Ge, Wen Shi, Xinying Wang, Wenwu Yao, Zhangnv Yang, Fang Xu, Chen Chen, Enfu Chen, Zhen Wang, Zhiping Chen, Jianmin Jiang, Chonggao Hu |
| **EPI_ISL_404253** | hCoV-19/USA/IL1/2020 | North America / USA / Illinois / Chicago | 2020-01-21 | IL Department of Public Health Chicago Laboratory | Pathogen Discovery, Respiratory Viruses Branch, Division of Viral Diseases, Centers for Dieases Control and Prevention | Ying Tao, Krista Queen, Clinton R. Paden, Jing Zhang, Yan Li, Anna Uehara, Xiaoyan Lu, Brian Lynch, Senthil Kumar K. Sakthivel, Brett L. Whitaker, Shifaq Kamili, Lijuan Wang, Janna' R. Murray, Susan I. Gerber, Stephen Lindstrom, Suxiang Tong |
| **EPI_ISL_404895** | hCoV-19/USA/WA1/2020 | North America / USA / Washington / Snohomish County | 2020-01-19 | Providence Regional Medical Center | Division of Viral Diseases, Centers for Disease Control and Prevention | Queen,K., Tao,Y., Li,Y., Paden,C.R., Lu,X., Zhang,J., Gerber,S.I., Lindstrom,S., Tong,S. |

| | | | | | | |
|---|---|---|---|---|---|---|
| **EPI_ISL_405839** | hCoV-19/Shenzhen/HKU-SZ-005/2020 | Asia / China / Guangdong / Shenzhen | 2020-01-11 | The University of Hong Kong - Shenzhen Hospital | Li Ka Shing Faculty of Medicine, The University of Hong Kong | Chan,J.F.-W., Yuan,S., Kok,K.H., To,K.K.-W., Chu,H., Yang,J., Xing,F., Liu,J., Yip,C.C.-Y., Poon,R.W.-S., Tsai,H.W., Lo,S.K.-F., Chan,K.H., Poon,V.K.-M., Chan,W.M., Ip,J.D., Cai,J.P., Cheng,V.C.-C., Chen,H., Hui,C.K.-M. and Yuen,K.Y. |
| **EPI_ISL_406031** | hCoV-19/Taiwan/2/2020 | Asia / Taiwan / Kaohsiung | 2020-01-23 | Centers for Disease Control, R.O.C. (Taiwan) | Centers for Disease Control, R.O.C. (Taiwan) | Ji-Rong Yang, Yu-Chi Lin, Jung-Jung Mu, Ming-Tsan Liu, Shu-Ying Li |
| **EPI_ISL_406034** | hCoV-19/USA/CA1/2020 | North America / USA / California / Los Angeles | 2020-01-23 | California Department of Public Health | Pathogen Discovery, Respiratory Viruses Branch, Division of Viral Diseases, Centers for Dieases Control and Prevention | Anna Uehara, Krista Queen, Ying Tao, Yan Li, Clinton R. Paden, Jing Zhang, Xiaoyan Lu, Brian Lynch, Senthil Kumar K. Sakthivel, Brett L. Whitaker, Shifaq Kamili, Lijuan Wang, Janna' R. Murray, Susan I. Gerber, Stephen Lindstrom, Suxiang Tong |
| **EPI_ISL_406036** | hCoV-19/USA/CA2/2020 | North America / USA / California / Orange County | 2020-01-22 | California Department of Public Health | Pathogen Discovery, Respiratory Viruses Branch, Division of Viral Diseases, Centers for Dieases Control and Prevention | Anna Uehara, Krista Queen, Ying Tao, Yan Li, Clinton R. Paden, Jing Zhang, Xiaoyan Lu, Brian Lynch, Senthil Kumar K. Sakthivel, Brett L. Whitaker, Shifaq Kamili, Lijuan Wang, Janna' R. Murray, Susan I. Gerber, Stephen Lindstrom, Suxiang Tong |
| **EPI_ISL_406223** | hCoV-19/USA/AZ1/2020 | North America / USA / Arizona / Phoenix | 2020-01-22 | Arizona Department of Health Services | Pathogen Discovery, Respiratory Viruses Branch, Division of Viral Diseases, Centers for Disease Control and Prevention | Ying Tao, Clinton R. Paden, Krista Queen, Anna Uehara, Yan Li, Jing Zhang, Xiaoyan Lu, Brian Lynch, Senthil Kumar K. Sakthivel, Brett L. Whitaker, Shifaq Kamili, Lijuan Wang, Janna' R. Murray, Susan I. Gerber, Stephen Lindstrom, Suxiang Tong |
| **EPI_ISL_406531** | hCoV-19/Guangdong/20SF174/2020 | Asia / China / Guangdong / Zhuhai | 2020-01-22 | Guangdong Provincial Center for Diseases Contorl and Prevention; Guangdong Provincial Public Health | Guangdong Provincial Center for Disease Control and Prevention | Min Kang, Jie Wu, Jing Lu, Tao Liu, Baisheng Li, Shujiang Mei, Feng Ruan, Lifeng Lin, Changwen Ke, Haojie Zhong, Yingtao Zhang, Lirong Zou, Xuguang Chen, Qi Zhu, Jianpeng Xiao, Jianxiang Geng, Zhe Liu, Jianxiong Hu, Weilin Zeng, Xing Li, Yuhuang Liao, Xiujuan Tang, Songjian Xiao, Ying Wang, Yingchao Song, Xue Zhuang, Lijun Liang, Guanhao He, Huihong Deng, Tie Song, Jianfeng He, Wenjun Ma |
| **EPI_ISL_406533** | hCoV-19/Guangzhou/20SF206/2020 | Asia / China / Guangdong / Guangzhou | 2020-01-22 | Guangdong Provincial Center for Diseases Contorl and Prevention; Guangdong Provincial Public Health | Guangdong Provincial Center for Diseases Control and Prevention | Min Kang, Jie Wu, Jing Lu, Tao Liu, Baisheng Li, Shujiang Mei, Feng Ruan, Lifeng Lin, Changwen Ke, Haojie Zhong, Yingtao Zhang, Lirong Zou, Xuguang Chen, Qi Zhu, Jianpeng Xiao, Jianxiang Geng, Zhe Liu, Jianxiong Hu, Weilin Zeng, Xing Li, Yuhuang Liao, Xiujuan Tang, Songjian Xiao, Ying Wang, Yingchao Song, Xue Zhuang, Lijun Liang, Guanhao He, Huihong Deng, Tie Song, Jianfeng He, Wenjun Ma |
| **EPI_ISL_406534** | hCoV-19/Foshan/20SF207/2020 | Asia / China / Guangdong / Foshan | 2020-01-22 | Guangdong Provincial Center for Diseases Control and Prevention; Guangdong Provincial Public Health | Guangdong Provincial Center for Diseases Control and Prevention | Min Kang, Jie Wu, Jing Lu, Tao Liu, Baisheng Li, Shujiang Mei, Feng Ruan, Lifeng Lin, Changwen Ke, Haojie Zhong, Yingtao Zhang, Lirong Zou, Xuguang Chen, Qi Zhu, Jianpeng Xiao, Jianxiang Geng, Zhe Liu, Jianxiong Hu, Weilin Zeng, Xing Li, Yuhuang Liao, Xiujuan Tang, Songjian Xiao, Ying Wang, Yingchao Song, Xue Zhuang, Lijun Liang, Guanhao He, Huihong Deng, Tie Song, Jianfeng He, Wenjun Ma |
| **EPI_ISL_406535** | hCoV-19/Foshan/20SF210/2020 | Asia / China / Guangdong / Foshan | 2020-01-22 | Guangdong Provincial Center for Diseases Control and Prevention; Guangdong Provincial Public Health | Guangdong Provincial Center for Diseases Control and Prevention | Min Kang, Jie Wu, Jing Lu, Tao Liu, Baisheng Li, Shujiang Mei, Feng Ruan, Lifeng Lin, Changwen Ke, Haojie Zhong, Yingtao Zhang, Lirong Zou, Xuguang Chen, Qi Zhu, Jianpeng Xiao, Jianxiang Geng, Zhe Liu, Jianxiong Hu, Weilin Zeng, Xing Li, Yuhuang Liao, Xiujuan Tang, Songjian Xiao, Ying Wang, Yingchao Song, Xue Zhuang, Lijun Liang, Guanhao He, Huihong Deng, Tie Song, Jianfeng He, Wenjun Ma |

| | | | | | | |
|---|---|---|---|---|---|---|
| **EPI_ISL_406536** | hCoV-19/Foshan/20SF211/2020 | Asia / China / Guangdong / Foshan | 2020-01-22 | Guangdong Provincial Center for Diseases Control and Prevention; Guangdong Provincial Public Health | Guangdong Provincial Center for Diseases Control and Prevention | Min Kang, Jie Wu, Jing Lu, Tao Liu, Baisheng Li, Shujiang Mei, Feng Ruan, Lifeng Lin, Changwen Ke, Haojie Zhong, Yingtao Zhang, Lirong Zou, Xuguang Chen, Qi Zhu, Jianpeng Xiao, Jianxiang Geng, Zhe Liu, Jianxiong Hu, Weilin Zeng, Xing Li, Yuhuang Liao, Xiujuan Tang, Songjian Xiao, Ying Wang, Yingchao Song, Xue Zhuang, Lijun Liang, Guanhao He, Huihong Deng, Tie Song, Jianfeng He, Wenjun Ma |
| **EPI_ISL_406538** | hCoV-19/Guangdong/20SF201/2020 | Asia / China / Guangdong | 2020-01-23 | Guangdong Provincial Center for Diseases Control and Prevention;Guangdong Provincial Institute of Public Health | Guangdong Provincial Center for Diseases Control and Prevention | Min Kang, Jie Wu, Jing Lu, Tao Liu, Baisheng Li, Shujiang Mei, Feng Ruan, Lifeng Lin, Changwen Ke, Haojie Zhong, Yingtao Zhang, Lirong Zou, Xuguang Chen, Qi Zhu, Jianpeng Xiao, Jianxiang Geng, Zhe Liu, Jianxiong Hu, Weilin Zeng, Xing Li, Yuhuang Liao, Xiujuan Tang, Songjian Xiao, Ying Wang, Yingchao Song, Xue Zhuang, Lijun Liang, Guanhao He, Huihong Deng, Tie Song, Jianfeng He, Wenjun Ma |
| **EPI_ISL_406593** | hCoV-19/Shenzhen/SZTH-002/2020 | Asia / China / Guangdong / Shenzhen | 2020-01-13 | Shenzhen Key Laboratory of Pathogen and Immunity, National Clinical Research Center for Infectious Disease, Shenzhen Third People's Hospital | Shenzhen Key Laboratory of Pathogen and Immunity, National Clinical Research Center for Infectious Disease, Shenzhen Third People's Hospital | Yang Yang, Chenguang Shen, Li Xing, Zhixiang Xu, Haixia Zheng, Yingxia Liu |
| **EPI_ISL_406594** | hCoV-19/Shenzhen/SZTH-003/2020 | Asia / China / Guangdong / Shenzhen | 2020-01-16 | Shenzhen Key Laboratory of Pathogen and Immunity, National Clinical Research Center for Infectious Disease, Shenzhen Third People's Hospital | Shenzhen Key Laboratory of Pathogen and Immunity, National Clinical Research Center for Infectious Disease, Shenzhen Third People's Hospital | Yang Yang, Chenguang Shen, Li Xing, Zhixiang Xu, Haixia Zheng, Yingxia Liu |
| **EPI_ISL_406596** | hCoV-19/France/IDF0372/2020 | Europe / France / Ile-de-France / Paris | 2020-01-23 | Department of Infectious and Tropical Diseases, Bichat Claude Bernard Hospital, Paris | National Reference Center for Viruses of Respiratory Infections, Institut Pasteur, Paris | Mélanie Albert, Marion Barbet, Sylvie Behillil, Méline Bizard, Angela Brisebarre, Flora Donati, Vincent Enouf, Maud Vanpeene, Sylvie van der Werf, Yazdan Yazdanpanah, Xavier Lescure. |
| **EPI_ISL_406597** | hCoV-19/France/IDF0373/2020 | Europe / France / Ile-de-France / Paris | 2020-01-23 | Department of Infectious and Tropical Diseases, Bichat Claude Bernard Hospital, Paris | National Reference Center for Viruses of Respiratory Infections, Institut Pasteur, Paris | Mélanie Albert, Marion Barbet, Sylvie Behillil, Méline Bizard, Angela Brisebarre, Flora Donati, Vincent Enouf, Maud Vanpeene, Sylvie van der Werf, Yazdan Yazdanpanah, Xavier Lescure. |
| **EPI_ISL_406716** | hCoV-19/Wuhan/WHU01/2020 | Asia / China / Hubei / Wuhan | 2020-01-02 | unknown | State Key Laboratory of Virology, Wuhan University | Chen,L., Liu,W., Zhang,Q., Xu,K., Ye,G., Wu,W., Sun,Z., Liu,F., Wu,K., Mei,Y., Zhang,W., Chen,Y., Li,Y., Shi,M., Lan,K. and Liu,Y. |
| **EPI_ISL_406717** | hCoV-19/Wuhan/WHU02/2020 | Asia / China / Hubei / Wuhan | 2020-01-02 | unknown | State Key Laboratory of Virology, Wuhan University | Chen,L., Liu,W., Zhang,Q., Xu,K., Ye,G., Wu,W., Sun,Z., Liu,F., Wu,K., Mei,Y., Zhang,W., Chen,Y., Li,Y., Shi,M., Lan,K. and Liu,Y. |
| **EPI_ISL_406798** | hCoV-19/Wuhan/WH01/2019 | Asia / China / Hubei / Wuhan | 2019-12-26 | General Hospital of Central Theater Command of People's Liberation Army of China | BGI & Institute of Microbiology, Chinese Academy of Sciences & Shandong First Medical University & Shandong Academy of Medical Sciences & General Hospital of Central Theater | Weijun Chen, Yuhai Bi, Weifeng Shi and Zhenhong Hu |

| | | | | | | |
|---|---|---|---|---|---|---|
| | | | | | Command of People's Liberation Army of China | |
| **EPI_ISL_406800** | hCoV-19/Wuhan/WH03/2020 | Asia / China / Hubei / Wuhan | 2020-01-01 | General Hospital of Central Theater Command of People's Liberation Army of China | BGI & Institute of Microbiology, Chinese Academy of Sciences & Shandong First Medical University & Shandong Academy of Medical Sciences & General Hospital of Central Theater Command of People's Liberation Army of China | Weijun Chen, Yuhai Bi, Weifeng Shi and Zhenhong Hu |
| **EPI_ISL_406801** | hCoV-19/Wuhan/WH04/2020 | Asia / China / Hubei / Wuhan | 2020-01-05 | General Hospital of Central Theater Command of People's Liberation Army of China | BGI & Institute of Microbiology, Chinese Academy of Sciences & Shandong First Medical University & Shandong Academy of Medical Sciences & General Hospital of Central Theater Command of People's Liberation Army of China | Weijun Chen, Yuhai Bi, Weifeng Shi and Zhenhong Hu |
| **EPI_ISL_406844** | hCoV-19/Australia/VIC01/2020 | Oceania / Australia / Victoria / Clayton | 2020-01-25 | Monash Medical Centre | Collaboration between the University of Melbourne at The Peter Doherty Institute for Infection and Immunity, and the Victorian Infectious Disease Reference Laboratory | Caly,L., Seemann,T., Schultz,M., Druce,J. and Taiaroa,G |
| **EPI_ISL_406862** | hCoV-19/Germany/BavPat1/2020 | Europe / Germany / Bavaria / Munich | 2020-01-28 | Charité Universitätsmedizin Berlin, Institute of Virology; Institut für Mikrobiologie der Bundeswehr, Munich | Charité Universitätsmedizin Berlin, Institute of Virology | Victor M Corman, Julia Schneider, Talitha Veith, Barbara Mühlemann, Markus Antwerpen, Christian Drosten, Roman Wölfel |
| **EPI_ISL_406970** | hCoV-19/Hangzhou/HZ-1/2020 | Asia / China / Zhejiang / Hangzhou | 2020-01-20 | Hangzhou Center for Disease and Control Microbiology Lab | Hangzhou Center for Disease and Control Microbiology Lab | Yu Hua, Wang Haoqiu, Li Jun, Yu Xinfeng |
| **EPI_ISL_406973** | hCoV-19/Singapore/1/2020 | Asia / Singapore | 2020-01-23 | Singapore General Hospital | National Public Health Laboratory | Mak, TM; Octavia S; Chavatte JM; Zhou, ZY; Cui, L; Lin, RTP |
| **EPI_ISL_407071** | hCoV-19/England/01/2020 | Europe / United Kingdom / England | 2020-01-29 | Respiratory Virus Unit, Microbiology Services Colindale, Public Health England | Respiratory Virus Unit, Microbiology Services Colindale, Public Health England | Monica Galiano, Shahjahan Miah, Richard Myers, Angie Lackenby, Omolola Akinbami, Tiina Talts, Leena Bhaw, Kirstin Edwards, Jonathan Hubb, Joanna Ellis, Maria Zambon |
| **EPI_ISL_407073** | hCoV-19/England/02/2020 | Europe / United Kingdom / England | 2020-01-29 | Respiratory Virus Unit, Microbiology Services Colindale, Public Health England | Respiratory Virus Unit, Microbiology Services Colindale, Public Health England | Monica Galiano, Shahjahan Miah, Richard Myers, Angie Lackenby, Omolola Akinbami, Tiina Talts, Leena Bhaw, Kirstin Edwards, Jonathan Hubb, Joanna Ellis, Maria Zambon. |

| | | | | | | |
|---|---|---|---|---|---|---|
| **EPI_ISL_407079** | hCoV-19/Finland/1/2020 | Europe / Finland / Lapland | 2020-01-29 | Lapland Central Hospital | Department of Virology, University of Helsinki and Helsinki University Hospital, Helsinki, Finland | Teemu Smura, Suvi Kuivanen, Hannimari Kallio-Kokko, Olli Vapalahti |
| **EPI_ISL_407084** | hCoV-19/Japan/AI-I-004/2020 | Asia / Japan / Aichi | 2020-01-25 | Department of Virology III, National Institute of Infectious Diseases | Pathogen Genomics Center, National Institute of Infectious Diseases | Tsuyoshi Sekizuka, Shutoku Matsuyama, Naganori Nao, Kazuya Shirato, Shinji Watanabe, Makoto Takeda, Makoto Kuroda |
| **EPI_ISL_407193** | hCoV-19/South Korea/KCDC03/2020 | Asia / South Korea / Gyeonggi-do | 2020-01-25 | Korea Centers for Disease Control & Prevention (KCDC) Center for Laboratory Control of Infectious Diseases Division of Viral Diseases | Korea Centers for Disease Control & Prevention (KCDC) Center for Laboratory Control of Infectious Diseases Division of Viral Diseases | Jeong-Min Kim, Yoon-Seok Chung, Namjoo Lee, Mi-Seon Kim, SangHee Woo, Hye-Joon Jo, Sehee Park, Heui Man Kim, Myung Guk Han |
| **EPI_ISL_407313** | hCoV-19/Hangzhou/HZCDC0001/2020 | Asia / China / Zhejiang / Hangzhou | 2020-01-19 | Hangzhou Center for Disease Control and Prevention | Hangzhou Center for Disease Control and Prevention | Jun Li, Haoqiu Wang, Hua Yu, Lingfeng Mao, Xinfen Yu, Zhou Sun, Qingxin Kong, Xin Qian, Shuchang Chen, Xuchu Wang |
| **EPI_ISL_407893** | hCoV-19/Australia/NSW01/2020 | Oceania / Australia / New South Wales / Sydney | 2020-01-24 | Centre for Infectious Diseases and Microbiology Laboratory Services | NSW Health Pathology - Institute of Clinical Pathology and Medical Research; Westmead Hospital; University of Sydney | Eden J-S, Carter I, Rahman H, Holmes EC, Rockett R, O’Sullivan MV, Sintchenko V, Chen SC, Maddocks S, Kok J and Dwyer DE for the 2019-nCoV Study Group |
| **EPI_ISL_407894** | hCoV-19/Australia/QLD01/2020 | Oceania / Australia / Queensland / Gold Coast | 2020-01-28 | Pathology Queensland | Public Health Virology Laboratory | Ben Huang, Alyssa Pyke, Amanda De Jong, Andrew Van Den Hurk, Carmel Taylor, David Warrilow, Doris Genge, Elisabeth Gamez, Glen Hewitson, Ian Maxwell Mackay, Inga Sultana, Jamie McMahon, Jean Barcelon, Judy Northill, Mitchell Finger, Natalie Simpson, Neelima Nair, Peter Burtonclay, Peter Moore, Sarah Wheatley, Sean Moody, Sonja Hall-Mendelin, Timothy Gardam, and Frederick Moore. |
| **EPI_ISL_407896** | hCoV-19/Australia/QLD02/2020 | Oceania / Australia / Queensland / Gold Coast | 2020-01-30 | Pathology Queensland | Public Health Virology Laboratory | Ben Huang, Alyssa Pyke, Amanda De Jong, Andrew Van Den Hurk, Carmel Taylor, David Warrilow, Doris Genge, Elisabeth Gamez, Glen Hewitson, Ian Maxwell Mackay, Inga Sultana, Jamie McMahon, Jean Barcelon, Judy Northill, Mitchell Finger, Natalie Simpson, Neelima Nair, Peter Burtonclay, Peter Moore, Sarah Wheatley, Sean Moody, Sonja Hall-Mendelin, Timothy Gardam, and Frederick Moore. |
| **EPI_ISL_407976** | hCoV-19/Belgium/GHB-03021/2020 | Europe / Belgium / Leuven | 2020-02-03 | KU Leuven, Clinical and Epidemiological Virology | KU Leuven, Clinical and Epidemiological Virology | Bert Vanmechelen, Elke Wollants, Annabel Rector, Els Keyaerts, Lies Laenen, Marc Van Ranst, and Piet Maes |
| **EPI_ISL_407987** | hCoV-19/Singapore/2/2020 | Asia / Singapore | 2020-01-25 | Singapore General Hospital | Programme in Emerging Infectious Diseases, Duke-NUS Medical School | Danielle E Anderson, Martin Linster, Yan Zhuang, Jayanthi Jayakumar, Kian Sing Chan, Lynette LE Oon, Jenny GH Low, Yvonne CF Su, Linfa Wang, Gavin JD Smith |
| **EPI_ISL_407988** | hCoV-19/Singapore/3/2020 | Asia / Singapore | 2020-02-01 | National Centre for Infectious Diseases | Programme in Emerging Infectious Diseases, Duke-NUS Medical School | Danielle E Anderson, Martin Linster, Yan Zhuang, Jayanthi Jayakumar, David CB Lye, Yee Sin Leo, Barnaby E Young, Yvonne CF Su, Linfa Wang, Gavin JD Smith |
| **EPI_ISL_408008** | hCoV-19/USA/CA3/2020 | North America / USA / California | 2020-01-29 | California Department of Health | Pathogen Discovery, Respiratory Viruses Branch, Division of Viral Diseases, Centers | Krista Queen, Jing Zhang, Yan Li, Ying Tao, Anna Uehara, Clinton Paden, Xiaoyan Lu, Brian Lynch, Senthil Kumar K. Sakthivel, Brett L. Whitaker, Shifaq Kamili, Lijuan Wang, Janna' R. Murray, |

| | | | | | | |
|---|---|---|---|---|---|---|
| | | | | | for Disease Control and Prevention | Susan I. Gerber, Stephen Lindstrom, Suxiang Tong |
| **EPI_ISL_ 408009** | hCoV-19/USA/ CA4/2020 | North America / USA / California | 2020-01-29 | California Department of Health | Pathogen Discovery, Respiratory Viruses Branch, Division of Viral Diseases, Centers for Dieases Control and Prevention | Krista Queen, Jing Zhang, Yan Li, Ying Tao, Anna Uehara, Clinton Paden, Xiaoyan Lu, Brian Lynch, Senthil Kumar K. Sakthivel, Brett L. Whitaker, Shifaq Kamili, Lijuan Wang, Janna' R. Murray, Susan I. Gerber, Stephen Lindstrom, Suxiang Tong |
| **EPI_ISL_ 408010** | hCoV-19/USA/ CA5/2020 | North America / USA / California | 2020-01-29 | California Department of Health | Pathogen Discovery, Respiratory Viruses Branch, Division of Viral Diseases, Centers for Dieases Control and Prevention | Ying Tao, Krista Queen, Jing Zhang, Yan Li, Anna Uehara, Clinton Paden, Xiaoyan Lu, Brian Lynch, Senthil Kumar K. Sakthivel, Brett L. Whitaker, Shifaq Kamili, Lijuan Wang, Janna' R. Murray, Susan I. Gerber, Stephen Lindstrom, Suxiang Tong |
| **EPI_ISL_ 408430** | hCoV-19/Franc e/IDF0515/202 0 | Europe / France / Ile-de-Franc e / Paris | 2020-01-29 | Department of Infectious and Tropical Diseases, Bichat Claude Bernard Hospital, Paris | National Reference Center for Viruses of Respiratory Infections, Institut Pasteur, Paris | Mélanie Albert, Marion Barbet, Sylvie Behillil, Méline Bizard, Angela Brisebarre, Flora Donati, Vincent Enouf, Maud Vanpeene, Sylvie van der Werf, Yazdan Yazdanpanah, Xavier Lescure |
| **EPI_ISL_ 408431** | hCoV-19/Franc e/IDF0626/202 0 | Europe / France / Ile-de-Franc e / Paris | 2020-01-29 | Sorbonne Université, Inserm et Assistance Publique-Hôpitau x de Paris (Pitié Salpétrière) | National Reference Center for Viruses of Respiratory Infections, Institut Pasteur, Paris | Mélanie Albert, Marion Barbet, Sylvie Behillil, Méline Bizard, Angela Brisebarre, Flora Donati, Vincent Enouf, Maud Vanpeene, Sylvie van der Werf, Sonia Burrel, Anne-Geneviève Marcelin, Vincent Calvez, David Boutolleau, Elise Klément, Valérie Pourcher, Eric Caumes. |
| **EPI_ISL_ 408478** | hCoV-19/Chon gqing/YC01/20 20 | Asia / China / Chongqinq / Yongchuan | 2020-01-21 | Yongchuan District Center for Disease Control and Prevention | Chongqing Municipal Center for Disease Control and Prevention | Ye Sheng, Tang Yun, Ling Hua,Yu zhen,Chen Shuang,Tan ZhangPing, Su Kun, Li Qing, Tang Wenge, Rong Rong |
| **EPI_ISL_ 408479** | hCoV-19/Chon gqing/ZX01/20 20 | Asia / China / Chongqing / Zhongxian | 2020-01-23 | Zhongxian Center for Disease Control and Prevention | Chongqing Municipal Center for Disease Control and Prevention | Ye Sheng, Tang Yun, Ling Hua, Zhang Hong, Yu zhen,Chen Shuang,Tan ZhangPing, Su Kun, Li Qin, Tang Wenge, Rong Rong |
| **EPI_ISL_ 408480** | hCoV-19/Yunn an/IVDC-YN-00 3/2020 | Asia / China / Yunnan / Kunming | 2020-01-17 | National Institute for Viral Disease Control and Prevention, China CDC | National Institute for Viral Disease Control & Prevention, CCDC | Wenjie Tan, Xiaoqing Fu, Xiang Zhao, Wenling Wang, Peihua Niu, Roujian Lu,Yanhong Sun, Baoying Huang, Li Zhao, Fei Ye, Wenbo Xu, George F. Gao, Guizhen Wu |
| **EPI_ISL_ 408481** | hCoV-19/Chon gqing/IVDC-CQ -001/2020 | Asia / China / Chongqing | 2020-01-18 | National Institute for Viral Disease Control and Prevention, China CDC | National Institute for Viral Disease Control & Prevention, CCDC | Wenjie Tan, Hengqin Wang, Xiang Zhao, Wenling Wang, Peihua Niu, Roujian Lu, Sheng Ye, Baoying Huang, Li Zhao, Fei Ye, Wenbo Xu, George F. Gao, Guizhen Wu |
| **EPI_ISL_ 408482** | hCoV-19/Shan dong/IVDC-SD-001/2020 | Asia / China / Shandong / Qingdao | 2020-01-19 | National Institute for Viral Disease Control and Prevention, China CDC | National Institute for Viral Disease Control & Prevention, CCDC | Wenjie Tan, Zhaoguo Wang, Xiang Zhao, Wenling Wang, Peihua Niu, Roujian Lu, Ti Liu, Baoying Huang, Li Zhao, Fei Ye, Wenbo Xu, George F. Gao, Guizhen Wu |
| **EPI_ISL_ 408484** | hCoV-19/Sichu an/IVDC-SC-00 1/2020 | Asia / China / Sichuan / Chengdu | 2020-01-15 | National Institute for Viral Disease Control and Prevention, China CDC | National Institute for Viral Disease Control & Prevention, CCDC | Wenjie Tan, Jianan Xu, Wenling Wang, Peihua Niu, Roujian Lu, Huiping Yang, Xiang Zhao, Baoying Huang, Li Zhao, Fei Ye, Wenbo Xu, George F. Gao, Guizhen Wu |
| **EPI_ISL_ 408486** | hCoV-19/Jiang xi/IVDC-JX-002 /2020 | Asia / China / Jiangxi / Pingxiang | 2020-01-11 | National Institute for Viral Disease Control and Prevention, China CDC | National Institute for Viral Disease Control & Prevention, CCDC | Wenjie Tan, Yong Shi, Wenling Wang, Peihua Niu, Roujian Lu, Jianxiong Li, Xiang Zhao, Baoying Huang, Li Zhao, Fei Ye, Wenbo Xu, George F. Gao, Guizhen Wu |
| **EPI_ISL_ 408488** | hCoV-19/Jiang su/IVDC-JS-001 /2020 | Asia / China / Jiangsu / Huaian | 2020-01-19 | National Institute for Viral Disease Control and | National Institute for Viral Disease Control & Prevention, CCDC | Wenjie Tan, Shenjiao Wang, Wenling Wang, Peihua Niu, Roujian Lu, Kangchen Zhao, Xiang |

| | | | | | | |
|---|---|---|---|---|---|---|
| | | | | Prevention, China CDC | | Zhao, Baoying Huang, Li Zhao, Fei Ye, Wenbo Xu, George F. Gao, Guizhen Wu |
| **EPI_ISL_ 408489** | hCoV-19/Taiwan/NTU01/2020 | Asia / Taiwan / Taipei | 2020-01-31 | Department of Laboratory Medicine, National Taiwan University Hospital | Microbial Genomics Core Lab, National Taiwan University Centers of Genomic and Precision Medicine | Shiou-Hwei Yeh, You-Yu Lin, Ya-Yun Lai, Chiao-Ling Li, Shan-Chwen Chang, Pei-Jer Chen, Sui-Yuan Chang |
| **EPI_ISL_ 408514** | hCoV-19/env/ Wuhan/IVDC-HBF13-20/2020 | Asia / China / Hubei / Wuhan | 2020-01-01 | Institute of Viral Disease Control and Prevention, China CDC | Institute of Viral Disease Control and Prevention, China CDC | William J. Liu, Peipei Liu, Xiang Zhao, Peihua Niu, Yingze Zhao, Wenwen Lei, Ziqian Xu, Shumei Zou, Wei Zhen, Beiwei Ye, Mengjie Yang, Weifeng Shi, Roujian Lu, Wenjie Tan, Zhixiao Chen, Yuchao Wu, Juan Song, Weimin Zhou, Dayan Wang, Jun Han, Wenbo Xu, George F. Gao, Guizhen Wu |
| **EPI_ISL_ 408515** | hCoV-19/env/ Wuhan/IVDC-HBF13-21/2020 | Asia / China / Hubei / Wuhan | 2020-01-01 | Institute of Viral Disease Control and Prevention, China CDC | Institute of Viral Disease Control and Prevention, China CDC | William J. Liu, Peipei Liu, Xiang Zhao, Peihua Niu, Yingze Zhao, Wenwen Lei, Ziqian Xu, Shumei Zou, Wei Zhen, Beiwei Ye, Mengjie Yang, Weifeng Shi, Roujian Lu, Wenjie Tan, Zhixiao Chen, Yuchao Wu, Juan Song, Weimin Zhou, Dayan Wang, Jun Han, Wenbo Xu, George F. Gao, Guizhen Wu |
| **EPI_ISL_ 408665** | hCoV-19/Japan/TY-WK-012/2020 | Asia / Japan / Tokyo | 2020-01-29 | Dept. of Virology III, National Institute of Infectious Diseases | Pathogen Genomics Center, National Institute of Infectious Diseases | Tsuyoshi Sekizuka, Shutoku Matsuyama, Naganori Nao, Kazuya Shirato, Makoto Takeda, Makoto Kuroda |
| **EPI_ISL_ 408666** | hCoV-19/Japan/TY-WK-501/2020 | Asia / Japan / Tokyo | 2020-01-31 | Dept. of Virology III, National Institute of Infectious Diseases | Pathogen Genomics Center, National Institute of Infectious Diseases | Tsuyoshi Sekizuka, Shutoku Matsuyama, Naganori Nao, Kazuya Shirato, Makoto Takeda, Makoto Kuroda |
| **EPI_ISL_ 408667** | hCoV-19/Japan/TY-WK-521/2020 | Asia / Japan / Tokyo | 2020-01-31 | Dept. of Virology III, National Institute of Infectious Diseases | Pathogen Genomics Center, National Institute of Infectious Diseases | Tsuyoshi Sekizuka, Shutoku Matsuyama, Naganori Nao, Kazuya Shirato, Makoto Takeda, Makoto Kuroda |
| **EPI_ISL_ 408668** | hCoV-19/Vietnam/VR03-38142/2020 | Asia / Vietnam / Thanh Hoa | 2020-01-24 | National Influenza Center - National Institute of Hygiene and Epidemiology (NIHE) | National Influenza Center - National Institute of Hygiene and Epidemiology (NIHE) | Ung Thi Hong Trang, Hoang Vu Mai Phuong, Nguyen Le Khanh Hang, Nguyen Vu Son, Le Thi Thanh, Vuong Duc Cuong, Nguyen Phuong Anh, Pham Thi Hien, Tran Thu Huong, Le Thi Quynh Mai, |
| **EPI_ISL_ 408669** | hCoV-19/Japan/KY-V-029/2020 | Asia / Japan / Kyoto | 2020-01-29 | Dept. of Virology III, National Institute of Infectious Diseases | Pathogen Genomics Center, National Institute of Infectious Diseases | Tsuyoshi Sekizuka, Shutoku Matsuyama, Naganori Nao, Kazuya Shirato, Makoto Takeda, Makoto Kuroda |
| **EPI_ISL_ 408670** | hCoV-19/USA/ WI1/2020 | North America / USA / Wisconsin | 2020-01-31 | Wisconsin Department of Health Services | Pathogen Discovery, Respiratory Viruses Branch, Division of Viral Diseases, Centers for Dieases Control and Prevention | Jing Zhang, Anna Uehara, Krista Queen, Yan Li, Ying Tao, Clinton R. Paden, Xiaoyan Lu, Brian Lynch, Senthil Kumar K. Sakthivel, Brett L. Whitaker, Shifaq Kamili, Lijuan Wang, Janna' R. Murray, Susan I. Gerber, Stephen Lindstrom, Suxiang Tong |
| **EPI_ISL_ 408976** | hCoV-19/Australia/NSW02/2020 | Oceania / Australia / New South Wales / Sydney | 2020-01-22 | Centre for Infectious Diseases and Microbiology Laboratory Services | NSW Health Pathology - Institute of Clinical Pathology and Medical Research; Westmead Hospital; University of Sydney | Rockett R, Sadsad R, Eden J-S, Carter I, Rahman H, Holmes EC, O'Sullivan MV, Sintchenko V, Chen SC, Maddocks S, Kok J and Dwyer DE for the 2019-nCoV Study Group* |
| **EPI_ISL_ 408977** | hCoV-19/Australia/NSW03/2020 | Oceania / Australia / New South | 2020-01-25 | Serology, Virology and OTDS Laboratories (SAViD), NSW | NSW Health Pathology - Institute of Clinical Pathology and Medical Research; Centre for | Eden J-S, Carter I, Rahman H, Rawlinson W, Holmes EC, Rockett R, O'Sullivan MV, Sintchenko V, Chen SC, Maddocks S, Kok J and Dwyer DE for the 2019-nCoV Study Group* |

| | | | | | | |
|---|---|---|---|---|---|---|
| | | Wales / Sydney | | Health Pathology Randwick | Infectious Diseases and Microbiology Laboratory Services; Westmead Hospital; University of Sydney | |
| **EPI_ISL_ 408978** | hCoV-19/Wuhan/WH05/2020 | Asia / China / Hubei / Wuhan | 2020-02-07 | Wuhan Fourth Hospital | Beijing Genomics Institute (BGI) | Weijun Chen |
| **EPI_ISL_ 409067** | hCoV-19/USA/MA1/2020 | North America / USA / Massachusetts | 2020-01-29 | Massachusetts Department of Public Health | Pathogen Discovery, Respiratory Viruses Branch, Division of Viral Diseases, Centers for Dieases Control and Prevention | Clinton R. Paden, Jing Zhang, Krista Queen, Yan Li, Ying Tao, Anna Uehara, Xiaoyan Lu, Brian Lynch, Senthil Kumar K. Sakthivel, Brett L. Whitaker, Shifaq Kamili, Lijuan Wang, Janna' R. Murray, Susan I. Gerber, Stephen Lindstrom, Suxiang Tong |
| **EPI_ISL_ 410044** | hCoV-19/USA/CA6/2020 | North America / USA / California | 2020-01-27 | California Department of Public Health | Pathogen Discovery, Respiratory Viruses Branch, Division of Viral Diseases, Centers for Dieases Control and Prevention | Jing Zhang, Krista Queen, Yan Li, Ying Tao, Anna Uehara, Clinton R. Paden, Xiaoyan Lu, Brian Lynch, Senthil Kumar K. Sakthivel, Brett L. Whitaker, Shifaq Kamili, Lijuan Wang, Janna' R. Murray, Susan I. Gerber, Stephen Lindstrom, Suxiang Tong |
| **EPI_ISL_ 410045** | hCoV-19/USA/IL2/2020 | North America / USA / Illinois | 2020-01-28 | IL Department of Public Health Chicago Laboratory | Pathogen Discovery, Respiratory Viruses Branch, Division of Viral Diseases, Centers for Dieases Control and Prevention | Yan Li, Jing Zhang, Krista Queen, Ying Tao, Anna Uehara, Clinton R. Paden, Xiaoyan Lu, Brian Lynch, Senthil Kumar K. Sakthivel, Brett L. Whitaker, Shifaq Kamili, Lijuan Wang, Janna' R. Murray, Susan I. Gerber, Stephen Lindstrom, Suxiang Tong |
| **EPI_ISL_ 410218** | hCoV-19/Taiwan/NTU02/2020 | Asia / Taiwan / Taipei | 2020-02-05 | Department of Laboratory Medicine, National Taiwan University Hospital | Microbial Genomics Core Lab, National Taiwan University Centers of Genomic and Precision Medicine | Shiou-Hwei Yeh, You-Yu Lin, Ya-Yun Lai, Chiao-Ling Li, Shan-Chwen Chang, Pei-Jer Chen, Sui-Yuan Chang |
| **EPI_ISL_ 410301** | hCoV-19/Nepal/61/2020 | Asia / Nepal / Kathmandu | 2020-01-13 | National Influenza Centre, National Public Health Laboratory, Kathmandu, Nepal | The University of Hong Kong | Ranjit Sah, Runa Jha, Daniel Chu, Haogao Gu, Malik Peiris, Anup Bastola, Alfonso J. Rodriguez-Morales, Bibek Kumar Lal, Basu Dev Pandey, Leo Poon |
| **EPI_ISL_ 410486** | hCoV-19/France/RA739/2020 | Europe / France / Rhone-Alpes / Contamines | 2020-02-08 | CNR Virus des Infections Respiratoires - France SUD | CNR Virus des Infections Respiratoires - France SUD | Bal, Antonin; Destras, Gregory; Gaymard, Alexandre; Bouscambert-Duchamp, Maude; Cheynet, Valérie; Brengel-Pesce, Karen; Morfin-Sherpa, Florence; Valette, Martine; Josset, Laurence; Lina, Bruno. |
| **EPI_ISL_ 410531** | hCoV-19/Japan/NA-20-05-1/2020 | Asia / Japan / Nara | 2020-01-25 | Dept. of Pathology, National Institute of Infectious Diseases | Pathogen Genomics Center, National Institute of Infectious Diseases | Tsuyoshi Sekizuka, Harutaka Katano, Shutoku Matsuyama, Naganori Nao, Kazuya Shirato, Motoi Suzuki, Hideki Hasegawa, Takaji Wakita, Makoto Takeda, Tadaki Suzuki, Makoto Kuroda |
| **EPI_ISL_ 410532** | hCoV-19/Japan/OS-20-07-1/2020 | Asia / Japan / Osaka | 2020-01-23 | Dept. of Pathology, National Institute of Infectious Diseases | Pathogen Genomics Center, National Institute of Infectious Diseases | Tsuyoshi Sekizuka, Harutaka Katano, Shutoku Matsuyama, Naganori Nao, Kazuya Shirato, Motoi Suzuki, Hideki Hasegawa, Takaji Wakita, Makoto Takeda, Tadaki Suzuki, Makoto Kuroda |
| **EPI_ISL_ 410535** | hCoV-19/Singapore/4/2020 | Asia / Singapore | 2020-02-03 | National Centre for Infectious Diseases | Programme in Emerging Infectious Diseases, Duke-NUS Medical School | Danielle E Anderson, Martin Linster, Yan Zhuang, Jayanthi Jayakumar, David CB Lye, Yee Sin Leo, Barnaby E Young, Yvonne CF Su, Gavin JD Smith |
| **EPI_ISL_ 410536** | hCoV-19/Singapore/5/2020 | Asia / Singapore | 2020-02-06 | Singapore General Hospital, Molecular Laboratory, | Programme in Emerging Infectious Diseases, Duke-NUS Medical School | Danielle E Anderson, Martin Linster, Yan Zhuang, Jayanthi Jayakumar, Kian Sing Chan, Lynette LE Oon, Shirin Kalimuddin, Jenny GH Low, Yvonne CF Su, Gavin JD Smith |

| | | | | | | |
|---|---|---|---|---|---|---|
| | | | | Division of Pathology | | |
| **EPI_ISL_ 410537** | hCoV-19/Singapore/6/2020 | Asia / Singapore | 2020-02-09 | Singapore General Hospital, Molecular Laboratory, Division of Pathology | Programme in Emerging Infectious Diseases, Duke-NUS Medical School | Danielle E Anderson, Martin Linster, Yan Zhuang, Jayanthi Jayakumar, Kian Sing Chan, Lynette LE Oon, Shirin Kalimuddin, Jenny GH Low, Yvonne CF Su, Gavin JD Smith |
| **EPI_ISL_ 410545** | hCoV-19/Italy/INMI1-isl/2020 | Europe / Italy / Rome | 2020-01-29 | INMI Lazzaro Spallanzani IRCCS | Laboratory of Virology, INMI Lazzaro Spallanzani IRCCS | Maria R. Capobianchi, Cesare E. M. Gruber, Martina Rueca, Barbara Bartolini, Francesco Messina, Emanuela Giombini, Francesca Colavita, Concetta Castilletti, Eleonora Lalle, Fabrizio Carletti, Emanuele Nicastri, Giuseppe Ippolito. |
| **EPI_ISL_ 410713** | hCoV-19/Singapore/7/2020 | Asia / Singapore | 2020-01-27 | National Public Health Laboratory, National Centre for Infectious Diseases | National Public Health Laboratory, National Centre for Infectious Diseases | Octavia S, Mak TM, Cui L, Lin RTP |
| **EPI_ISL_ 410714** | hCoV-19/Singapore/8/2020 | Asia / Singapore | 2020-02-03 | National Public Health Laboratory, National Centre for Infectious Diseases | National Public Health Laboratory, National Centre for Infectious Diseases | Octavia S, Mak TM, Cui L, Lin RTP |
| **EPI_ISL_ 410715** | hCoV-19/Singapore/9/2020 | Asia / Singapore | 2020-02-04 | National Public Health Laboratory, National Centre for Infectious Diseases | National Public Health Laboratory, National Centre for Infectious Diseases | Octavia S, Mak TM, Cui L, Lin RTP |
| **EPI_ISL_ 410716** | hCoV-19/Singapore/10/2020 | Asia / Singapore | 2020-02-04 | National Public Health Laboratory, National Centre for Infectious Diseases | National Centre for Infectious Diseases, National Centre for Infectious Diseases | Octavia S, Mak TM, Cui L, Lin RTP |
| **EPI_ISL_ 410717** | hCoV-19/Australia/QLD03/2020 | Oceania / Australia / Queensland / Gold Coast | 2020-02-05 | Pathology Queensland | Public Health Virology Laboratory | Ben Huang, Alyssa Pyke, Amanda De Jong, Andrew Van Den Hurk, Carmel Taylor, David Warrilow, Doris Genge, Elisabeth Gamez, Glen Hewitson, Ian Maxwell Mackay, Inga Sultana, Jamie McMahon, Jean Barcelon, Judy Northill, Mitchell Finger, Natalie Simpson, Neelima Nair, Peter Burtonclay, Peter Moore, Sarah Wheatley, Sean Moody, Sonja Hall-Mendelin, Timothy Gardam, and Frederick Moore. |
| **EPI_ISL_ 410718** | hCoV-19/Australia/QLD04/2020 | Oceania / Australia / Queensland / Gold Coast | 2020-02-05 | Pathology Queensland | Public Health Virology Laboratory | Ben Huang, Alyssa Pyke, Amanda De Jong, Andrew Van Den Hurk, Carmel Taylor, David Warrilow, Doris Genge, Elisabeth Gamez, Glen Hewitson, Ian Maxwell Mackay, Inga Sultana, Jamie McMahon, Jean Barcelon, Judy Northill, Mitchell Finger, Natalie Simpson, Neelima Nair, Peter Burtonclay, Peter Moore, Sarah Wheatley, Sean Moody, Sonja Hall-Mendelin, Timothy Gardam, and Frederick Moore. |
| **EPI_ISL_ 410719** | hCoV-19/Singapore/11/2020 | Asia / Singapore | 2020-02-02 | National Public Health Laboratory | National Public Health Laboratory | Octavia S, Mak TM, Cui L, Lin RTP |
| **EPI_ISL_ 411060** | hCoV-19/Fujian/8/2020 | Asia / China / Fujian | 2020-01-21 | Fujian Center for Disease Control and Prevention | Fujian Center for Disease Control and Prevention | Chen Wei, Zhang Yanhua, He Wenxiang, Weng Yuwei |

| | | | | | | |
|---|---|---|---|---|---|---|
| **EPI_ISL_411066** | hCoV-19/Fujian/13/2020 | Asia / China / Fujian | 2020-01-22 | Fujian Center for Disease Control and Prevention | Fujian Center for Disease Control and Prevention | Chen Wei, Zhang Yanhua, He Wenxiang, Weng Yuwei |
| **EPI_ISL_411218** | hCoV-19/France/IDF0571/2020 | Europe / France / Ile-de-France / Paris | 2020-02-02 | Department of Infectious and Tropical Diseases, Bichat Claude Bernard Hospital, Paris | Laboratoire Virpath, CIRI U111, UCBL1, INSERM, CNRS, ENS Lyon | Olivier Terrier, Aurélien Traversier, Julien Fouret, Yazdan Yazdanpanah, Xavier Lescure, Catherine Legras-Lachuer, Alexandre Gaymard, Bruno Lina, Manuel Rosa-Calatrava |
| **EPI_ISL_411219** | hCoV-19/France/IDF0386-islP1/2020 | Europe / France / Ile-de-France / Paris | 2020-01-28 | Department of Infectious and Tropical Diseases, Bichat Claude Bernard Hospital, Paris | Laboratoire Virpath, CIRI U111, UCBL1, INSERM, CNRS, ENS Lyon | Olivier Terrier, Aurélien Traversier, Julien Fouret, Yazdan Yazdanpanah, Xavier Lescure, Alexandre Gaymard, Bruno Lina, Manuel Rosa-Calatrava |
| **EPI_ISL_411902** | hCoV-19/Cambodia/0012/2020 | Asia / Cambodia / Sihanoukville | 2020-01-27 | Virology Unit, Institut Pasteur du Cambodge. | Virology Unit, Institut Pasteur du Cambodge (Sequencing done by: Jessica E Manning/Jennifer A Bohl at Malaria and Vector Research Research Laboratory, National Institute of Allergy and Infectious Diseases and Vida Ahyong from Chan-Zuckerberg Biohub) | Erik A Karlsson, Jennifer A Bohl, Vida Ahyong, Veasna Duong, Philippe Dussart, Jessica E Manning. |
| **EPI_ISL_411950** | hCoV-19/Jiangsu/JS01/2020 | Asia / China / Jiangsu | 2020-01-23 | NHC Key laboratory of Enteric Pathogenic Microbiology, Institute of Pathogenic Microbiology | Jiangsu Provincial Center for Disease Control & Prevention | Lunbiao Cui,Kangchen Zhao,Xiaojuan Zhu,Yiyue Ge,Tao Wu,Bin Wu,Yin Chen,Fengcai Zhu,Baoli Zhu,Ming Wu |
| **EPI_ISL_411951** | hCoV-19/Sweden/01/2020 | Europe / Sweden | 2020-02-07 | unknown | Unit for Laboratory Development and Technology Transfer, Public Health Agency of Sweden | Bengner,M., Palmerus,M., Lindsjo,O., Lind Karlberg,M., Monteil,V., Appelberg,S., Brave,A., Muradrasoli,S. and Tegmark-Wisell,K. |
| **EPI_ISL_411952** | hCoV-19/Jiangsu/JS02/2020 | Asia / China / Jiangsu | 2020-01-24 | NHC Key laboratory of Enteric Pathogenic Microbiology, Institute of Pathogenic Microbiology | Jiangsu Provincial Center for Disease Control & Prevention | Kangchen Zhao, Xiaojuan Zhu, Lunbiao Cui, Tao Wu, Yiyue Ge, Bin Wu, Yin Chen, Fengcai Zhu, Baoli Zhu, Ming Wu |
| **EPI_ISL_411953** | hCoV-19/Jiangsu/JS03/2020 | Asia / China / Jiangsu | 2020-01-24 | NHC Key laboratory of Enteric Pathogenic Microbiology, Institute of Pathogenic Microbiology | Jiangsu Provincial Center for Disease Control & Prevention | Kangchen Zhao, Xiaojuan Zhu, Lunbiao Cui, Tao Wu, Yiyue Ge, Bin Wu, Yin Chen, Fengcai Zhu, Baoli Zhu, Ming Wu |
| **EPI_ISL_411954** | hCoV-19/USA/CA7/2020 | North America / USA / California | 2020-02-06 | California Department of Public Health | Pathogen Discovery, Respiratory Viruses Branch, Division of Viral Diseases, Centers for Dieases Control and Prevention | Krista Queen, Anna Uehara, Jing Zhang, Yan Li, Ying Tao, Clinton R. Paden, Haibin Wang, Shifaq Kamili, Xiaoyan Lu, Brian Lynch, Senthil Kumar K. Sakthivel, Brett L. Whitaker, Lijuan Wang, Janna' R. Murray, Susan I. Gerber, Stephen Lindstrom, Suxiang Tong |

| | | | | | | |
|---|---|---|---|---|---|---|
| **EPI_ISL_ 411955** | hCoV-19/USA/ CA8/2020 | North America / USA / California | 2020-02-10 | California Department of Public Health | Pathogen Discovery, Respiratory Viruses Branch, Division of Viral Diseases, Centers for Dieases Control and Prevention | Krista Queen, Anna Uehara, Jing Zhang, Yan Li, Ying Tao, Clinton R. Paden, Haibin Wang, Shifaq Kamili, Xiaoyan Lu, Brian Lynch, Senthil Kumar K. Sakthivel, Brett L. Whitaker, Lijuan Wang, Janna' R. Murray, Susan I. Gerber, Stephen Lindstrom, Suxiang Tong |
| **EPI_ISL_ 411956** | hCoV-19/USA/ TX1/2020 | North America / USA / Texas | 2020-02-11 | Texas Department of State Health Services | Pathogen Discovery, Respiratory Viruses Branch, Division of Viral Diseases, Centers for Dieases Control and Prevention | Krista Queen, Anna Uehara, Jing Zhang, Yan Li, Ying Tao, Clinton R. Paden, Haibin Wang, Shifaq Kamili, Xiaoyan Lu, Brian Lynch, Senthil Kumar K. Sakthivel, Brett L. Whitaker, Lijuan Wang, Janna' R. Murray, Susan I. Gerber, Stephen Lindstrom, Suxiang Tong |
| **EPI_ISL_ 412026** | hCoV-19/Hefei /2/2020 | Asia / China / Anhui / Hefei | 2020-02-23 | Second Hospital of Anhui Medical University | Second Hospital of Anhui Medical University | Changtai Wang, Zhongping Liua, Zixiang Chen, Xin Huang, Mengyuan Xua, Tengfei He, Mengji Lu, Zhenhua Zhang |
| **EPI_ISL_ 412028** | hCoV-19/Hong Kong/VM2000 1061/2020 | Asia / Hong Kong | 2020-01-22 | Hong Kong Department of Health | School of Public Health, The University of Hon g Kong | Dominic N.C. Tsang, Daniel K.W. Chu, Leo L.M. Poon, Malik Peiris |
| **EPI_ISL_ 412029** | hCoV-19/Hong Kong/VB20024 950/2020 | Asia / Hong Kong | 2020-01-30 | Hong Kong Department of Health | The University of Hong Kong | Dominic N.C. Tsang, Daniel K.W. Chu, Leo L.M. Poon, Malik Peiris |
| **EPI_ISL_ 412030** | hCoV-19/Hong Kong/VB20026 565/2020 | Asia / Hong Kong | 2020-02-01 | Hong Kong Department of Health | School of Public Health, The University of Hon g Kong | Dominic N.C. Tsang, Daniel K.W. Chu, Leo L.M. Poon, Malik Peiris |
| **EPI_ISL_ 412116** | hCoV-19/Engla nd/09c/2020 | Europe / United Kingdom / England | 2020-02-09 | Respiratory Virus Unit, Microbiology Services Colindale, Public Health England | Respiratory Virus Unit, Microbiology Services Colindale, Public Health England | Monica Galiano, Shahjahan Miah, Angie Lackenby, Omolola Akinbami, Tiina Talts, Leena Bhaw, Richard Myers, Steven Platt, Kirstin Edwards, Jonathan Hubb, Joanna Ellis, Maria Zambon |
| **EPI_ISL_ 412459** | hCoV-19/Jingz hou/HBCDC-H B-01/2020 | Asia / China / Hubei / Jingzhou | 2020-01-08 | Jingzhou Center for Disease Control and Prevention | Hubei Provincial Center for Disease Control and Prevention | Bin Fang, Xiang Li, Xiao Yu, Linlin Liu, Bo Yang, Faxian Zhan, Guojun Ye, Xixiang Huo, Junqiang Xu, Bo Yu, Kun Cai, Jing Li, Maoyi Chen,Jie Hu, Chunlin Mao, Yongzhong Jiang. |
| **EPI_ISL_ 412862** | hCoV-19/USA/ CA9/2020 | North America / USA / California / Solano | 2020-02-23 | California Department of Public Health | Pathogen Discovery, Respiratory Viruses Branch, Division of Viral Diseases, Centers for Disease Control and Prevention | Krista Queen, Anna Uehara, Jing Zhang, Yan Li, Ying Tao, Clinton R. Paden, Haibin Wang, Shifaq Kamili, Xiaoyan Lu, Brian Lynch, Senthil Kumar K. Sakthivel, Brett L. Whitaker, Lijuan Wang, Janna' R. Murray, Jasmine Padilla, Justin Lee, Susan I. Gerber, Stephen Lindstrom, Suxiang Tong |
| **EPI_ISL_ 412869** | hCoV-19/Sout h Korea/KCDC05 /2020 | Asia / South Korea /Seoul | 2020-01-30 | Division of Viral Diseases, Center for Laboratory Control of Infectious Diseases, Korea Centers for Diseases Control and Prevention | Division of Viral Diseases, Center for Laboratory Control of Infectious Diseases, Korea Centers for Diseases Control and Prevention | Jeong-Min Kim, Yoon-Seok Chung, Namjoo Lee, Mi-Seon Kim, Sang Hee Woo, Hye-Jun Jo, Sehee Park, Heui Man Kim, Myung Guk Han |
| **EPI_ISL_ 412870** | hCoV-19/Sout h Korea/KCDC06 /2020 | Asia / South Korea/ Seoul | 2020-01-30 | Division of Viral Diseases, Center for Laboratory Control of Infectious Diseases, Korea Centers for Diseases Control and Prevention | Division of Viral Diseases, Center for Laboratory Control of Infectious Diseases, Korea Centers for Diseases Control and Prevention | Jeong-Min Kim, Yoon-Seok Chung, Namjoo Lee, Mi-Seon Kim, Sang Hee Woo, Hye-Jun Jo, Sehee Park, Heui Man Kim, Myung Guk Han |
| **EPI_ISL_ 412871** | hCoV-19/Sout h Korea/KCDC07 /2020 | Asia / South Korea / Seoul | 2020-01-31 | Division of Viral Diseases, Center for Laboratory Control of | Division of Viral Diseases, Center for Laboratory Control of Infectious Diseases, | Jeong-Min Kim, Yoon-Seok Chung, Namjoo Lee, Mi-Seon Kim, Sang Hee Woo, Hye-Jun Jo, Sehee Park, Heui Man Kim, Myung Guk Han |

| | | | | | | |
|---|---|---|---|---|---|---|
| | | | | Infectious Diseases, Korea Centers for Diseases Control and Prevention | Korea Centers for Diseases Control and Prevention | |
| **EPI_ISL_ 412872** | hCoV-19/South Korea/KCDC12 /2020 | Asia / South Korea / Gyeonggi-do | 2020-02-01 | Division of Viral Diseases, Center for Laboratory Control of Infectious Diseases, Korea Centers for Diseases Control and Prevention | Division of Viral Diseases, Center for Laboratory Control of Infectious Diseases, Korea Centers for Diseases Control and Prevention | Jeong-Min Kim, Yoon-Seok Chung, Namjoo Lee, Mi-Seon Kim, Sang Hee Woo, Hye-Jun Jo, Sehee Park, Heui Man Kim, Myung Guk Han |
| **EPI_ISL_ 412873** | hCoV-19/South Korea/KCDC24 /2020 | Asia / South Korea / Chungcheongnam-do | 2020-02-06 | Division of Viral Diseases, Center for Laboratory Control of Infectious Diseases, Korea Centers for Diseases Control and Prevention | Division of Viral Diseases, Center for Laboratory Control of Infectious Diseases, Korea Centers for Diseases Control and Prevention | Jeong-Min Kim, Yoon-Seok Chung, Namjoo Lee, Mi-Seon Kim, Sang Hee Woo, Hye-Jun Jo, Sehee Park, Heui Man Kim, Myung Guk Han |
| **EPI_ISL_ 412898** | hCoV-19/Wuhan/HBCDC-HB-02/2019 | Asia / China / Hubei / Wuhan | 2019-12-30 | Wuhan Jinyintan Hospital | Hubei Provincial Center for Disease Control and Prevention | Bin Fang, Xiang Li, Xiao Yu, Linlin Liu, Bo Yang, Faxian Zhan, Guojun Ye, Xixiang Huo, Junqiang Xu, Bo Yu, Kun Cai, Jing Li, Yongzhong Jiang. |
| **EPI_ISL_ 412899** | hCoV-19/Wuhan/HBCDC-HB-03/2019 | Asia / China / Hubei / Wuhan | 2019-12-30 | Wuhan Jinyintan Hospital | Hubei Provincial Center for Disease Control and Prevention | Bin Fang, Xiang Li, Xiao Yu, Linlin Liu, Bo Yang, Faxian Zhan, Guojun Ye, Xixiang Huo, Junqiang Xu, Bo Yu, Kun Cai, Jing Li, Yongzhong Jiang. |
| **EPI_ISL_ 412912** | hCoV-19/Germany/Baden-Wuerttemberg-1/ 2020 | Europe / Germany / Baden-Wuerttemberg | 2020-02-25 | State Health Office Baden-Wuerttemberg | Charité Universitätsmedizin Berlin, Institute of Virology | Victor M Corman, Julia Schneider, Barbara Mühlemann, Talitha Veith, Jörn Beheim-Schwarzbach, Terry Jones, Rainer Oehme, Silke Fischer, Christian Drosten |
| **EPI_ISL_ 412964** | hCoV-19/Brazil /SPBR-01/2020 | South America / Brazil / Sao Paulo / Sao Paulo | 2020-02-25 | Hospital Israelita Albert Einstein | Instituto Adolfo Lutz Interdisciplinary Procedures Center Strategic Laboratory | Jaqueline Goes de Jesus, Claudio Tavares Sacchi, Daniela Bernardes Borges da Silva, Ingra Morales Claro, Flávia Cristina da Silva Sales, Claudia Regina Gonçalves, Joshua Quick, Maria do Carmo, Sampaio Tavares Timenetsky, Nicholas James Loman, Andrew Rambaut, Ester Cerdeira Sabino, Nuno Rodrigues Faria |
| **EPI_ISL_ 412965** | hCoV-19/Canada/BC_37_0-2/ 2020 | North America / Canada / British Columbia | 2020-02-16 | BCCDC Public Health Laboratory | BCCDC Public Health Laboratory | Harrigan, Prystajecky, Krajden, Lee, Kamelian, Lapointe, Choi, Hoang, Sekirov, Levett, Tyson, Loman, Quick, Li, Gilmour |
| **EPI_ISL_ 412966** | hCoV-19/China /IQTC01/2020 | Asia / China / Guangdong / Guangzhou | 2020-02-05 | unknown | Technology Centre, Guangzhou Customs | Shi,Y., Sun,J., Zheng,K., Huang,J. and Zhao,J. |
| **EPI_ISL_ 412967** | hCoV-19/China /IQTC02/2020 | Asia / China / Guangzhou | 2020-01-29 | unknown | Technology Centre, Guangzhou Customs | Shi,Y., Zheng,K., Sun,J., Huang,J., Zhu,A., Zhuang,Z., Dai,J., Chen,Z., Sun,F., Zhang,Z., Li,X. and Wang,Y. |
| **EPI_ISL_ 412968** | hCoV-19/Japan /Hu_DP_Kng_19-020/2020 | Asia / Japan | 2020-02-10 | unknown | Takayuki Hishiki Kanagawa Prefectural Institute of Public Health, Department of Microbiology | Hishiki,T., Suzuki,R., Sakuragi,J., Usui,K., Tanaka,Y., Kawai,J., Kogo,Y., Matsuki,Y., An,T., Hayashizaki,Y. and Takasaki,T. |

| | | | | | | |
|---|---|---|---|---|---|---|
| **EPI_ISL_ 412969** | hCoV-19/Japan/Hu_DP_Kng_19-027/2020 | Asia / Japan | 2020-02-10 | unknown | Takayuki Hishiki Kanagawa Prefectural Institute of Public Health, Department of Microbiology | Hishiki,T., Suzuki,R., Sakuragi,J., Usui,K., Tanaka,Y., Kawai,J., Kogo,Y., Matsuki,Y., An,T., Hayashizaki,Y. and Takasaki,T. |
| **EPI_ISL_ 412970** | hCoV-19/USA/WA2/2020 | North America / USA / Washington / Snohomish County | 2020-02-24 | Washington State Department of Health | Seattle Flu Study | Helen Chu, Michael Boeckh, Janet Englund, Michael Famulare, Barry Lutz, Deborah Nickerson, Mark Rieder, Lea Starita, Matthew Thompson, Jay Shendure, and Trevor Bedford |
| **EPI_ISL_ 412971** | hCoV-19/Finland/FIN-25/2020 | Europe / Finland / Helsinki | 2020-02-25 | HUS Diagnostiikkakeskus, Hallinto | Department of Virology Faculty of Medicine, Medicum University of Helsinki | Teemu Smura, Suvi Kuivanen, Hannimari Kallio-Kokko, Olli Vapalahti |
| **EPI_ISL_ 412972** | hCoV-19/Mexico/CDMX-InDRE_01/2020 | North America / Mexico / Mexico City | 2020-02-27 | Instituto Nacional de Enfermedades Respiratorias | Instituto de Diagnostico y Referencia Epidemiologicos (INDRE) | Ramirez-Gonzalez Ernesto, Garces-Ayala Fabiola, Araiza-Rodriguez Adnan, Mendieta-Condado Edgar, Rodriguez-Maldonado Abril, Wong-Arambula Claudia, Vazquez-Perez Joel, Martinez Arturo, Boukadida Celia, Munoz-Medina Esteban, Sanchez Alejandro, Isa Pavel, Taboada Blanca, Lopez Susana, Arias Carlos, Barrera-Badillo Gisela, Hernandez-Rivas Lucia, Lopez-Martinez Irma |
| **EPI_ISL_ 412973** | hCoV-19/Italy/CDG1/2020 | Europe / Italy / Lombardy | 2020-02-20 | Department of Infectious Diseases, Istituto Superiore di Sanità, Roma, Italy | Virology Laboratory, Scientific Department, Army Medical Center | Paola Stefanelli, Stefano Fiore, Antonella Marchi, Eleonora Benedetti, Concetta Fabiani, Giovanni Faggioni, Antonella Fortunato, Riccardo De Santis, Silvia Fillo, Anna Anselmo, Andrea Ciammaruconi, Stefano Palomba, Florigio Lista |
| **EPI_ISL_ 412974** | hCoV-19/Italy/SPL1/2020 | Europe / Italy / Rome | 2020-01-29 | Department of Infectious Diseases, Istituto Superiore di Sanità, Rome, Italy | Virology Laboratory, Scientific Department, Army Medical Center | Paola Stefanelli, Stefano Fiore, Antonella Marchi, Eleonora Benedetti, Concetta Fabiani, Giovanni Faggioni, Antonella Fortunato, Silvia Fillo, Riccardo De Santis, Andrea Ciammaruconi, Giancarlo Petralito, Filippo Molinari, Florigio Lista |
| **EPI_ISL_ 412975** | hCoV-19/Australia/NSW05/2020 | Oceania / Australia / New South Wales / Sydney | 2020-02-28 | Centre for Infectious Diseases and Microbiology Laboratory Services | NSW Health Pathology - Institute of Clinical Pathology and Medical Research; Westmead Hospital; University of Sydney | Eden J-S, Carter I, Rahman H, Holmes EC, Rockett R, O'Sullivan MV, Sintchenko V, Chen SC, Maddocks S, Kok J and Dwyer DE for the 2019-nCoV Study Group |
| **EPI_ISL_ 412978** | hCoV-19/Wuhan/HBCDC-HB-02/2020 | Asia / China / Hubei / Wuhan | 2020-01-17 | The Central Hospital Of Wuhan | Hubei Provincial Center for Disease Control and Prevention | Bin Fang, Xiang Li, Xiao Yu, Linlin Liu, Bo Yang, Faxian Zhan, Guojun Ye, Xixiang Huo, Junqiang Xu, Bo Yu, Kun Cai, Jing Li, Yongzhong Jiang. |
| **EPI_ISL_ 412979** | hCoV-19/Wuhan/HBCDC-HB-03/2020 | Asia / China / Hubei / Wuhan | 2020-01-18 | Union Hospital of Tongji Medical College, Huazhong University of Science and Technology | Hubei Provincial Center for Disease Control and Prevention | Bin Fang, Xiang Li, Xiao Yu, Linlin Liu, Bo Yang, Faxian Zhan, Guojun Ye, Xixiang Huo, Junqiang Xu, Bo Yu, Kun Cai, Jing Li, Yongzhong Jiang. |
| **EPI_ISL_ 412980** | hCoV-19/Wuhan/HBCDC-HB-04/2020 | Asia / China / Hubei / Wuhan | 2020-01-18 | Union Hospital of Tongji Medical College, Huazhong University of Science and Technology | Hubei Provincial Center for Disease Control and Prevention | Bin Fang, Xiang Li, Xiao Yu, Linlin Liu, Bo Yang, Faxian Zhan, Guojun Ye, Xixiang Huo, Junqiang Xu, Bo Yu, Kun Cai, Jing Li, Yongzhong Jiang. |
| **EPI_ISL_ 412981** | hCoV-19/Wuhan/HBCDC-HB-05/2020 | Asia / China / Hubei / Wuhan | 2020-01-18 | CR&WISCO GENERAL HOSPITAL | Hubei Provincial Center for Disease | Bin Fang, Xiang Li, Xiao Yu, Linlin Liu, Bo Yang, Faxian Zhan, Guojun Ye, Xixiang Huo, Junqiang Xu, Bo Yu, Kun Cai, Jing Li, Yongzhong Jiang. |

| | | | | | | |
|---|---|---|---|---|---|---|
| | | | | | Control and Prevention | |
| **EPI_ISL_ 412982** | hCoV-19/Wuhan/HBCDC-HB-06/2020 | Asia / China / Hubei / Wuhan | 2020-02-07 | Wuhan Lung Hospital | Hubei Provincial Center for Disease Control and Prevention | Bin Fang, Xiang Li, Xiao Yu, Linlin Liu, Bo Yang, Faxian Zhan, Guojun Ye, Xixiang Huo, Junqiang Xu, Bo Yu, Kun Cai, Jing Li, Yongzhong Jiang. |
| **EPI_ISL_ 412983** | hCoV-19/Tianmen/HBCDC-HB-07/2020 | Asia / China / Hubei / Tianmen | 2020-02-08 | Tianmen Center for Disease Control and Prevention | Hubei Provincial Center for Disease Control and Prevention | Bin Fang, Xiang Li, Xiao Yu, Linlin Liu, Bo Yang, Faxian Zhan, Guojun Ye, Xixiang Huo, Junqiang Xu, Bo Yu, Kun Cai, Jing Li, YiFa Zhu, Yangyang Tao,Xierong Li,Yongzhong Jiang. |
| **EPI_ISL_ 413014** | hCoV-19/Canada/ON-PHL2445/2020 | North America / Canada / Ontario | 2020-01-25 | Public Health Ontario Laboratory | Ontario Agency for Health Protection and Promotion (OAHPP) | Alireza Eshaghi, Samir N Patel, Jonathan B Gubbay, Vanessa G Allen, Christine Frantz, Aimin Li, Sandeep Nagra |
| **EPI_ISL_ 413015** | hCoV-19/Canada/ON-VIDO-01/2020 | North America / Canada / Ontario | 2020-01-23 | Public Health Ontario Laboratory | National Microbiology Laboratory | Shari Tyson, Anna Majer, Erika Landry, Morag Graham, Grace Seo, Philip Mabon, Natalie Knox, Adrian Zetner, Samira Mubareka, Rob Kozak, Jocelyne Lew, Darryl Falzarano, Gerdts Volker, Jonathan Gubbay, Stephanie Booth, Guillaume Poliquin, Tom Graefenhan, Matthew Gilmour, Nathalie Bastien, Yan Li, Timothy Booth |
| **EPI_ISL_ 413016** | hCoV-19/Brazil/SPBR-02/2020 | South America / Brazil / Sao Paulo / Sao Paulo | 2020-02-28 | Hospital Israelita Albert Einstein | Instituto Adolfo Lutz, Interdisciplinary Procedures Center, Strategic Laboratory | Jaqueline Goes de Jesus, Claudio Tavares Sacchi, Fabiana Cristina Pereira dos Santos, Ingra Morales Claro, Flávia Cristina da Silva Sales, Claudia Regina Gonçalves, Joshua Quick, Maria do Carmo Sampaio Tavares Timenetsky, Nicholas James Loman, Andrew Rambaut, Ester Cerdeira Sabino, Nuno Rodrigues Faria |
| **EPI_ISL_ 413017** | hCoV-19/South Korea/KUMC01/2020 | Asia / South Korea | 2020-02-06 | Department of Microbiology, Institute for Viral Diseases, College of Medicine, Korea University | Department of Microbiology, Institute for Viral Diseases, College of Medicine, Korea University | Changmin Kang, Joon-Yong Bae, Jungmin Lee, Heedo Park, Juyoung Cho, Jeonghun Kim, Gee eun Lee, Cui Chunguang, Kyeong-ryeol Shin, Dong Min Kim, Jin Il Kim, Man-Seong Park |
| **EPI_ISL_ 413018** | hCoV-19/South Korea/KUMC02/2020 | Asia / South Korea | 2020-02-06 | Department of Microbiology, Institute for Viral Diseases, College of Medicine, Korea University | Department of Microbiology, Institute for Viral Diseases, College of Medicine, Korea University | Changmin Kang, Joon-Yong Bae, Jungmin Lee, Heedo Park, Juyoung Cho, Jeonghun Kim, Gee eun Lee, Cui Chunguang, Kyeong-ryeol Shin, Dong Min Kim, Jin Il Kim, Man-Seong Park |
| **EPI_ISL_ 413019** | hCoV-19/Switzerland/1000477102/2020 | Europe / Switzerland / Zurich | 2020-02-26 | Department of Internal Medicine, Triemli Hospital | Institute of Medical Virology, University of Zurich | Stefan Schmutz, Maryam Zaheri, Verena Kufner, Patrick Redli, Fiona Steiner, Jon Huder, Riccarda Capaul, Andrea Zbinden, Jürg Böni, Michael Huber, Gerhard Eich, Alexandra Trkola |
| **EPI_ISL_ 413020** | hCoV-19/Switzerland/1000477377/2020 | Europe / Switzerland / Zurich | 2020-02-27 | Department of Internal Medicine, Triemli Hospital | Institute of Medical Virology, University of Zurich | Stefan Schmutz, Maryam Zaheri, Verena Kufner, Patrick Redli, Fiona Steiner, Jon Huder, Riccarda Capaul, Andrea Zbinden, Jürg Böni, Michael Huber, Gerhard Eich, Alexandra Trkola |
| **EPI_ISL_ 413021** | hCoV-19/Switzerland/1000477757/2020 | Europe / Switzerland / Zurich | 2020-02-29 | Klinik Hirslanden Zurich | Institute of Medical Virology, University of Zurich | Stefan Schmutz, Maryam Zaheri, Verena Kufner, Gabriela Ziltener, Patrick Redli, Fiona Steiner, Jon Huder, Riccarda Capaul, Andrea Zbinden, Jürg Böni, Michael Huber, Christian Ruef, Alexandra Trkola |
| **EPI_ISL_ 413022** | hCoV-19/Switzerland/1000477796/2020 | Europe / Switzerland / Zurich | 2020-02-29 | Division of Infectious Diseases, University Hospital Zurich | Institute of Medical Virology, University of Zurich | Stefan Schmutz, Maryam Zaheri, Verena Kufner, Gabriela Ziltener, Patrick Redli, Fiona Steiner, Jon Huder, Riccarda Capaul, Andrea Zbinden, Jürg Böni, Michael Huber, Roberto Speck, Alexandra Trkola |
| **EPI_ISL_ 413023** | hCoV-19/Switzerland/1000477797/2020 | Europe / Switzerland / Zurich | 2020-02-29 | Division of Infectious Diseases, | Institute of Medical Virology, University of Zurich | Stefan Schmutz, Maryam Zaheri, Verena Kufner, Gabriela Ziltener, Patrick Redli, Fiona Steiner, Jon Huder, Riccarda Capaul, Andrea Zbinden, Jürg |

| | | | | | | |
|---|---|---|---|---|---|---|
| | | | | University Hospital Zurich | | Böni, Michael Huber, Roberto Speck, Alexandra Trkola |
| **EPI_ISL_ 413024** | hCoV-19/Switzerland/1000477806/2020 | Europe / Switzerland / Zurich | 2020-02-29 | Division of Infectious Diseases, University Hospital Zurich | Institute of Medical Virology, University of Zurich | Stefan Schmutz, Maryam Zaheri, Verena Kufner, Gabriela Ziltener, Patrick Redli, Fiona Steiner, Jon Huder, Riccarda Capaul, Andrea Zbinden, Jürg Böni, Michael Huber, Roberto Speck, Alexandra Trkola |
| **EPI_ISL_ 413213** | hCoV-19/Australia/NSW06/2020 | Oceania / Australia / New South Wales / Sydney | 2020-02-29 | Centre for Infectious Diseases and Microbiology Laboratory Services | NSW Health Pathology - Institute of Clinical Pathology and Medical Research; Westmead Hospital; University of Sydney | Eden J-S, Carter I, Rahman H, Holmes EC, Rockett R, O'Sullivan MV, Sintchenko V, Chen SC, Maddocks S, Kok J and Dwyer DE for the 2019-nCoV Study Group* |
| **EPI_ISL_ 413214** | hCoV-19/Australia/NSW07/2020 | Oceania / Australia / New South Wales / Sydney | 2020-02-29 | Centre for Infectious Diseases and Microbiology Laboratory Services | NSW Health Pathology - Institute of Clinical Pathology and Medical Research; Westmead Hospital; University of Sydney | Eden J-S, Carter I, Rahman H, Holmes EC, Rockett R, O'Sullivan MV, Sintchenko V, Chen SC, Maddocks S, Kok J and Dwyer DE for the 2019-nCoV Study Group* |
| **EPI_ISL_ 413221** | hCoV-19/Scotland/CVR01/2020 | Europe / United Kingdom / Scotland | 2020-03-02 | West of Scotland Specialist Virology Centre, NHSGGC | MRC-University of Glasgow Centre for Virus Research | Emma Thomson, Antonia Ho; James Shephard, Shirin Ashraf; Kathy Smollett, Daniel Mair, Stephen Carmichael, Ana da Silva Filipe; Richard Orton, Josh Singer, David L Robertson; Andrew Rambaut; Alasdair MacLean, Rory Gunson. |
| **EPI_ISL_ 413455** | hCoV-19/USA/WA4-UW2/2020 | North America / USA / Washington | 2020-02-28 | Washington State Public Health Lab | University of Washington Virology Lab | Pavitra Roychoudhury, Arun Nalla, Hong Xie, Keith Jerome, Alexander Greninger |
| **EPI_ISL_ 413456** | hCoV-19/USA/WA-S2/2020 | North America / USA / Washington / King County | 2020-02-20 | Seattle Flu Study | Seattle Flu Study | Chu et al |
| **EPI_ISL_ 413457** | hCoV-19/USA/WA6-UW3/2020 | North America / USA / Washington | 2020-02-29 | Washington State Public Health Lab | UW Virology Lab | Pavitra Roychoudhury, Arun Nalla, Hong Xie, Keith Jerome, Alexander Greninger |
| **EPI_ISL_ 413458** | hCoV-19/USA/WA7-UW4/2020 | North America / USA / Washington | 2020-03-01 | Washington State Public Health Lab | UW Virology Lab | Pavitra Roychoudhury, Arun Nalla, Hong Xie, Keith Jerome, Alexander Greninger |
| **EPI_ISL_ 413485** | hCoV-19/Anhui/SZ005/2020 | Asia / China / Anhui / Suzhou | 2020-01-24 | Department of microbiology laboratory,Anhui Provincial Center for Disease Control and Prevention | Department of microbiology laboratory,Anhui Provincial Center for Disease Control and Prevention | Weiwei Li,Jun He,Yong Sun,Junling Yu,Qingqing Chen,Yuan Yuan,Yonglin Shi,Zhuhui Zhang,Yinglu Ge,Weidong Li,Bin Su,Zhirong Liu |
| **EPI_ISL_ 413486** | hCoV-19/USA/WA8-UW5/2020 | North America / USA / Washington | 2020-03-01 | Valley Medical Center | University of Washington Virology Lab | Pavitra Roychoudhury, Arun Nalla, Hong Xie, Keith Jerome, Alexander Greninger |
| **EPI_ISL_ 413487** | hCoV-19/USA/WA9-UW6/2020 | North America / USA / Washington | 2020-03-01 | Harborview Medical Center | University of Washington Virology Lab | Pavitra Roychoudhury, Arun Nalla, Hong Xie, Keith Jerome, Alexander Greninger |
| **EPI_ISL_ 413488** | hCoV-19/Germany/NRW-01/2020 | Europe / Germany / North Rhine Westphalia / | 2020-02-28 | Center of Medical Microbiology, Virology, and Hospital Hygiene, | Center of Medical Microbiology, Virology, and Hospital Hygiene, University of Duesseldorf | Ortwin Adams, Marcel Andree, Alexander Dilthey, Torsten Feldt, Sandra Hauka, Torsten Houwaart, Björn-Erik Jensen, Detlef Kindgen-Milles, Malte Kohns Vasconcelos, Klaus |

| | | | | | | |
|---|---|---|---|---|---|---|
| | | Heinsberg District | | University of Duesseldorf | | Pfeffer, Tina Senff, Daniel Strelow, Jörg Timm, Andreas Walker, Tobias Wienemann |
| **EPI_ISL_ 413489** | hCoV-19/Italy/ UniSR1/2020 | Europe / Italy / Lombardy / Milan | 2020-03-03 | Laboratorio di Microbiologia e Virologia, Università Vita-Salute San Raffaele, Milano | Laboratorio di Microbiologia e Virologia, Università Vita-Salute San Raffaele, Milano | R.A Diotti, E. Criscuolo, M. Castelli, V. Caputo, R. Ferrarese, M. Sampaolo, E. Boeri, I. Negri, V. Amato, G. Lo Raso, C. Di Resta, R. Burioni, M. Clementi, N. Mancini & N. Clementi |
| **EPI_ISL_ 413490** | hCoV-19/New Zealand/01/2020 | Oceania / New Zealand / Auckland | 2020-02-27 | Auckland Hospital | Institute of Environmental Science and Research (ESR) | Matt Storey, Xiaoyun Ren, Gary McAuliffe, Sally Roberts, Matthew Blakiston, Erasmus Smit, Lauren Jelly, Joep de Ligt |
| **EPI_ISL_ 413519** | hCoV-19/Beijing/231/2020 | Asia / China / Beijing | 2020-01-28 | unknown | Infectious Disease Control Center | Li,J., Li,L., Li,Z., Qiu,S., Song,H., Li,P. and Li,P. |
| **EPI_ISL_ 413520** | hCoV-19/Beijing/233/2020 | Asia / China / Beijing | 2020-01-28 | unknown | Infectious Disease Control Center | Li,J., Li,L., Li,Z., Qiu,S., Song,H., Li,P. and Li,P. |
| **EPI_ISL_ 413521** | hCoV-19/Beijing/235/2020 | Asia / China / Beijing | 2020-01-28 | unknown | Infectious Disease Control Center | Li,J., Li,L., Li,Z., Qiu,S., Song,H., Li,P. and Li,P. |
| **EPI_ISL_ 413522** | hCoV-19/India/1-27/2020 | Asia / India / Kerala | 2020-01-27 | Indian Council of Medical Research - National Institute of Virology | National Influenza Center, Indian Council of Medical Research - National Institute of Virology | Potdar V, Yadav PD, Choudhary ML, Shete-Aich A |
| **EPI_ISL_ 413523** | hCoV-19/India/1-31/2020 | Asia / India / Kerala | 2020-01-31 | Indian Council of Medical Research-National Institute of Virology | National Influenza Center, Indian Council of Medical Research-National Institute of Virology | Potdar V, Yadav PD, Choudhary ML, Shete-Aich A |
| **EPI_ISL_ 413550** | hCoV-19/Nigeria/Lagos01/2020 | Africa / Nigeria / Lagos | 2020-02-27 | Centre for Human and Zoonotic Virology (CHAZVY), College of Medicine University of Lagos/Lagos University Teaching Hospital (LUTH), part of the Laboratory Network of the Nigeria Centre for Disease Control (NCDC) | African Centre of Excellence for Genomics of Infectious Diseases (ACEGID), Redeemer's University, Ede, Osun State, Nigeria | Oluniyi P.E., Ajogbasile F.V., Kayode A., Oguzie J., Folarin O.A., Ihekweazu C. Happi C.T. |
| **EPI_ISL_ 413555** | hCoV-19/Wales/PHW1/2020 | Europe / United Kingdom / Wales | 2020-02-27 | Wales Specialist Virology Centre | Public Health Wales Microbiology Cardiff | Catherine Moore, Cen Sabu, Joanne Watkins, Sally Corden, Tom Connor |
| **EPI_ISL_ 413556** | hCoV-19/Wales/PHW2/2020 | Europe / United Kingdom / Wales | 2020-03-04 | Wales Specialist Virology Centre | Public Health Wales Microbiology Cardiff | Catherine Moore, Tim Jones, Joanne Watkins, Sally Corden, Tom Connor |
| **EPI_ISL_ 413557** | hCoV-19/USA/ CA-CDPH-UC1/ 2020 | North America / USA / California / Sonoma County | 2020-02-28 | California Department of Public Health | Chiu Laboratory, University of California, San Francisco | Xianding Deng, Scot Federman, Chao-Yang Pan, Hugo Guevara,Wei Gu, Debra A. Wadford, and Charles Y. Chiu |
| **EPI_ISL_ 413558** | hCoV-19/USA/ CA-CDPH-UC2/ 2020 | North America / USA / California / | 2020-02-27 | California Department of Public Health | Chiu Laboratory, University of California, San Francisco | Xianding Deng, Scot Federman, Chao-Yang Pan, Hugo Guevara,Wei Gu, Debra A. Wadford, and Charles Y. Chiu |

| | | | | | | |
|---|---|---|---|---|---|---|
| | | Solano County | | | | |
| **EPI_ISL_ 413559** | hCoV-19/USA/ CA-CDPH-UC3/ 2020 | North America / USA / California / Solano County | 2020-02-27 | California Department of Public Health | Chiu Laboratory, University of California, San Francisco | Xianding Deng, Scot Federman, Chao-Yang Pan, Hugo Guevara,Wei Gu, Debra A. Wadford, and Charles Y. Chiu |
| **EPI_ISL_ 413560** | hCoV-19/USA/ WA-S3/2020 | North America / USA / Washington | 2020-02-28 | Seattle Flu Study | Seattle Flu Study | Chu et al |
| **EPI_ISL_ 413561** | hCoV-19/USA/ CA-CDPH-UC4/ 2020 | North America / USA / California / Solano County | 2020-02-27 | California Department of Public Health | Chiu Laboratory, University of California, San Francisco | Xianding Deng, Scot Federman, Chao-Yang Pan, Hugo Guevara,Wei Gu, Debra A. Wadford, and Charles Y. Chiu |
| **EPI_ISL_ 413562** | hCoV-19/USA/ WA11-UW7/2 020 | North America / USA / Washington | 2020-03-02 | UW Virology Lab | UW Virology Lab | Pavitra Roychoudhury, Hong Xie, Keith Jerome, Alexander Greninger |
| **EPI_ISL_ 413563** | hCoV-19/USA/ WA12-UW8/2 020 | North America / USA / Washington | 2020-03-03 | UW Virology Lab | UW Virology Lab | Pavitra Roychoudhury, Hong Xie, Keith Jerome, Alexander Greninger |
| **EPI_ISL_ 413564** | hCoV-19/Neth erlands/Andel _1365066/202 0 | Europe / Netherlands / Andel | 2020-03-01 | MHC West-Brabant | Erasmus Medical Center | David Nieuwenhuijse, Bas Oude Munnink, Reina Sikkema, Claudia Schapendonk, Irina Chestakova, Anne van der Linden, Mark Pronk, Pascal Lexmond, Corien Swaan, Manon Haverkate, Madelief Mollers, Mart Stein, Sandra Kengne Kamga Mobou, Jeroen van Kampen, Jolanda Voermans, Aura Timen, Corine GeurtsvanKessel, Annemiek van der Eijk, Richard Molenkamp, Marion Koopmans, on behalf of the Dutch national COVID-19 response team. |
| **EPI_ISL_ 413565** | hCoV-19/Neth erlands/Berlicu m_1363564/2 020 | Europe / Netherlands / Berlicum | 2020-02-24 | Foundation Pamm | Erasmus Medical Center | David Nieuwenhuijse, Bas Oude Munnink, Reina Sikkema, Claudia Schapendonk, Irina Chestakova, Anne van der Linden, Mark Pronk, Pascal Lexmond, Corien Swaan, Manon Haverkate, Madelief Mollers, Mart Stein, Sandra Kengne Kamga Mobou, Jeroen van Kampen, Jolanda Voermans, Aura Timen, Corine GeurtsvanKessel, Annemiek van der Eijk, Richard Molenkamp, Marion Koopmans, on behalf of the Dutch national COVID-19 response team. |
| **EPI_ISL_ 413566** | hCoV-19/Neth erlands/Blaricu m_1364780/2 020 | Europe / Netherlands / Blaricum | 2020-03-02 | MHC Gooi & Vechtstreek | Erasmus Medical Center | David Nieuwenhuijse, Bas Oude Munnink, Reina Sikkema, Claudia Schapendonk, Irina Chestakova, Anne van der Linden, Mark Pronk, Pascal Lexmond, Corien Swaan, Manon Haverkate, Madelief Mollers, Mart Stein, Sandra Kengne Kamga Mobou, Jeroen van Kampen, Jolanda Voermans, Aura Timen, Corine GeurtsvanKessel, Annemiek van der Eijk, Richard Molenkamp, Marion Koopmans, on behalf of the Dutch national COVID-19 response team. |
| **EPI_ISL_ 413568** | hCoV-19/Neth erlands/Dalen _1363624/202 0 | Europe / Netherlands / Dalen | 2020-03-01 | MHC Drente | Erasmus Medical Center | David Nieuwenhuijse, Bas Oude Munnink, Reina Sikkema, Claudia Schapendonk, Irina Chestakova, Anne van der Linden, Mark Pronk, Pascal Lexmond, Corien Swaan, Manon Haverkate, Madelief Mollers, Mart Stein, Sandra Kengne Kamga Mobou, Jeroen van Kampen, Jolanda Voermans, Aura Timen, Corine GeurtsvanKessel, Annemiek van der Eijk, Richard Molenkamp, |

| | | | | | | |
|---|---|---|---|---|---|---|
| | | | | | | Marion Koopmans, on behalf of the Dutch national COVID-19 response team. |
| **EPI_ISL_ 413569** | hCoV-19/Netherlands/Delft_1363424/2020 | Europe / Netherlands / Delft | 2020-02-28 | RIVM | Erasmus Medical Center | David Nieuwenhuijse, Bas Oude Munnink, Reina Sikkema, Claudia Schapendonk, Irina Chestakova, Anne van der Linden, Mark Pronk, Pascal Lexmond, Corien Swaan, Manon Haverkate, Madelief Mollers, Mart Stein, Sandra Kengne Kamga Mobou, Jeroen van Kampen, Jolanda Voermans, Aura Timen, Corine GeurtsvanKessel, Annemiek van der Eijk, Richard Molenkamp, Marion Koopmans, on behalf of the Dutch national COVID-19 response team. |
| **EPI_ISL_ 413570** | hCoV-19/Netherlands/Diemen_1363454/2020 | Europe / Netherlands / Diemen | 2020-02-28 | RIVM | Erasmus Medical Center | David Nieuwenhuijse, Bas Oude Munnink, Reina Sikkema, Claudia Schapendonk, Irina Chestakova, Anne van der Linden, Mark Pronk, Pascal Lexmond, Corien Swaan, Manon Haverkate, Madelief Mollers, Mart Stein, Sandra Kengne Kamga Mobou, Jeroen van Kampen, Jolanda Voermans, Aura Timen, Corine GeurtsvanKessel, Annemiek van der Eijk, Richard Molenkamp, Marion Koopmans, on behalf of the Dutch national COVID-19 response team. |
| **EPI_ISL_ 413571** | hCoV-19/Netherlands/Eindhoven_1363782/2020 | Europe / Netherlands / Eindhoven | 2020-03-02 | MHC Brabant Zuidoost | Erasmus Medical Center | David Nieuwenhuijse, Bas Oude Munnink, Reina Sikkema, Claudia Schapendonk, Irina Chestakova, Anne van der Linden, Mark Pronk, Pascal Lexmond, Corien Swaan, Manon Haverkate, Madelief Mollers, Mart Stein, Sandra Kengne Kamga Mobou, Jeroen van Kampen, Jolanda Voermans, Aura Timen, Corine GeurtsvanKessel, Annemiek van der Eijk, Richard Molenkamp, Marion Koopmans, on behalf of the Dutch national COVID-19 response team. |
| **EPI_ISL_ 413572** | hCoV-19/Netherlands/Haarlem_1363688/2020 | Europe / Netherlands / Haarlem | 2020-03-01 | MHC Kennemerland | Erasmus Medical Center | David Nieuwenhuijse, Bas Oude Munnink, Reina Sikkema, Claudia Schapendonk, Irina Chestakova, Anne van der Linden, Mark Pronk, Pascal Lexmond, Corien Swaan, Manon Haverkate, Madelief Mollers, Mart Stein, Sandra Kengne Kamga Mobou, Jeroen van Kampen, Jolanda Voermans, Aura Timen, Corine GeurtsvanKessel, Annemiek van der Eijk, Richard Molenkamp, Marion Koopmans, on behalf of the Dutch national COVID-19 response team. |
| **EPI_ISL_ 413573** | hCoV-19/Netherlands/Hardinxveld_Giessendam_1364806/2020 | Europe / Netherlands / Hardinxveld Giessendam | 2020-03-02 | Dienst Gezondheid & Jeugd Zuid-Holland Zuid | Erasmus Medical Center | David Nieuwenhuijse, Bas Oude Munnink, Reina Sikkema, Claudia Schapendonk, Irina Chestakova, Anne van der Linden, Mark Pronk, Pascal Lexmond, Corien Swaan, Manon Haverkate, Madelief Mollers, Mart Stein, Sandra Kengne Kamga Mobou, Jeroen van Kampen, Jolanda Voermans, Aura Timen, Corine GeurtsvanKessel, Annemiek van der Eijk, Richard Molenkamp, Marion Koopmans, on behalf of the Dutch national COVID-19 response team. |
| **EPI_ISL_ 413574** | hCoV-19/Netherlands/Helmond_1363548/2020 | Europe / Netherlands / Helmond | 2020-02-29 | MHC West-Brabant | Erasmus Medical Center | David Nieuwenhuijse, Bas Oude Munnink, Reina Sikkema, Claudia Schapendonk, Irina Chestakova, Anne van der Linden, Mark Pronk, Pascal Lexmond, Corien Swaan, Manon Haverkate, Madelief Mollers, Mart Stein, Sandra Kengne Kamga Mobou, Jeroen van Kampen, Jolanda Voermans, Aura Timen, Corine GeurtsvanKessel, Annemiek van der Eijk, Richard Molenkamp, Marion Koopmans, on behalf of the Dutch national COVID-19 response team. |
| **EPI_ISL_ 413575** | hCoV-19/NetherlandsL/Hout | Europe / Netherlands / Houten | 2020-02-29 | RIVM | Erasmus Medical Center | David Nieuwenhuijse, Bas Oude Munnink, Reina Sikkema, Claudia Schapendonk, Irina Chestakova, Anne van der Linden, Mark Pronk, Pascal |

| | | | | | | |
|---|---|---|---|---|---|---|
| | en_1363498/2020 | | | | | Lexmond, Corien Swaan, Manon Haverkate, Madelief Mollers, Mart Stein, Sandra Kengne Kamga Mobou, Jeroen van Kampen, Jolanda Voermans, Aura Timen, Corine GeurtsvanKessel, Annemiek van der Eijk, Richard Molenkamp, Marion Koopmans, on behalf of the Dutch national COVID-19 response team. |
| **EPI_ISL_413576** | hCoV-19/Netherlands/Loon_op_zand_1363512/2020 | Europe / Netherlands / Loon op zand | 2020-02-29 | RIVM | Erasmus Medical Center | David Nieuwenhuijse, Bas Oude Munnink, Reina Sikkema, Claudia Schapendonk, Irina Chestakova, Anne van der Linden, Mark Pronk, Pascal Lexmond, Corien Swaan, Manon Haverkate, Madelief Mollers, Mart Stein, Sandra Kengne Kamga Mobou, Jeroen van Kampen, Jolanda Voermans, Aura Timen, Corine GeurtsvanKessel, Annemiek van der Eijk, Richard Molenkamp, Marion Koopmans, on behalf of the Dutch national COVID-19 response team. |
| **EPI_ISL_413577** | hCoV-19/Netherlands/Naarden_1364774/2020 | Europe / Netherlands / Naarden | 2020-03-02 | MHC Gooi & Vechtstreek | Erasmus Medical Center | David Nieuwenhuijse, Bas Oude Munnink, Reina Sikkema, Claudia Schapendonk, Irina Chestakova, Anne van der Linden, Mark Pronk, Pascal Lexmond, Corien Swaan, Manon Haverkate, Madelief Mollers, Mart Stein, Sandra Kengne Kamga Mobou, Jeroen van Kampen, Jolanda Voermans, Aura Timen, Corine GeurtsvanKessel, Annemiek van der Eijk, Richard Molenkamp, Marion Koopmans, on behalf of the Dutch national COVID-19 response team. |
| **EPI_ISL_413578** | hCoV-19/Netherlands/Nieuwendijk_1363582/2020 | Europe / Netherlands / Nieuwendijk | 2020-03-01 | ErasmusMC | Erasmus Medical Center | David Nieuwenhuijse, Bas Oude Munnink, Reina Sikkema, Claudia Schapendonk, Irina Chestakova, Anne van der Linden, Mark Pronk, Pascal Lexmond, Corien Swaan, Manon Haverkate, Madelief Mollers, Mart Stein, Sandra Kengne Kamga Mobou, Jeroen van Kampen, Jolanda Voermans, Aura Timen, Corine GeurtsvanKessel, Annemiek van der Eijk, Richard Molenkamp, Marion Koopmans, on behalf of the Dutch national COVID-19 response team. |
| **EPI_ISL_413579** | hCoV-19/Netherlands/Nootdorp_1364222/2020 | Europe / Netherlands / Nootdorp | 2020-03-03 | MHC Haaglanden | Erasmus Medical Center | David Nieuwenhuijse, Bas Oude Munnink, Reina Sikkema, Claudia Schapendonk, Irina Chestakova, Anne van der Linden, Mark Pronk, Pascal Lexmond, Corien Swaan, Manon Haverkate, Madelief Mollers, Mart Stein, Sandra Kengne Kamga Mobou, Jeroen van Kampen, Jolanda Voermans, Aura Timen, Corine GeurtsvanKessel, Annemiek van der Eijk, Richard Molenkamp, Marion Koopmans, on behalf of the Dutch national COVID-19 response team. |
| **EPI_ISL_413580** | hCoV-19/Netherlands/Oisterwijk_1364072/2020 | Europe / Netherlands / Oisterwijk | 2020-03-02 | MHC Hart voor Brabant | Erasmus Medical Center | David Nieuwenhuijse, Bas Oude Munnink, Reina Sikkema, Claudia Schapendonk, Irina Chestakova, Anne van der Linden, Mark Pronk, Pascal Lexmond, Corien Swaan, Manon Haverkate, Madelief Mollers, Mart Stein, Sandra Kengne Kamga Mobou, Jeroen van Kampen, Jolanda Voermans, Aura Timen, Corine GeurtsvanKessel, Annemiek van der Eijk, Richard Molenkamp, Marion Koopmans, on behalf of the Dutch national COVID-19 response team. |
| **EPI_ISL_413581** | hCoV-19/Netherlands/Oss_1363500/2020 | Europe / Netherlands / Oss | 2020-02-29 | RIVM | Erasmus Medical Center | David Nieuwenhuijse, Bas Oude Munnink, Reina Sikkema, Claudia Schapendonk, Irina Chestakova, Anne van der Linden, Mark Pronk, Pascal Lexmond, Corien Swaan, Manon Haverkate, Madelief Mollers, Mart Stein, Sandra Kengne Kamga Mobou, Jeroen van Kampen, Jolanda Voermans, Aura Timen, Corine GeurtsvanKessel, Annemiek van der Eijk, Richard Molenkamp, |

| | | | | | | |
|---|---|---|---|---|---|---|
| | | | | | | Marion Koopmans, on behalf of the Dutch national COVID-19 response team. |
| **EPI_ISL_413582** | hCoV-19/Netherlands/Rotterdam_1363790/2020 | Europe / Netherlands / Rotterdam | 2020-03-01 | ErasmusMC | Erasmus Medical Center | David Nieuwenhuijse, Bas Oude Munnink, Reina Sikkema, Claudia Schapendonk, Irina Chestakova, Anne van der Linden, Mark Pronk, Pascal Lexmond, Corien Swaan, Manon Haverkate, Madelief Mollers, Mart Stein, Sandra Kengne Kamga Mobou, Jeroen van Kampen, Jolanda Voermans, Aura Timen, Corine GeurtsvanKessel, Annemiek van der Eijk, Richard Molenkamp, Marion Koopmans, on behalf of the Dutch national COVID-19 response team. |
| **EPI_ISL_413583** | hCoV-19/Netherlands/Rotterdam_1364040/2020 | Europe / Netherlands / Rotterdam | 2020-03-02 | MHC Rotterdam-Rijnmond | Erasmus Medical Center | David Nieuwenhuijse, Bas Oude Munnink, Reina Sikkema, Claudia Schapendonk, Irina Chestakova, Anne van der Linden, Mark Pronk, Pascal Lexmond, Corien Swaan, Manon Haverkate, Madelief Mollers, Mart Stein, Sandra Kengne Kamga Mobou, Jeroen van Kampen, Jolanda Voermans, Aura Timen, Corine GeurtsvanKessel, Annemiek van der Eijk, Richard Molenkamp, Marion Koopmans, on behalf of the Dutch national COVID-19 response team. |
| **EPI_ISL_413584** | hCoV-19/Netherlands/Rotterdam_1364740/2020 | Europe / Netherlands / Rotterdam | 2020-03-03 | unknown | Erasmus Medical Center | David Nieuwenhuijse, Bas Oude Munnink, Reina Sikkema, Claudia Schapendonk, Irina Chestakova, Anne van der Linden, Mark Pronk, Pascal Lexmond, Corien Swaan, Manon Haverkate, Madelief Mollers, Mart Stein, Sandra Kengne Kamga Mobou, Jeroen van Kampen, Jolanda Voermans, Aura Timen, Corine GeurtsvanKessel, Annemiek van der Eijk, Richard Molenkamp, Marion Koopmans, on behalf of the Dutch national COVID-19 response team. |
| **EPI_ISL_413586** | hCoV-19/Netherlands/Tilburg_1363354/2020 | Europe / Netherlands / Tilburg | 2020-02-27 | Foundation Elisabeth-Tweesteden Ziekenhuis | Erasmus Medical Center | David Nieuwenhuijse, Bas Oude Munnink, Reina Sikkema, Claudia Schapendonk, Irina Chestakova, Anne van der Linden, Mark Pronk, Pascal Lexmond, Corien Swaan, Manon Haverkate, Madelief Mollers, Mart Stein, Sandra Kengne Kamga Mobou, Jeroen van Kampen, Jolanda Voermans, Aura Timen, Corine GeurtsvanKessel, Annemiek van der Eijk, Richard Molenkamp, Marion Koopmans, on behalf of the Dutch national COVID-19 response team. |
| **EPI_ISL_413587** | hCoV-19/Netherlands/Tilburg_1364286/2020 | Europe / Netherlands / Tilburg | 2020-03-03 | Foundation Elisabeth-Tweesteden Ziekenhuis | Erasmus Medical Center | David Nieuwenhuijse, Bas Oude Munnink, Reina Sikkema, Claudia Schapendonk, Irina Chestakova, Anne van der Linden, Mark Pronk, Pascal Lexmond, Corien Swaan, Manon Haverkate, Madelief Mollers, Mart Stein, Sandra Kengne Kamga Mobou, Jeroen van Kampen, Jolanda Voermans, Aura Timen, Corine GeurtsvanKessel, Annemiek van der Eijk, Richard Molenkamp, Marion Koopmans, on behalf of the Dutch national COVID-19 response team. |
| **EPI_ISL_413588** | hCoV-19/Netherlands/Utrecht_1363564/2020 | Europe / Netherlands / Utrecht | 2020-03-01 | MHC Utrecht | Erasmus Medical Center | David Nieuwenhuijse, Bas Oude Munnink, Reina Sikkema, Claudia Schapendonk, Irina Chestakova, Anne van der Linden, Mark Pronk, Pascal Lexmond, Corien Swaan, Manon Haverkate, Madelief Mollers, Mart Stein, Sandra Kengne Kamga Mobou, Jeroen van Kampen, Jolanda Voermans, Aura Timen, Corine GeurtsvanKessel, Annemiek van der Eijk, Richard Molenkamp, Marion Koopmans, on behalf of the Dutch national COVID-19 response team. |
| **EPI_ISL_413589** | hCoV-19/Netherlands/Utrech | Europe / Netherlands / Utrecht | 2020-03-01 | MHC Utrecht | Erasmus Medical Center | David Nieuwenhuijse, Bas Oude Munnink, Reina Sikkema, Claudia Schapendonk, Irina Chestakova, Anne van der Linden, Mark Pronk, Pascal |

| | | | | | | |
|---|---|---|---|---|---|---|
| | t_1363628/2020 | | | | | Lexmond, Corien Swaan, Manon Haverkate, Madelief Mollers, Mart Stein, Sandra Kengne Kamga Mobou, Jeroen van Kampen, Jolanda Voermans, Aura Timen, Corine GeurtsvanKessel, Annemiek van der Eijk, Richard Molenkamp, Marion Koopmans, on behalf of the Dutch national COVID-19 response team. |
| **EPI_ISL_413590** | hCoV-19/Netherlands/Utrecht_1364066/2020 | Europe / Netherlands / Utrecht | 2020-03-02 | MHC Utrecht | Erasmus Medical Center | David Nieuwenhuijse, Bas Oude Munnink, Reina Sikkema, Claudia Schapendonk, Irina Chestakova, Anne van der Linden, Mark Pronk, Pascal Lexmond, Corien Swaan, Manon Haverkate, Madelief Mollers, Mart Stein, Sandra Kengne Kamga Mobou, Jeroen van Kampen, Jolanda Voermans, Aura Timen, Corine GeurtsvanKessel, Annemiek van der Eijk, Richard Molenkamp, Marion Koopmans, on behalf of the Dutch national COVID-19 response team. |
| **EPI_ISL_413591** | hCoV-19/Netherlands/Zeewolde_1365080/2020 | Europe / Netherlands / Zeewolde | 2020-03-02 | MHC Flevoland | Erasmus Medical Center | David Nieuwenhuijse, Bas Oude Munnink, Reina Sikkema, Claudia Schapendonk, Irina Chestakova, Anne van der Linden, Mark Pronk, Pascal Lexmond, Corien Swaan, Manon Haverkate, Madelief Mollers, Mart Stein, Sandra Kengne Kamga Mobou, Jeroen van Kampen, Jolanda Voermans, Aura Timen, Corine GeurtsvanKessel, Annemiek van der Eijk, Richard Molenkamp, Marion Koopmans, on behalf of the Dutch national COVID-19 response team. |
| **EPI_ISL_413592** | hCoV-19/Taiwan/NTU03/2020 | Asia / Taiwan / Taipei | 2020-03-02 | Department of Laboratory Medicine, National Taiwan University Hospital | Microbial Genomics Core Lab, National Taiwan University Centers of Genomic and Precision Medicine | Shiou-Hwei Yeh, You-Yu Lin, Ya-Yun Lai, Chiao-Ling Li, Shan-Chwen Chang, Pei-Jer Chen, Sui-Yuan Chang |
| **EPI_ISL_413593** | hCoV-19/Luxembourg/Lux1/2020 | Europe / Luxembourg | 2020-02-29 | Laboratoire National de Santé | Erasmus Medical Center | David Nieuwenhuijse, Bas Oude Munnink, Reina Sikkema, Claudia Schapendonk, Irina Chestakova, Anne van der Linden, Mark Pronk, Pascal Lexmond, T. Abdelrahman, G. Fournier, J. Mossong, T. Nguyen, Jeroen van Kampen, Jolanda Voermans, Corine GeurtsvanKessel, Annemiek van der Eijk, Richard Molenkamp, Marion Koopmans, on behalf of the Dutch national COVID-19 response team. |
| **EPI_ISL_413594** | hCoV-19/Australia/NSW08/2020 | Oceania / Australia / NSW / Sydney | 2020-02-28 | Centre for Infectious Diseases and Microbiology Laboratory Services | NSW Health Pathology - Institute of Clinical Pathology and Medical Research; Westmead Hospital; University of Sydney | Rockett R, Eden J-S, Lam C, Gray K, Timms, V, Gall, M, Alicia, A, Carter I, Rahman H, Holmes EC,, O'Sullivan MV, Sintchenko V, Chen SC, Maddocks S, Kok J and Dwyer DE for the 2019-nCoV Study Group* |
| **EPI_ISL_413596** | hCoV-19/Australia/NSW10/2020 | Oceania / Australia / NSW / Sydney | 2020-02-28 | Centre for Infectious Diseases and Microbiology - Public Health | NSW Health Pathology - Institute of Clinical Pathology and Medical Research; Westmead Hospital; University of Sydney | Rockett R, Eden J-S, Lam C, Gray K, Timms, V, Gall, M, Carter I, Rahman H, Holmes EC, O'Sullivan MV, Sintchenko V, Chen SC, Maddocks S, Kok J and Dwyer DE for the 2019-nCoV Study Group* |
| **EPI_ISL_413597** | hCoV-19/Australia/NSW11/2020 | Oceania / Australia / NSW / Sydney | 2020-03-02 | Centre for Infectious Diseases and Microbiology-Public Health | NSW Health Pathology - Institute of Clinical Pathology and Medical Research; Westmead Hospital; University of Sydney | Lam C, Eden J-S, Rockett R, Gray K, Timms, V, Gall, M, Carter I, Rahman H, Holmes EC, O'Sullivan MV, Sintchenko V, Chen SC, Maddocks S, Kok J and Dwyer DE for the 2019-nCoV Study Group* |
| **EPI_ISL_413598** | hCoV-19/Australia/NSW12/2020 | Oceania / Australia / NSW / Sydney | 2020-03-04 | Centre for Infectious Diseases and Microbiology - Public Health | NSW Health Pathology - Institute of Clinical Pathology and Medical Research; Westmead | Gray K, Eden J-S, Lam C, Rockett R, Timms, V, Gall, M, Carter I, Rahman H, Holmes EC, O'Sullivan MV, Sintchenko V, Chen SC, Maddocks S, Kok J and Dwyer DE for the 2019-nCoV Study Group* |

| | | | | | | |
|---|---|---|---|---|---|---|
| | | | | | Hospital; University of Sydney | |
| **EPI_ISL_ 413599** | hCoV-19/Australia/NSW13/2020 | Oceania / Australia / NSW / Sydney | 2020-03-04 | Centre for Infectious Diseases and Microbiology - Public Health | NSW Health Pathology - Institute of Clinical Pathology and Medical Research; Westmead Hospital; University of Sydney | Timms, V, Eden J-S, Lam C, Gray K, Rockett R, Gall, M, Carter I, Rahman H, Holmes EC, O'Sullivan MV, Sintchenko V, Chen SC, Maddocks S, Kok J and Dwyer DE for the 2019-nCoV Study Group* |
| **EPI_ISL_ 413600** | hCoV-19/Australia/NSW14/2020 | Oceania / Australia / NSW / Sydney | 2020-03-03 | Centre for Infectious Diseases and Microbiology - Public Health | NSW Health Pathology - Institute of Clinical Pathology and Medical Research; Westmead Hospital; University of Sydney | Gall, M, Eden J-S, Lam C, Gray K, Timms, V, Rockett R, Carter I, Rahman H, Holmes EC, O'Sullivan MV, Sintchenko V, Chen SC, Maddocks S, Kok J and Dwyer DE for the 2019-nCoV Study Group* |
| **EPI_ISL_ 413601** | hCoV-19/USA/WA13-UW9/2020 | North America / USA / Washington | 2020-03-02 | UW Virology Lab | UW Virology Lab | Pavitra Roychoudhury, Hong Xie, Keith Jerome, Alexander Greninger |
| **EPI_ISL_ 413602** | hCoV-19/Finland/FIN03032020A/2020 | Europe / Finland / Helsinki | 2020-03-03 | Department of Virology and Immunology, University of Helsinki and Helsinki University Hospital, Huslab Finland | Department of Virology, Faculty of Medicine, University of Helsinki, Helsinki, Finland | Teemu Smura, Hannimari Kallio-Kokko, Olli Vapalahti |
| **EPI_ISL_ 413603** | hCoV-19/Finland/FIN03032020B/2020 | Europe / Finland / Helsinki | 2020-03-03 | Department of Virology and Immunology, University of Helsinki and Helsinki University Hospital, Huslab Finland | Department of Virology, Faculty of Medicine, University of Helsinki, Helsinki, Finland | Teemu Smura, Hannimari Kallio-Kokko, Olli Vapalahti |
| **EPI_ISL_ 413604** | hCoV-19/Finland/FIN03032020C/2020 | Europe / Finland / Helsinki | 2020-03-03 | Department of Virology and Immunology, University of Helsinki and Helsinki University Hospital, Huslab Finland | Department of Virology, Faculty of Medicine, University of Helsinki, Helsinki, Finland | Teemu Smura, Hannimari Kallio-Kokko, Olli Vapalahti |
| **EPI_ISL_ 413605** | hCoV-19/Finland/FIN01032020/2020 | Europe / Finland / Helsinki | 2020-03-01 | Department of Virology and Immunology, University of Helsinki and Helsinki University Hospital, Huslab Finland | Department of Virology, Faculty of Medicine, University of Helsinki, Helsinki, Finland | Teemu Smura, Hannimari Kallio-Kokko, Olli Vapalahti |
| **EPI_ISL_ 413647** | hCoV-19/Portugal/CV62/2020 | Europe / Portugal | 2020-03-01 | Centro Hospital do Porto, E.P.E. - H. Geral de Santo Antonio | Instituto Nacional de Saude (INSA) | Raquel Guiomar, Inês Costa, Pedro Pechirra, Joana Mendonça, Luís Vieira, Helena Ramos, Joana Isidro, Vítor Borges, João Paulo Gomes |
| **EPI_ISL_ 413648** | hCoV-19/Portugal/CV63/2020 | Europe / Portugal | 2020-03-01 | Centro Hospitalar e Universitário de Sao Joao, Porto | Instituto Nacional de Saude (INSA) | Raquel Guiomar, Inês Costa, Pedro Pechirra, Joana Mendonça, Luís Vieira, João Tiago Guimarães, Joana Isidro, Vítor Borges, João Paulo Gomes |

| | | | | | | |
|---|---|---|---|---|---|---|
| **EPI_ISL_413649** | hCoV-19/USA/WA14-UW10/2020 | North America / USA / Washington | 2020-03-05 | UW Virology Lab | UW Virology Lab | Pavitra Roychoudhury, Hong Xie, Keith Jerome, Alexander Greninger |
| **EPI_ISL_413650** | hCoV-19/USA/WA15-UW11/2020 | North America / USA / Washington | 2020-03-05 | UW Virology Lab | UW Virology Lab | Pavitra Roychoudhury, Hong Xie, Keith Jerome, Alexander Greninger |
| **EPI_ISL_413651** | hCoV-19/USA/WA16-UW12/2020 | North America / USA / Washington | 2020-03-05 | UW Virology Lab | UW Virology Lab | Pavitra Roychoudhury, Hong Xie, Keith Jerome, Alexander Greninger |
| **EPI_ISL_413652** | hCoV-19/USA/WA17-UW13/2020 | North America / USA / Washington | 2020-03-05 | UW Virology Lab | UW Virology Lab | Pavitra Roychoudhury, Hong Xie, Keith Jerome, Alexander Greninger |
| **EPI_ISL_413653** | hCoV-19/USA/WA18-UW14/2020 | North America / USA / Washington | 2020-03-05 | UW Virology Lab | UW Virology Lab | Pavitra Roychoudhury, Hong Xie, Keith Jerome, Alexander Greninger |
| **EPI_ISL_413691** | hCoV-19/China/WF0001/2020 | Asia / China | 2020-01 | Weifang Center for Disease Control and Prevention | Weifang Center for Disease Control and Prevention & BGI-Shenzhen | Qing Nie, Xingguang Li, Erik M Volz, Han Fu, Haowei Wang, Xiaoyue Xi, Wei Chen, Dehui Liu, Yingying Chen, Mengmeng Tian, Wei Tan, Junjie Zai, Wanying Sun, Jiandong Li, Junhua Li |
| **EPI_ISL_413692** | hCoV-19/China/WF0002/2020 | Asia / China | 2020-01 | Weifang Center for Disease Control and Prevention | Weifang Center for Disease Control and Prevention & BGI-Shenzhen | Qing Nie, Xingguang Li, Erik M Volz, Han Fu, Haowei Wang, Xiaoyue Xi, Wei Chen, Dehui Liu, Yingying Chen, Mengmeng Tian, Wei Tan, Junjie Zai, Wanying Sun, Jiandong Li, Junhua Li |
| **EPI_ISL_413693** | hCoV-19/China/WF0003/2020 | Asia / China | 2020-01 | Weifang Center for Disease Control and Prevention | Weifang Center for Disease Control and Prevention & BGI-Shenzhen | Qing Nie, Xingguang Li, Erik M Volz, Han Fu, Haowei Wang, Xiaoyue Xi, Wei Chen, Dehui Liu, Yingying Chen, Mengmeng Tian, Wei Tan, Junjie Zai, Wanying Sun, Jiandong Li, Junhua Li |
| **EPI_ISL_413694** | hCoV-19/China/WF0004/2020 | Asia / China | 2020-01 | Weifang Center for Disease Control and Prevention | Weifang Center for Disease Control and Prevention & BGI-Shenzhen | Qing Nie, Xingguang Li, Erik M Volz, Han Fu, Haowei Wang, Xiaoyue Xi, Wei Chen, Dehui Liu, Yingying Chen, Mengmeng Tian, Wei Tan, Junjie Zai, Wanying Sun, Jiandong Li, Junhua Li |
| **EPI_ISL_413697** | hCoV-19/China/WF0012/2020 | Asia / China | 2020-02 | Weifang Center for Disease Control and Prevention | Weifang Center for Disease Control and Prevention & BGI-Shenzhen | Qing Nie, Xingguang Li, Erik M Volz, Han Fu, Haowei Wang, Xiaoyue Xi, Wei Chen, Dehui Liu, Yingying Chen, Mengmeng Tian, Wei Tan, Junjie Zai, Wanying Sun, Jiandong Li, Junhua Li |
| **EPI_ISL_413711** | hCoV-19/China/WF0014/2020 | Asia / China | 2020-02 | Weifang Center for Disease Control and Prevention | Weifang Center for Disease Control and Prevention & BGI-Shenzhen | Qing Nie, Xingguang Li, Erik M Volz, Han Fu, Haowei Wang, Xiaoyue Xi, Wei Chen, Dehui Liu, Yingying Chen, Mengmeng Tian, Wei Tan, Junjie Zai, Wanying Sun, Jiandong Li, Junhua Li |
| **EPI_ISL_413729** | hCoV-19/China/WF0015/2020 | Asia / China | 2020-02 | Weifang Center for Disease Control and Prevention | Weifang Center for Disease Control and Prevention & BGI-Shenzhen | Qing Nie, Xingguang Li, Erik M Volz, Han Fu, Haowei Wang, Xiaoyue Xi, Wei Chen, Dehui Liu, Yingying Chen, Mengmeng Tian, Wei Tan, Junjie Zai, Wanying Sun, Jiandong Li, Junhua Li |
| **EPI_ISL_413746** | hCoV-19/China/WF0016/2020 | Asia / China | 2020-02 | Weifang Center for Disease Control and Prevention | Weifang Center for Disease Control and Prevention & BGI-Shenzhen | Qing Nie, Xingguang Li, Erik M Volz, Han Fu, Haowei Wang, Xiaoyue Xi, Wei Chen, Dehui Liu, Yingying Chen, Mengmeng Tian, Wei Tan, Junjie Zai, Wanying Sun, Jiandong Li, Junhua Li |
| **EPI_ISL_413748** | hCoV-19/China/WF0018/2020 | Asia / China | 2020-02 | Weifang Center for Disease Control and Prevention | Weifang Center for Disease Control and Prevention & BGI-Shenzhen | Qing Nie, Xingguang Li, Erik M Volz, Han Fu, Haowei Wang, Xiaoyue Xi, Wei Chen, Dehui Liu, Yingying Chen, Mengmeng Tian, Wei Tan, Junjie Zai, Wanying Sun, Jiandong Li, Junhua Li |
| **EPI_ISL_413749** | hCoV-19/China/WF0019/2020 | Asia / China | 2020-02 | Weifang Center for Disease Control and Prevention | Weifang Center for Disease Control and Prevention & BGI-Shenzhen | Qing Nie, Xingguang Li, Erik M Volz, Han Fu, Haowei Wang, Xiaoyue Xi, Wei Chen, Dehui Liu, Yingying Chen, Mengmeng Tian, Wei Tan, Junjie Zai, Wanying Sun, Jiandong Li, Junhua Li |

| | | | | | | |
|---|---|---|---|---|---|---|
| **EPI_ISL_413750** | hCoV-19/China/WF0020/2020 | Asia / China | 2020-02 | Weifang Center for Disease Control and Prevention | Weifang Center for Disease Control and Prevention & BGI-Shenzhen | Qing Nie, Xingguang Li, Erik M Volz, Han Fu, Haowei Wang, Xiaoyue Xi, Wei Chen, Dehui Liu, Yingying Chen, Mengmeng Tian, Wei Tan, Junjie Zai, Wanying Sun, Jiandong Li, Junhua Li |
| **EPI_ISL_413751** | hCoV-19/China/WF0021/2020 | Asia / China | 2020-02 | Weifang Center for Disease Control and Prevention | Weifang Center for Disease Control and Prevention & BGI-Shenzhen | Qing Nie, Xingguang Li, Erik M Volz, Han Fu, Haowei Wang, Xiaoyue Xi, Wei Chen, Dehui Liu, Yingying Chen, Mengmeng Tian, Wei Tan, Junjie Zai, Wanying Sun, Jiandong Li, Junhua Li |
| **EPI_ISL_413752** | hCoV-19/China/WF0023/2020 | Asia / China | 2020-02 | Weifang Center for Disease Control and Prevention | Weifang Center for Disease Control and Prevention & BGI-Shenzhen | Qing Nie, Xingguang Li, Erik M Volz, Han Fu, Haowei Wang, Xiaoyue Xi, Wei Chen, Dehui Liu, Yingying Chen, Mengmeng Tian, Wei Tan, Junjie Zai, Wanying Sun, Jiandong Li, Junhua Li |
| **EPI_ISL_413753** | hCoV-19/China/WF0024/2020 | Asia / China | 2020-02 | Weifang Center for Disease Control and Prevention | Weifang Center for Disease Control and Prevention & BGI-Shenzhen | Qing Nie, Xingguang Li, Erik M Volz, Han Fu, Haowei Wang, Xiaoyue Xi, Wei Chen, Dehui Liu, Yingying Chen, Mengmeng Tian, Wei Tan, Junjie Zai, Wanying Sun, Jiandong Li, Junhua Li |
| **EPI_ISL_413761** | hCoV-19/China/WF0026/2020 | Asia / China | 2020-02 | Weifang Center for Disease Control and Prevention | Weifang Center for Disease Control and Prevention & BGI-Shenzhen | Qing Nie, Xingguang Li, Erik M Volz, Han Fu, Haowei Wang, Xiaoyue Xi, Wei Chen, Dehui Liu, Yingying Chen, Mengmeng Tian, Wei Tan, Junjie Zai, Wanying Sun, Jiandong Li, Junhua Li |
| **EPI_ISL_413791** | hCoV-19/China/WF0028/2020 | Asia / China | 2020-02 | Weifang Center for Disease Control and Prevention | Weifang Center for Disease Control and Prevention & BGI-Shenzhen | Qing Nie, Xingguang Li, Erik M Volz, Han Fu, Haowei Wang, Xiaoyue Xi, Wei Chen, Dehui Liu, Yingying Chen, Mengmeng Tian, Wei Tan, Junjie Zai, Wanying Sun, Jiandong Li, Junhua Li |
| **EPI_ISL_413809** | hCoV-19/China/WF0029/2020 | Asia / China | 2020-02 | Weifang Center for Disease Control and Prevention | Weifang Center for Disease Control and Prevention & BGI-Shenzhen | Qing Nie, Xingguang Li, Erik M Volz, Han Fu, Haowei Wang, Xiaoyue Xi, Wei Chen, Dehui Liu, Yingying Chen, Mengmeng Tian, Wei Tan, Junjie Zai, Wanying Sun, Jiandong Li, Junhua Li |
| **EPI_ISL_413850** | hCoV-19/Guangdong/GDFS2020054-P0005/2020 | Asia / China / Guangdong | 2020-02-10 | Guangdong Provincial Institution of Public Health, Guangdong Provinical Center for Disease Control and Prevention | Guangdong Provincial Institution of Public Health | Jing Lu, Louis du Plessis, Liu Zhe, Jiufeng Sun, Sarah François, Huifang Lin, Moritz Kraemer, Jingju Peng, Qianlin Xiong, Runyu Yuan, Lilian Zeng, Pingping Zhou, Chuming Liang, Tao Liu, Wei Li, Juan Su, Huanying Zheng, Kang Min, Song Tie, Bo Peng, Shisong Fang, Wenzhe Su, Kuibiao Li, Ruilin Sun, Ru bai, Xi Tang, Minfeng Liang, Nuno Faria, Josh Quick, Andrew Rambaut, Verity Hill, Wenjun Ma, Nick Loman, Oliver Pybus, Changwen Ke |
| **EPI_ISL_413851** | hCoV-19/Guangdong/2020XN4373-P0039/2020 | Asia / China / Guangdong | 2020-01-30 | Guangdong Provincial Institution of Public Health, Guangdong Provinical Center for Disease Control and Prevention | Guangdong Provincial Institution of Public Health | Jing Lu, Louis du Plessis, Liu Zhe, Jiufeng Sun, Sarah François, Huifang Lin, Moritz Kraemer, Jingju Peng, Qianlin Xiong, Runyu Yuan, Lilian Zeng, Pingping Zhou, Chuming Liang, Tao Liu, Wei Li, Juan Su, Huanying Zheng, Kang Min, Song Tie, Bo Peng, Shisong Fang, Wenzhe Su, Kuibiao Li, Ruilin Sun, Ru bai, Xi Tang, Minfeng Liang, Nuno Faria, Josh Quick, Andrew Rambaut, Verity Hill, Wenjun Ma, Nick Loman, Oliver Pybus, Changwen Ke |
| **EPI_ISL_413852** | hCoV-19/Guangdong/2020XN4433-P0040/2020 | Asia / China / Guangdong | 2020-01-30 | Guangdong Provincial Institution of Public Health, Guangdong Provinical Center for Disease Control and Prevention | Guangdong Provincial Institution of Public Health | Jing Lu, Louis du Plessis, Liu Zhe, Jiufeng Sun, Sarah François, Huifang Lin, Moritz Kraemer, Jingju Peng, Qianlin Xiong, Runyu Yuan, Lilian Zeng, Pingping Zhou, Chuming Liang, Tao Liu, Wei Li, Juan Su, Huanying Zheng, Kang Min, Song Tie, Bo Peng, Shisong Fang, Wenzhe Su, Kuibiao Li, Ruilin Sun, Ru bai, Xi Tang, Minfeng Liang, Nuno Faria, Josh Quick, Andrew Rambaut, Verity Hill, Wenjun Ma, Nick Loman, Oliver Pybus, Changwen Ke |
| **EPI_ISL_413853** | hCoV-19/Guangdong/2020XN4243-P0035/2020 | Asia / China / Guangdong | 2020-01-30 | Guangdong Provincial Institution of Public Health, Guangdong | Guangdong Provincial Institution of Public Health | Jing Lu, Louis du Plessis, Liu Zhe, Jiufeng Sun, Sarah François, Huifang Lin, Moritz Kraemer, Jingju Peng, Qianlin Xiong, Runyu Yuan, Lilian Zeng, Pingping Zhou, Chuming Liang, Tao Liu, Wei Li, Juan Su, Huanying Zheng, Kang Min, Song |

| | | | | | | |
|---|---|---|---|---|---|---|
| | | | | Provincial Center for Disease Control and Prevention | | Tie, Bo Peng, Shisong Fang, Wenzhe Su, Kuibiao Li, Ruilin Sun, Ru bai, Xi Tang, Minfeng Liang, Nuno Faria, Josh Quick, Andrew Rambaut, Verity Hill, Wenjun Ma, Nick Loman, Oliver Pybus, Changwen Ke |
| **EPI_ISL_413854** | hCoV-19/Guangdong/2020XN4475-P0042/2020 | Asia / China / Guangdong | 2020-01-30 | Guangdong Provincial Institution of Public Health, Guangdong Provincial Center for Disease Control and Prevention | Guangdong Provincial Institution of Public Health | Jing Lu, Louis du Plessis, Liu Zhe, Jiufeng Sun, Sarah François, Huifang Lin, Moritz Kraemer, Jingju Peng, Qianlin Xiong, Runyu Yuan, Lilian Zeng, Pingping Zhou, Chuming Liang, Tao Liu, Wei Li, Juan Su, Huanying Zheng, Kang Min, Song Tie, Bo Peng, Shisong Fang, Wenzhe Su, Kuibiao Li, Ruilin Sun, Ru bai, Xi Tang, Minfeng Liang, Nuno Faria, Josh Quick, Andrew Rambaut, Verity Hill, Wenjun Ma, Nick Loman, Oliver Pybus, Changwen Ke |
| **EPI_ISL_413855** | hCoV-19/Guangdong/GD2020234-P0023/2020 | Asia / China / Guangdong | 2020-02-07 | Guangdong Provincial Institution of Public Health, Guangdong Provincial Center for Disease Control and Prevention | Guangdong Provincial Institution of Public Health | Jing Lu, Louis du Plessis, Liu Zhe, Jiufeng Sun, Sarah François, Huifang Lin, Moritz Kraemer, Jingju Peng, Qianlin Xiong, Runyu Yuan, Lilian Zeng, Pingping Zhou, Chuming Liang, Tao Liu, Wei Li, Juan Su, Huanying Zheng, Kang Min, Song Tie, Bo Peng, Shisong Fang, Wenzhe Su, Kuibiao Li, Ruilin Sun, Ru bai, Xi Tang, Minfeng Liang, Nuno Faria, Josh Quick, Andrew Rambaut, Verity Hill, Wenjun Ma, Nick Loman, Oliver Pybus, Changwen Ke |
| **EPI_ISL_413856** | hCoV-19/Guangdong/GD2020233-P0027/2020 | Asia / China / Guangdong | 2020-02-07 | Guangdong Provincial Institution of Public Health, Guangdong Provincial Center for Disease Control and Prevention | Guangdong Provincial Institution of Public Health | Jing Lu, Louis du Plessis, Liu Zhe, Jiufeng Sun, Sarah François, Huifang Lin, Moritz Kraemer, Jingju Peng, Qianlin Xiong, Runyu Yuan, Lilian Zeng, Pingping Zhou, Chuming Liang, Tao Liu, Wei Li, Juan Su, Huanying Zheng, Kang Min, Song Tie, Bo Peng, Shisong Fang, Wenzhe Su, Kuibiao Li, Ruilin Sun, Ru bai, Xi Tang, Minfeng Liang, Nuno Faria, Josh Quick, Andrew Rambaut, Verity Hill, Wenjun Ma, Nick Loman, Oliver Pybus, Changwen Ke |
| **EPI_ISL_413857** | hCoV-19/Guangdong/2020XN4448-P0002/2020 | Asia / China / Guangdong | 2020-01-31 | Guangdong Provincial Institution of Public Health, Guangdong Provincial Center for Disease Control and Prevention | Guangdong Provincial Institution of Public Health | Jing Lu, Louis du Plessis, Liu Zhe, Jiufeng Sun, Sarah François, Huifang Lin, Moritz Kraemer, Jingju Peng, Qianlin Xiong, Runyu Yuan, Lilian Zeng, Pingping Zhou, Chuming Liang, Tao Liu, Wei Li, Juan Su, Huanying Zheng, Kang Min, Song Tie, Bo Peng, Shisong Fang, Wenzhe Su, Kuibiao Li, Ruilin Sun, Ru bai, Xi Tang, Minfeng Liang, Nuno Faria, Josh Quick, Andrew Rambaut, Verity Hill, Wenjun Ma, Nick Loman, Oliver Pybus, Changwen Ke |
| **EPI_ISL_413858** | hCoV-19/Guangdong/2020XN4459-P0041/2020 | Asia / China / Guangdong | 2020-01-30 | Guangdong Provincial Institution of Public Health, Guangdong Provincial Center for Disease Control and Prevention | Guangdong Provincial Institution of Public Health | Jing Lu, Louis du Plessis, Liu Zhe, Jiufeng Sun, Sarah François, Huifang Lin, Moritz Kraemer, Jingju Peng, Qianlin Xiong, Runyu Yuan, Lilian Zeng, Pingping Zhou, Chuming Liang, Tao Liu, Wei Li, Juan Su, Huanying Zheng, Kang Min, Song Tie, Bo Peng, Shisong Fang, Wenzhe Su, Kuibiao Li, Ruilin Sun, Ru bai, Xi Tang, Minfeng Liang, Nuno Faria, Josh Quick, Andrew Rambaut, Verity Hill, Wenjun Ma, Nick Loman, Oliver Pybus, Changwen Ke |
| **EPI_ISL_413859** | hCoV-19/Guangdong/2020XN4276-P0037/2020 | Asia / China / Guangdong | 2020-01-31 | Guangdong Provincial Institution of Public Health, Guangdong Provincial Center for Disease Control and Prevention | Guangdong Provincial Institution of Public Health | Jing Lu, Louis du Plessis, Liu Zhe, Jiufeng Sun, Sarah François, Huifang Lin, Moritz Kraemer, Jingju Peng, Qianlin Xiong, Runyu Yuan, Lilian Zeng, Pingping Zhou, Chuming Liang, Tao Liu, Wei Li, Juan Su, Huanying Zheng, Kang Min, Song Tie, Bo Peng, Shisong Fang, Wenzhe Su, Kuibiao Li, Ruilin Sun, Ru bai, Xi Tang, Minfeng Liang, Nuno Faria, Josh Quick, Andrew Rambaut, Verity Hill, Wenjun Ma, Nick Loman, Oliver Pybus, Changwen Ke |

| | | | | | | |
|---|---|---|---|---|---|---|
| **EPI_ISL_413860** | hCoV-19/Guangdong/2020XN4273-P0036/2020 | Asia / China / Guangdong | 2020-01-30 | Guangdong Provincial Institution of Public Health, Guangdong Provinical Center for Disease Control and Prevention | Guangdong Provincial Institution of Public Health | Jing Lu, Louis du Plessis, Liu Zhe, Jiufeng Sun, Sarah François, Huifang Lin, Moritz Kraemer, Jingju Peng, Qianlin Xiong, Runyu Yuan, Lilian Zeng, Pingping Zhou, Chuming Liang, Tao Liu, Wei Li, Juan Su, Huanying Zheng, Kang Min, Song Tie, Bo Peng, Shisong Fang, Wenzhe Su, Kuibiao Li, Ruilin Sun, Ru bai, Xi Tang, Minfeng Liang, Nuno Faria, Josh Quick, Andrew Rambaut, Verity Hill, Wenjun Ma, Nick Loman, Oliver Pybus, Changwen Ke |
| **EPI_ISL_413861** | hCoV-19/Guangdong/GD2020080-P0010/2020 | Asia / China / Guangdong | 2020-02-01 | Guangdong Provincial Institution of Public Health, Guangdong Provinical Center for Disease Control and Prevention | Guangdong Provincial Institution of Public Health | Jing Lu, Louis du Plessis, Liu Zhe, Jiufeng Sun, Sarah François, Huifang Lin, Moritz Kraemer, Jingju Peng, Qianlin Xiong, Runyu Yuan, Lilian Zeng, Pingping Zhou, Chuming Liang, Tao Liu, Wei Li, Juan Su, Huanying Zheng, Kang Min, Song Tie, Bo Peng, Shisong Fang, Wenzhe Su, Kuibiao Li, Ruilin Sun, Ru bai, Xi Tang, Minfeng Liang, Nuno Faria, Josh Quick, Andrew Rambaut, Verity Hill, Wenjun Ma, Nick Loman, Oliver Pybus, Changwen Ke |
| **EPI_ISL_413862** | hCoV-19/Guangdong/GD2020086-P0021/2020 | Asia / China / Guangdong | 2020-02-01 | Guangdong Provincial Institution of Public Health, Guangdong Provinical Center for Disease Control and Prevention | Guangdong Provincial Institution of Public Health | Jing Lu, Louis du Plessis, Liu Zhe, Jiufeng Sun, Sarah François, Huifang Lin, Moritz Kraemer, Jingju Peng, Qianlin Xiong, Runyu Yuan, Lilian Zeng, Pingping Zhou, Chuming Liang, Tao Liu, Wei Li, Juan Su, Huanying Zheng, Kang Min, Song Tie, Bo Peng, Shisong Fang, Wenzhe Su, Kuibiao Li, Ruilin Sun, Ru bai, Xi Tang, Minfeng Liang, Nuno Faria, Josh Quick, Andrew Rambaut, Verity Hill, Wenjun Ma, Nick Loman, Oliver Pybus, Changwen Ke |
| **EPI_ISL_413863** | hCoV-19/Guangdong/GD2020087-P0008/2020 | Asia / China / Guangdong | 2020-02-01 | Guangdong Provincial Institution of Public Health, Guangdong Provinical Center for Disease Control and Prevention | Guangdong Provincial Institution of Public Health | Jing Lu, Louis du Plessis, Liu Zhe, Jiufeng Sun, Sarah François, Huifang Lin, Moritz Kraemer, Jingju Peng, Qianlin Xiong, Runyu Yuan, Lilian Zeng, Pingping Zhou, Chuming Liang, Tao Liu, Wei Li, Juan Su, Huanying Zheng, Kang Min, Song Tie, Bo Peng, Shisong Fang, Wenzhe Su, Kuibiao Li, Ruilin Sun, Ru bai, Xi Tang, Minfeng Liang, Nuno Faria, Josh Quick, Andrew Rambaut, Verity Hill, Wenjun Ma, Nick Loman, Oliver Pybus, Changwen Ke |
| **EPI_ISL_413864** | hCoV-19/Guangdong/GD2020246-P0028/2020 | Asia / China / Guangdong | 2020-02-09 | Guangdong Provincial Institution of Public Health, Guangdong Provinical Center for Disease Control and Prevention | Guangdong Provincial Institution of Public Health | Jing Lu, Louis du Plessis, Liu Zhe, Jiufeng Sun, Sarah François, Huifang Lin, Moritz Kraemer, Jingju Peng, Qianlin Xiong, Runyu Yuan, Lilian Zeng, Pingping Zhou, Chuming Liang, Tao Liu, Wei Li, Juan Su, Huanying Zheng, Kang Min, Song Tie, Bo Peng, Shisong Fang, Wenzhe Su, Kuibiao Li, Ruilin Sun, Ru bai, Xi Tang, Minfeng Liang, Nuno Faria, Josh Quick, Andrew Rambaut, Verity Hill, Wenjun Ma, Nick Loman, Oliver Pybus, Changwen Ke |
| **EPI_ISL_413866** | hCoV-19/Guangdong/GDSZ202008-P0020/2020 | Asia / China / Guangdong | 2020-02-05 | Guangdong Provincial Institution of Public Health, Guangdong Provinical Center for Disease Control and Prevention | Guangdong Provincial Institution of Public Health | Jing Lu, Louis du Plessis, Liu Zhe, Jiufeng Sun, Sarah François, Huifang Lin, Moritz Kraemer, Jingju Peng, Qianlin Xiong, Runyu Yuan, Lilian Zeng, Pingping Zhou, Chuming Liang, Tao Liu, Wei Li, Juan Su, Huanying Zheng, Kang Min, Song Tie, Bo Peng, Shisong Fang, Wenzhe Su, Kuibiao Li, Ruilin Sun, Ru bai, Xi Tang, Minfeng Liang, Nuno Faria, Josh Quick, Andrew Rambaut, Verity Hill, Wenjun Ma, Nick Loman, Oliver Pybus, Changwen Ke |
| **EPI_ISL_413867** | hCoV-19/Guangdong/GDSZ202015-P0019/2020 | Asia / China / Guangdong | 2020-02-05 | Guangdong Provincial Institution of Public Health, Guangdong Provinical Center | Guangdong Provincial Institution of Public Health | Jing Lu, Louis du Plessis, Liu Zhe, Jiufeng Sun, Sarah François, Huifang Lin, Moritz Kraemer, Jingju Peng, Qianlin Xiong, Runyu Yuan, Lilian Zeng, Pingping Zhou, Chuming Liang, Tao Liu, Wei Li, Juan Su, Huanying Zheng, Kang Min, Song Tie, Bo Peng, Shisong Fang, Wenzhe Su, Kuibiao |

| | | | | | | |
|---|---|---|---|---|---|---|
| | | | | for Disease Control and Prevention | | Li, Ruilin Sun, Ru bai, Xi Tang, Minfeng Liang, Nuno Faria, Josh Quick, Andrew Rambaut, Verity Hill, Wenjun Ma, Nick Loman, Oliver Pybus, Changwen Ke |
| **EPI_ISL_413868** | hCoV-19/Guangdong/DG-S9-P0045/2020 | Asia / China / Guangdong | 2020-02-28 | Guangdong Provincial Institution of Public Health, Guangdong Provinical Center for Disease Control and Prevention | Guangdong Provincial Institution of Public Health | Jing Lu, Louis du Plessis, Liu Zhe, Jiufeng Sun, Sarah François, Huifang Lin, Moritz Kraemer, Jingju Peng, Qianlin Xiong, Runyu Yuan, Lilian Zeng, Pingping Zhou, Chuming Liang, Tao Liu, Wei Li, Juan Su, Huanying Zheng, Kang Min, Song Tie, Bo Peng, Shisong Fang, Wenzhe Su, Kuibiao Li, Ruilin Sun, Ru bai, Xi Tang, Minfeng Liang, Nuno Faria, Josh Quick, Andrew Rambaut, Verity Hill, Wenjun Ma, Nick Loman, Oliver Pybus, Changwen Ke |
| **EPI_ISL_413869** | hCoV-19/Guangdong/FS-S42-P0046/2020 | Asia / China / Guangdong | 2020-02-28 | Guangdong Provincial Institution of Public Health, Guangdong Provinical Center for Disease Control and Prevention | Guangdong Provincial Institution of Public Health | Jing Lu, Louis du Plessis, Liu Zhe, Jiufeng Sun, Sarah François, Huifang Lin, Moritz Kraemer, Jingju Peng, Qianlin Xiong, Runyu Yuan, Lilian Zeng, Pingping Zhou, Chuming Liang, Tao Liu, Wei Li, Juan Su, Huanying Zheng, Kang Min, Song Tie, Bo Peng, Shisong Fang, Wenzhe Su, Kuibiao Li, Ruilin Sun, Ru bai, Xi Tang, Minfeng Liang, Nuno Faria, Josh Quick, Andrew Rambaut, Verity Hill, Wenjun Ma, Nick Loman, Oliver Pybus, Changwen Ke |
| **EPI_ISL_413870** | hCoV-19/Guangdong/FS-S48-P0047/2020 | Asia / China / Guangdong | 2020-02-28 | Guangdong Provincial Institution of Public Health, Guangdong Provinical Center for Disease Control and Prevention | Guangdong Provincial Institution of Public Health | Jing Lu, Louis du Plessis, Liu Zhe, Jiufeng Sun, Sarah François, Huifang Lin, Moritz Kraemer, Jingju Peng, Qianlin Xiong, Runyu Yuan, Lilian Zeng, Pingping Zhou, Chuming Liang, Tao Liu, Wei Li, Juan Su, Huanying Zheng, Kang Min, Song Tie, Bo Peng, Shisong Fang, Wenzhe Su, Kuibiao Li, Ruilin Sun, Ru bai, Xi Tang, Minfeng Liang, Nuno Faria, Josh Quick, Andrew Rambaut, Verity Hill, Wenjun Ma, Nick Loman, Oliver Pybus, Changwen Ke |
| **EPI_ISL_413871** | hCoV-19/Guangdong/MM-S1-P0048/2020 | Asia / China / Guangdong | 2020-02-28 | Guangdong Provincial Institution of Public Health, Guangdong Provinical Center for Disease Control and Prevention | Guangdong Provincial Institution of Public Health | Jing Lu, Louis du Plessis, Liu Zhe, Jiufeng Sun, Sarah François, Huifang Lin, Moritz Kraemer, Jingju Peng, Qianlin Xiong, Runyu Yuan, Lilian Zeng, Pingping Zhou, Chuming Liang, Tao Liu, Wei Li, Juan Su, Huanying Zheng, Kang Min, Song Tie, Bo Peng, Shisong Fang, Wenzhe Su, Kuibiao Li, Ruilin Sun, Ru bai, Xi Tang, Minfeng Liang, Nuno Faria, Josh Quick, Andrew Rambaut, Verity Hill, Wenjun Ma, Nick Loman, Oliver Pybus, Changwen Ke |
| **EPI_ISL_413872** | hCoV-19/Guangdong/SZ-N59-P0049/2020 | Asia / China / Guangdong | 2020-02-28 | Guangdong Provincial Institution of Public Health, Guangdong Provinical Center for Disease Control and Prevention | Guangdong Provincial Institution of Public Health | Jing Lu, Louis du Plessis, Liu Zhe, Jiufeng Sun, Sarah François, Huifang Lin, Moritz Kraemer, Jingju Peng, Qianlin Xiong, Runyu Yuan, Lilian Zeng, Pingping Zhou, Chuming Liang, Tao Liu, Wei Li, Juan Su, Huanying Zheng, Kang Min, Song Tie, Bo Peng, Shisong Fang, Wenzhe Su, Kuibiao Li, Ruilin Sun, Ru bai, Xi Tang, Minfeng Liang, Nuno Faria, Josh Quick, Andrew Rambaut, Verity Hill, Wenjun Ma, Nick Loman, Oliver Pybus, Changwen Ke |
| **EPI_ISL_413873** | hCoV-19/Guangdong/GZ-S6-P0050/2020 | Asia / China / Guangdong | 2020-02-28 | Guangdong Provincial Institution of Public Health, Guangdong Provinical Center for Disease Control and Prevention | Guangdong Provincial Institution of Public Health | Jing Lu, Louis du Plessis, Liu Zhe, Jiufeng Sun, Sarah François, Huifang Lin, Moritz Kraemer, Jingju Peng, Qianlin Xiong, Runyu Yuan, Lilian Zeng, Pingping Zhou, Chuming Liang, Tao Liu, Wei Li, Juan Su, Huanying Zheng, Kang Min, Song Tie, Bo Peng, Shisong Fang, Wenzhe Su, Kuibiao Li, Ruilin Sun, Ru bai, Xi Tang, Minfeng Liang, Nuno Faria, Josh Quick, Andrew Rambaut, Verity Hill, Wenjun Ma, Nick Loman, Oliver Pybus, Changwen Ke |

| | | | | | | |
|---|---|---|---|---|---|---|
| **EPI_ISL_413874** | hCoV-19/Guangdong/GDFS2020127-P0026/2020 | Asia / China / Guangdong | 2020-02-12 | Guangdong Provincial Institution of Public Health, Guangdong Provinical Center for Disease Control and Prevention | Guangdong Provincial Institution of Public Health | Jing Lu, Louis du Plessis, Liu Zhe, Jiufeng Sun, Sarah François, Huifang Lin, Moritz Kraemer, Jingju Peng, Qianlin Xiong, Runyu Yuan, Lilian Zeng, Pingping Zhou, Chuming Liang, Tao Liu, Wei Li, Juan Su, Huanying Zheng, Kang Min, Song Tie, Bo Peng, Shisong Fang, Wenzhe Su, Kuibiao Li, Ruilin Sun, Ru bai, Xi Tang, Minfeng Liang, Nuno Faria, Josh Quick, Andrew Rambaut, Verity Hill, Wenjun Ma, Nick Loman, Oliver Pybus, Changwen Ke |
| **EPI_ISL_413875** | hCoV-19/Guangdong/GDFS2020052-P0025/2020 | Asia / China / Guangdong | 2020-02-10 | Guangdong Provincial Institution of Public Health, Guangdong Provinical Center for Disease Control and Prevention | Guangdong Provincial Institution of Public Health | Jing Lu, Louis du Plessis, Liu Zhe, Jiufeng Sun, Sarah François, Huifang Lin, Moritz Kraemer, Jingju Peng, Qianlin Xiong, Runyu Yuan, Lilian Zeng, Pingping Zhou, Chuming Liang, Tao Liu, Wei Li, Juan Su, Huanying Zheng, Kang Min, Song Tie, Bo Peng, Shisong Fang, Wenzhe Su, Kuibiao Li, Ruilin Sun, Ru bai, Xi Tang, Minfeng Liang, Nuno Faria, Josh Quick, Andrew Rambaut, Verity Hill, Wenjun Ma, Nick Loman, Oliver Pybus, Changwen Ke |
| **EPI_ISL_413882** | hCoV-19/Guangdong/GD2020139-P0007/2020 | Asia / China / Guangdong | 2020-02-02 | Guangdong Provincial Institution of Public Health, Guangdong Provinical Center for Disease Control and Prevention | Guangdong Provincial Institution of Public Health | Jing Lu, Louis du Plessis, Liu Zhe, Jiufeng Sun, Sarah François, Huifang Lin, Moritz Kraemer, Jingju Peng, Qianlin Xiong, Runyu Yuan, Lilian Zeng, Pingping Zhou, Chuming Liang, Tao Liu, Wei Li, Juan Su, Huanying Zheng, Kang Min, Song Tie, Bo Peng, Shisong Fang, Wenzhe Su, Kuibiao Li, Ruilin Sun, Ru bai, Xi Tang, Minfeng Liang, Nuno Faria, Josh Quick, Andrew Rambaut, Verity Hill, Wenjun Ma, Nick Loman, Oliver Pybus, Changwen Ke |
| **EPI_ISL_413883** | hCoV-19/Guangdong/GD2020134-P0031/2020 | Asia / China / Guangdong | 2020-02-02 | Guangdong Provincial Institution of Public Health, Guangdong Provinical Center for Disease Control and Prevention | Guangdong Provincial Institution of Public Health | Jing Lu, Louis du Plessis, Liu Zhe, Jiufeng Sun, Sarah François, Huifang Lin, Moritz Kraemer, Jingju Peng, Qianlin Xiong, Runyu Yuan, Lilian Zeng, Pingping Zhou, Chuming Liang, Tao Liu, Wei Li, Juan Su, Huanying Zheng, Kang Min, Song Tie, Bo Peng, Shisong Fang, Wenzhe Su, Kuibiao Li, Ruilin Sun, Ru bai, Xi Tang, Minfeng Liang, Nuno Faria, Josh Quick, Andrew Rambaut, Verity Hill, Wenjun Ma, Nick Loman, Oliver Pybus, Changwen Ke |
| **EPI_ISL_413884** | hCoV-19/Guangdong/GD2020115-P0009/2020 | Asia / China / Guangdong | 2020-02-01 | Guangdong Provincial Institution of Public Health, Guangdong Provinical Center for Disease Control and Prevention | Guangdong Provincial Institution of Public Health | Jing Lu, Louis du Plessis, Liu Zhe, Jiufeng Sun, Sarah François, Huifang Lin, Moritz Kraemer, Jingju Peng, Qianlin Xiong, Runyu Yuan, Lilian Zeng, Pingping Zhou, Chuming Liang, Tao Liu, Wei Li, Juan Su, Huanying Zheng, Kang Min, Song Tie, Bo Peng, Shisong Fang, Wenzhe Su, Kuibiao Li, Ruilin Sun, Ru bai, Xi Tang, Minfeng Liang, Nuno Faria, Josh Quick, Andrew Rambaut, Verity Hill, Wenjun Ma, Nick Loman, Oliver Pybus, Changwen Ke |
| **EPI_ISL_413888** | hCoV-19/Guangdong/2020XN4239-P0034/2020 | Asia / China / Guangdong | 2020-01-30 | Guangdong Provincial Institution of Public Health, Guangdong Provinical Center for Disease Control and Prevention | Guangdong Provincial Institution of Public Health | Jing Lu, Louis du Plessis, Liu Zhe, Jiufeng Sun, Sarah François, Huifang Lin, Moritz Kraemer, Jingju Peng, Qianlin Xiong, Runyu Yuan, Lilian Zeng, Pingping Zhou, Chuming Liang, Tao Liu, Wei Li, Juan Su, Huanying Zheng, Kang Min, Song Tie, Bo Peng, Shisong Fang, Wenzhe Su, Kuibiao Li, Ruilin Sun, Ru bai, Xi Tang, Minfeng Liang, Nuno Faria, Josh Quick, Andrew Rambaut, Verity Hill, Wenjun Ma, Nick Loman, Oliver Pybus, Changwen Ke |
| **EPI_ISL_413889** | hCoV-19/Guangdong/2020XN4291-P0038/2020 | Asia / China / Guangdong | 2020-01-30 | Guangdong Provincial Institution of Public Health, Guangdong Provinical Center | Guangdong Provincial Institution of Public Health | Jing Lu, Louis du Plessis, Liu Zhe, Jiufeng Sun, Sarah François, Huifang Lin, Moritz Kraemer, Jingju Peng, Qianlin Xiong, Runyu Yuan, Lilian Zeng, Pingping Zhou, Chuming Liang, Tao Liu, Wei Li, Juan Su, Huanying Zheng, Kang Min, Song Tie, Bo Peng, Shisong Fang, Wenzhe Su, Kuibiao |

| | | | | | | |
|---|---|---|---|---|---|---|
| | | | | for Disease Control and Prevention | | Li, Ruilin Sun, Ru bai, Xi Tang, Minfeng Liang, Nuno Faria, Josh Quick, Andrew Rambaut, Verity Hill, Wenjun Ma, Nick Loman, Oliver Pybus, Changwen Ke |
